# Synthetic genome modules designed for programmable silencing of functions and chromosomes

**DOI:** 10.1101/2024.03.22.586311

**Authors:** Xinyu Lu, William M Shaw, Anima Sutradhar, Giovanni Stracquadanio, Tom Ellis

## Abstract

Unlike in bacteria, eukaryotes rarely cluster sets of genes in their genomes according to function, instead having most genes spread randomly across different chromosomes and loci. However, with the advent of genome engineering, synthetic co-location of genes that together encode a cell function has now become possible. Here, using *Saccharomyces cerevisiae* we demonstrate the feasibility of reorganising a set of yeast genes encoding a cell function, tryptophan biosynthesis, into a *synthetic genome module* by deleting these genes and their regulatory elements from their native genomic loci while in parallel reconstructing them into gene cluster format by synthetic DNA assembly. As part of synthetic module design, loxPsym sequences recognised by Cre recombinase are placed between all module genes, and we leverage these for a novel master regulation system we call dCreSIR. Using dCreSIR we externally control silencing of synthetic modules by targeted binding of chromatin recruiters to loxPsym sites and this leads to inhibition of local transcription. We further show that dCreSIR can go beyond modules and be used to specifically downregulate expression across an entire synthetic yeast chromosome containing loxPsym sites. Together, our work offers insights into yeast genome organisation and establishes new principles and tools for the future design and construction of modular synthetic yeast genomes.

## Introduction

Synthetic genomics is a sub-field of synthetic biology that is dedicated to the design, construction and manipulation of artificial genomes by using synthetic DNA at scale. While considerable progress has been achieved in synthesising prokaryotic genomes^1–5^ and eukaryotic genomes^6–10^, the field has not yet advanced to fully *de novo* design and the construction of whole genomes from modular DNA parts^11^. The Synthetic Yeast Genome Project (Sc2.0) stands out by incorporating unique design features and extensive sequence modifications into its synthetic chromosomes^8^; however, it is still fundamentally guided by the gene layout and structure of the native yeast genome. The complexity of genome content and in particular the organisational rules of chromosomes in yeast is as yet a largely unexplored question in synthetic genomics but could be the key to unlocking a future modular synthetic yeast genome^11^.

Whether the genes of a genome can be rearranged into modules, for example based on function, was first experimentally explored in the development of the synthetic minimal *Mycoplasma* genome JCVI-syn3.0^4^. In a design variant for this synthetic genome, approximately 12% of its genome was reorganised into modular segments, with each gene grouped into a section according to cellular function^4^. This strategy, termed “defragmentation”, is a concept analogous to optimizing a computer hard drive so that all files for a program are located together for more efficient control. In the context of yeast, genome defragmentation involves a strategic reorganisation of genes that are natively scattered throughout the 16 chromosomes, bringing these together into distinct, functional modules. Previous studies have shown that genes encoding the glycolysis pathway and the tRNA-encoding genes can be removed from their native genomic loci and relocated as functional and transferable units either at a single genomic locus or on a neochromosome^12–14^. These radical genome modifications did not significantly impact cell growth, pointing to the ability of the yeast genome to tolerate major changes in gene content and arrangement. However, despite these examples of successful synthetic gene clustering, the scope and extent of defragmentation possible at the genome scale in yeast remains uncertain.

In eukaryotes, particularly in plants and fungi, natural gene clusters encoding secondary metabolism are found to evolve in telomere-proximal heterochromatin for potential co-regulation^15^. However, such gene clustering is less common in the budding yeast genome. The *GAL* gene cluster, encoding the galactose catabolism pathway in yeast, has been shown to locate three *GAL* genes adjacently to help synchronize their expression and thus minimising the accumulation the cytotoxic intermediate galactose-1-P^16^. Similarly, the six-gene *DAL* cluster, encoding allantoin degradation pathway, exhibits evolutionary pressures that have driven such genes to cluster at telomeric region for epigenetic regulation^17^. Despite the advantages observed in these few cases when the expression of the genes in a specific metabolic pathway needs to stay constant or be co-regulated, the broader benefits of gene clustering to other metabolic functions in yeast remain to be explored. Investigating the defragmentation of genes encoding conditionally essential metabolic pathways, such as those in the amino acid biosynthesis pathways, offers a compelling case for improving our understanding of the pathway robustness under both selective and non-selective conditions, and may also provide strains that can enable industrial applications.

Introducing functional modularity into eukaryotic genomes also opens the possibility of engineering control over entire module functions via long range regulation methods, where whole sets of genes can be silenced by programmable epigenetic changes^18^. In *S. cerevisiae* loci-specific, natural epigenetic regulation is already seen in several places in the genome^19–22^. For example, at the telomeric region, epigenetic silencing is mediated by multiple proteins, among which the SIR (Silent Information Regulator) proteins are the key functional elements^23,24^. The SIR proteins (Sir2, Sir3 and Sir4) assemble into a complex in 1:1:1 stoichiometry, facilitating the establishment and spreading of heterochromatin by enzymatically modifying histones within the telomeric regions^25,26^. Manipulation of chromatin and epigenetic states in yeast cells using synthetic biology tools has already provided new ways to uncover relationships between chromatin modifications and gene regulation, and to achieve programmable epigenetic control of single genes^27^. In a future modular yeast genome, such customised synthetic epigenetic systems could be used to achieve targeted silencing of modules depending on their function.

Here, we report the design, construction, and controlled regulation of a synthetic genome module for yeast that encodes the function of tryptophan biosynthesis. We use CRISPR genome engineering to delete the five *TRP* genes and their natural regulatory elements from their native genomic loci, and then reintegrate these genes together at a single gene locus as a synthetic *TRP* module. We then characterised how genomic location effects *TRP* module function by integrating it into various genomic loci to determine any context dependencies under selective and non-selective conditions. Based on the module design, we designed and optimised a new master regulation system termed dCreSIR that can efficiently control multi-gene expression within the synthetic module. We then go on to demonstrate that this system can even scale to widespread repression across an entire yeast synthetic chromosome and discuss the potential of applying this for future studies controlling other modular genetic systems.

## Results

### Design and construction of the synthetic *TRP* module

To show the proof of concept of defragmenting a yeast genome, we started by building the native set of genes involved in tryptophan biosynthesis pathway into a synthetic functional module. The tryptophan biosynthesis pathway in yeast proceeds in 5 steps from chorismate to L-tryptophan and requires five genes, namely *TRP1*, *TRP2*, *TRP3*, *TRP4* and *TRP5*, which are essential or non-essential depending on the presence of tryptophan and aromatic amino acids in growth media (**Figure 1A**). As a test case for our ‘learn-by-building’ approach, we proceeded by first deleting the coding sequences and flanking regulatory DNA of each of the five *TRP* genes from their native genomic loci, and then reassembled all five genes together in a modular format, integrating this at the *URA3* locus. In this first version, each gene was assembled still flanked by its native promoters and 5’ and 3’ untranslated regions (UTRs) (**Figure 1B**).

**Figure 1.**
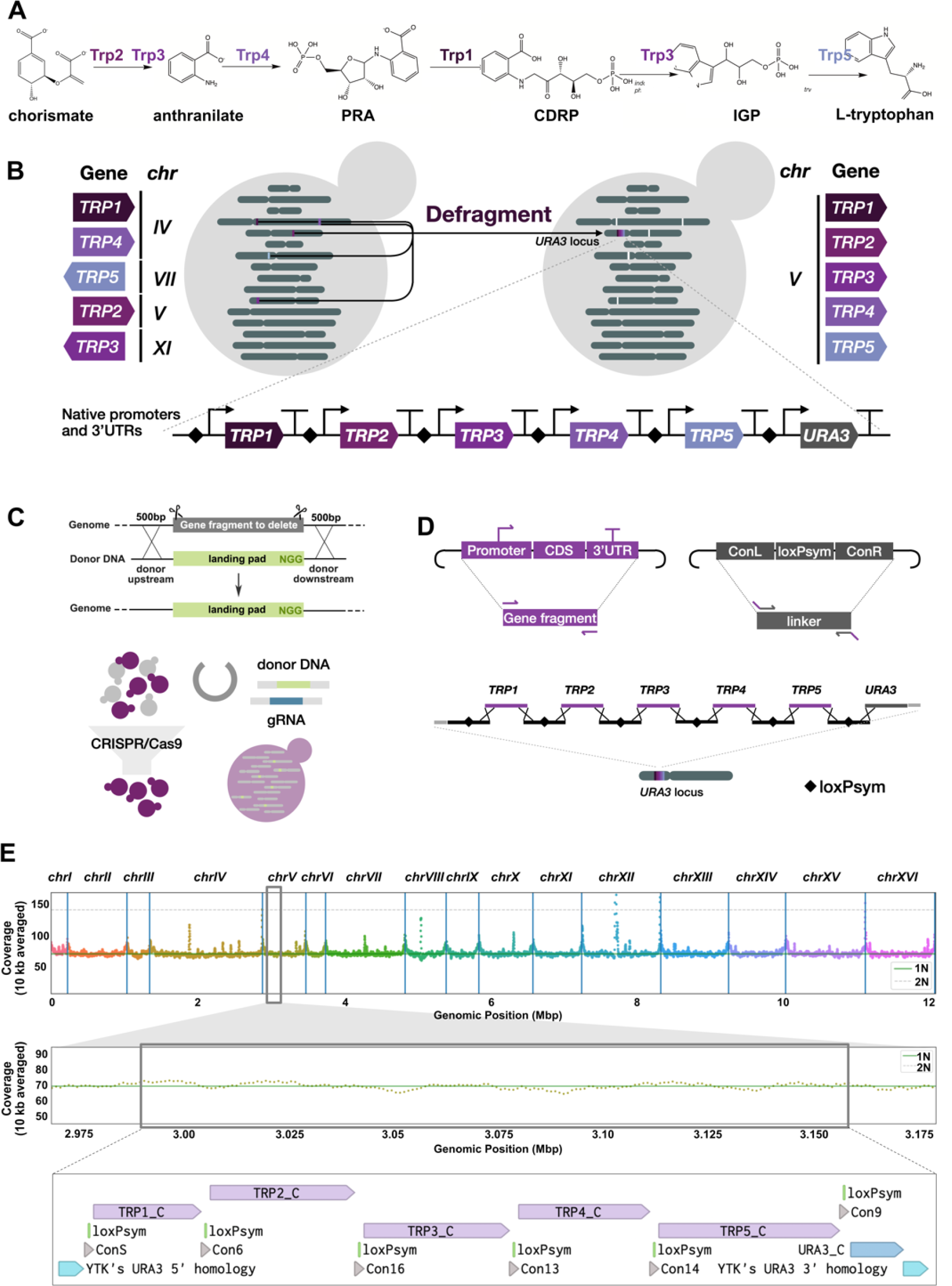
Construction of the synthetic *TRP* module. (A) Metabolic pathway of L-tryptophan biosynthesis from chorismate in *S. cerevisiae*. Intermediate metabolites *N*-(5-phospho-D-ribosyl)-anthranilate, 1-(o-carboxyphenylamino)-1’-deoxyribulose-5’-phosphate, and indole-3-glycerol-phosphate are shown as their abbreviations “PRA”, “CDRP” and “IGP”, respectively. (B) Schematic overview of defragmentation of *TRP* genes into a synthetic module. The five *TRP* genes involved in tryptophan biosynthesis, *TRP1*, *TRP2*, *TRP3*, *TRP4* and *TRP5*, were deleted from their native genomic loci and relocated together at the *URA3* locus. (C) Schematic of gene deletion via CRISPR/Cas9 editing and yeast homology directed repair (HDR)-based integration. Each deleted gene is replaced with a 23 bp landing pad containing a unique CRISPR/Cas9 target site. The CRISPR/Cas9 plasmids containing *URA3* marker is removed by growing cells in YPD overnight and counter-selecting with 5-FOA after each gene deletion round. (D) Schematic of process to generate synthetic *TRP* module via linearisation of DNA from pre-assembled entry-level plasmids and assembly into a module by homologous recombination in yeast. *TRP* gene cassettes were assembled by inserting wildtype genes with 1 kb upstream and 0.5 kb downstream sequences into vector pYTK001 using Gibson assembly. Linker plasmids were constructed by inserting a loxPsym sequence into synthetic connectors from the yeast Moclo Toolkit by PCR, phosphorylation, and ligation. (E) Top: read coverage of Illumina sequencing over the whole genome of strain yXL086, in which *TRP* genes are relocated to the *URA3* locus; Middle: a zoom-in read coverage of Illumina sequencing across the synthetic *TRP* module; Bottom: in silico design of the synthetic *TRP* module.

Deletion of *TRP* genes was accomplished through a markerless CRISPR/Cas9 editing method that can remove a gene or region of a chromosome and leave behind a minimal scar (**Figure 1C**). To ease future engineering of the sites left behind by gene deletion, in our design we substituted each deleted sequences with an individual 23 bp “landing pad” that encodes a unique CRISPR/Cas9 target sequence^28^. If restoration of local gene expression is needed, the deleted sequences could be reintroduced by targeting the designed landing pad with CRISPR/Cas9.

Prior to transformation into the recipient strain, the gene cassettes and the linkers were first constructed into individual entry-level plasmids using the standardised plasmids from yeast MoClo Toolkit^29^ (**Figure 1D**). In this context, gene cassettes contain the coding sequences of the *TRP* genes and their native flanking regulatory elements. The synthetic linkers (∼200 bp) connecting the gene cassettes were adapted from the assembly connectors used in YTK cloning, but specially designed to embed a 34 bp loxPsym sequence in the centre to allow for Cre mediated recombination within synthetic modules, similar to the SCRaMbLE system used in the Sc2.0 project^8^. Five *TRP* gene cassettes, six linkers, and a *URA3* selectable marker were linearised from the assembled plasmids and then integrated as a synthetic module (∼15 kb) at the *URA3* locus using in-yeast assembly to link them together by homology-dependent recombination. Genomic changes were confirmed by junction PCR to identify successful gene deletion and to show correct cluster assembly (**Figure S1**).

Three out of the eight tested colonies revealed full assembly of the complete 15 kb *TRP* cluster at the correct genome loci. One of these clones (yXL086) was selected for further examination and shown by whole genome sequencing to have had clean deletion of the 5 native *TRP* genes, successful assembly of the synthetic *TRP* module at the *URA3* locus, and a consistent level of read coverage across all other regions of the genome (**Figure 1E**). A summary of the strains and plasmids generated through the process of *TRP* gene deletion and *TRP* cluster assembly is shown in **Table S1** and **Table S2**, respectively. The synthetic linker plasmids are listed in **Table S3**.

### Defragmentation of *TRP* genes to the *URA3* locus caused no defects on cell fitness or pathway function

Removal of genes from their native loci risks disrupting the local sequence context, and therefore affecting gene expression of neighbouring genes left behind at the deletion site, potentially causing fitness defects^30^. In addition, changes to the local sequence context for relocated genes and their regulatory sequences may alter their expression when relocated into a module at a new genomic locus^31,32^. In the case of the *TRP* module, this would yield inefficient tryptophan biosynthesis. To assess the function of the synthetic *TRP* module, we inserted a pigment synthesis pathway (the 5 gene violacein biosynthesis pathway) that utilises tryptophan as a precursor into the genome to give a visual output of tryptophan availability (**Figure 2A**). As well as integrating this pathway into the *HO* locus of yXL086, we also generated two control strains, yXL085 and yXL094, where the violacein biosynthesis pathway genes were similarly integrated into a strain with all five *TRP* genes absent and into a BY4741 wildtype (WT) strain, respectively. We then performed growth spot assays on YPD, SC and SC-Trp agar media to assess the relative fitness and tryptophan pathway function of the *TRP* cluster-containing strain vs these two controls (**Figure 2C** and **Figure S2A**).

**Figure 2.**
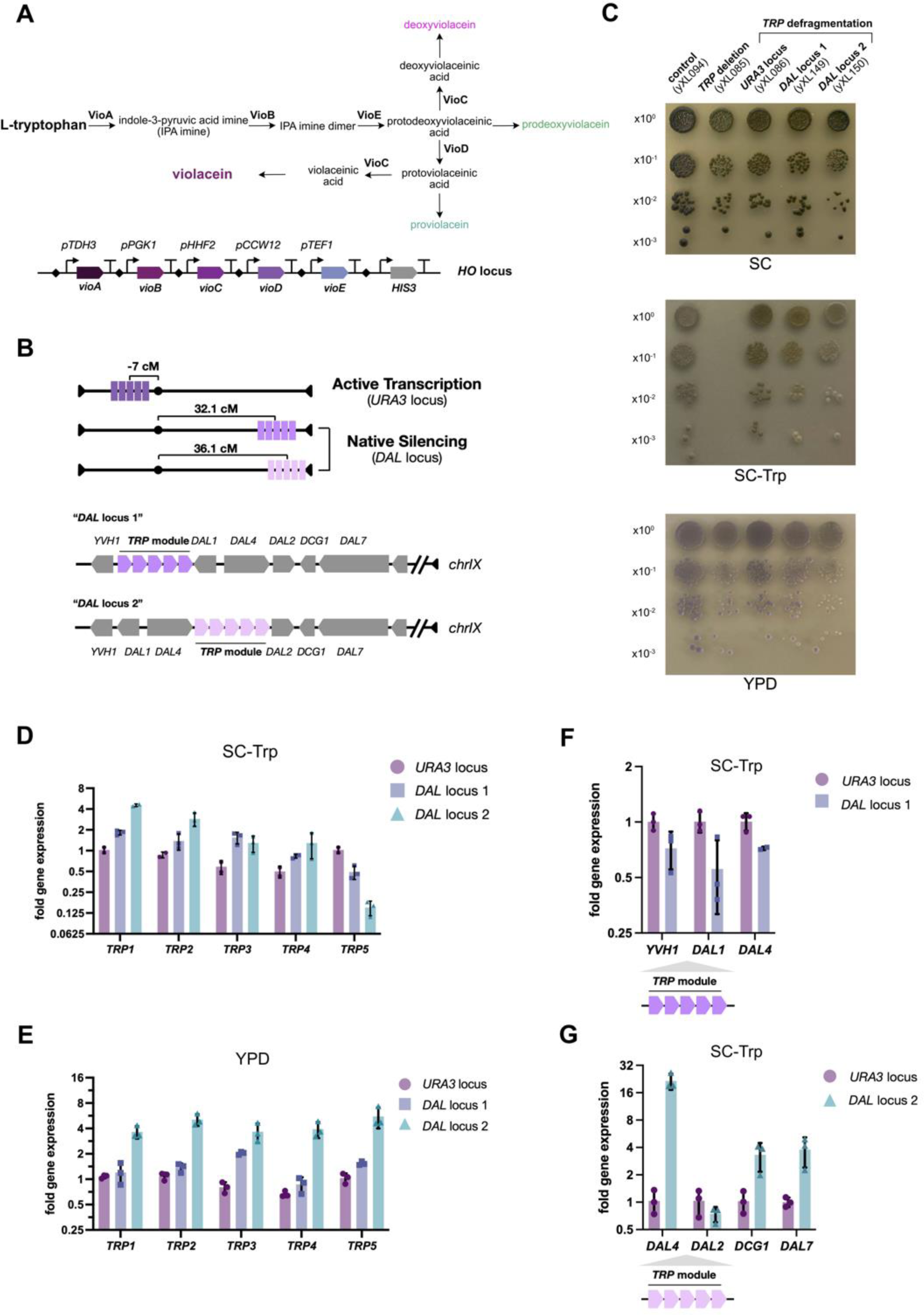
Functional characterisation of relocated synthetic *TRP* modules. (A) Illustration of the chromosomal integration of the violacein biosynthesis pathway for visualising tryptophan biosynthesis. L-tryptophan is utilised as a precursor for violacein synthesis. The *Vio* cluster, consisting of 5 genes (*vioA*, *vioB*, *vioC*, *vioD* and *vioE*) all under the control of strong constitutive promoters, was assembled through Golden Gate assembly in *E. coli* and subsequently integrated at *HO* locus with a *HIS3* selectable marker via the yeast homologous recombination-based assembly. (B) Schematic of integration of the synthetic *TRP* modules at different genomic loci to determine the spatial effects on gene silencing. (C) Spot assays of the strain yXL094, yXL085, yXL086, yXL149 and yXL150 on SC-Trp, SC and YPD agar to assess the cellular fitness and tryptophan biosynthesis. Cultures normalised to OD_600_ = 1.0 were serially diluted and spotted from top to bottom. (D and E) Quantification of the transcripts level of genes within the synthetic *TRP* module integrated at the *URA3* locus, “*DAL* locus 1” and “*DAL* locus 2” respectively in (D) SC-Trp and (E) YPD media, using *ACT1* as the reference gene, n=3. (F and G) Quantification of the transcripts level of neighbouring genes flanking the synthetic *TRP* module integrated at (F) “*DAL* locus 1” and (G) “*DAL* locus 2” in SC-Trp medium. Individual data points are plotted as round dots (defragmented *TRP* cluster at the *URA3* locus), squares (defragmented *TRP* cluster at the “*DAL* locus 1”) and triangles (defragmented *TRP* cluster at the “*DAL* locus 2”). Means of fold change expression are denoted by bar height. Error bars represent standard deviation.

Control strain yXL085, with an inability to perform tryptophan biosynthesis did not grow on SC-Trp and grew visibly slower than the WT control in YPD and SC media (**Figure S2A**). The strain with the synthetic *TRP* module at the *URA3* locus, yXL086, exhibited improved growth in YPD, SC and SC-Trp media compared to yXL085, however it did not attain the maximum growth rate of the WT control (**Figure S2A**). While yXL086 did not show a substantial reduction in viability compared to yXL094, it exhibited slight differences in tryptophan synthesis, as indicated by the differences in pigment intensity from the violacein reporter under the tested conditions (**Figure 2C**).

We next used quantitative RT-PCR (qPCR) to determine if differences in tryptophan biosynthesis were due to the changes in the mRNA levels of the *TRP* genes, perhaps due to relocation of the genes in the genome or another sequence context dependent effect. We confirmed that in both rich and SC-Trp media, the relocated *TRP1*, *TRP2* and *TRP5* genes had very similar mRNA levels compared to the native genes at their native genomic loci, while *TRP3* and *TRP4* exhibited a small reduction in their mRNA levels (**Figure 2D** and **2E**). Trp3 and Trp4 participate in catalysing the initial two steps of tryptophan biosynthesis from chorismate. Trp3 also catalyses the conversion of 1-(2-carboxyphenylamino)-1-deoxy-D-ribulose 5-phosphate to indoleglycerol phosphate, both of which are intermediates in the tryptophan biosynthesis. The small but potentially significant changes in *TRP3* and *TRP4* transcription could be an explanation for a visible slight decrease in violacein pigmentation.

Overall, relocation of the five *TRP* genes into a synthetic module at the *URA3* locus had no significant impact on cell viability in rich and selective media, and did not significantly impair function, as abundant tryptophan biosynthesis is still evidenced by the ability of the strains to produce violacein. However, it is important to note that the fitness and function of yXL086 has not yet been determined in a larger variety of stresses and growth conditions, where issues may arise.

### Assessing the effects of gene clustering in varied genomic contexts

The relationship between genomic context and gene expression regulation remains incompletely understood even in yeast^33^ and is of particular interest when considering the design and construction of synthetic modules, chromosomes and genomes. Synthetic genome modules where multiple genes that together perform a single testable function are clustered in a movable unit, offer an exciting new tool to investigate this. With this in mind, we set out to assess the functionality of our synthetic *TRP* module when relocated to a different genomic context. For this we examined a subtelomeric site 9.4 kb from chromosome IX’s right telomere, known for heterochromatin-induced transcriptional repression, and for harbouring the largest metabolic gene cluster in yeast, the *DAL* cluster, where specific histone modifications are established to be present^17,34^. We wanted to explore whether integration of a module into typically repressed region of a chromosome, results in silencing of its function.

To test loci specific effects in this region, we re-integrated the synthetic *TRP* module into two different sites within the *DAL* cluster of strain yXL085 (**Figure 2B**). This was done using the same cluster assembly method previously employed for assembling the *TRP* module at the *URA3* locus and yielded two distinct strains, named as yXL149 and yXL150. In yXL149, the *TRP* module was integrated upstream of the *DAL1* gene from the *DAL* cluster. We named this integration site “*DAL* locus 1”. While in yXL150, the *TRP* module was inserted within the *DAL* cluster at “*DAL* locus 2”, flanked by the *DAL4* and *DAL2* genes.

To assess for native silencing of the synthetic *TRP* cluster in each position at *DAL* locus, we performed growth assays and cluster functionality tests on strains yXL149 and yXL150, comparing these to the results of wildtype cells and yXL085 and yXL086 (**Figure 2C** and **S2A**). These assays were carried out under both selective and non-selective conditions. Both yXL149 and yXL150 exhibited a moderate growth defect and reduced tryptophan biosynthesis compared to yXL094 and yXL086 when grown in YPD, SC and SC-Trp, with yXL150 showing more obvious deficiencies under all conditions tested (**Figure 2C, S2A** and **S2C**). These results suggest that the tryptophan pathway may be repressed when relocated to a native silencing locus. As slow growth phenotypes we observed might also be attributed to a stress response under insufficient tryptophan synthesis, we visually inspected these strains by microscopy. The microscopy images did not reveal any enlarged budding cells in either yXL149 or yXL150, when compared with the control strains (yXL094 and yXL085) (**Figure S2B**). However, within the population of yXL150, we observed instances of cell aggregation when tryptophan was absent (**Figure S2B**). Without further information, we speculate that cells may flocculate as a stress response due to sustained nutrient limitation.

We next used qPCR to assess transcript levels of the *TRP* modules genes and their flanking genes at the *DAL* locus, comparing the mRNA levels with those observed in control strain yXL094. The qPCR data revealed distinctive transcriptional profiles when yXL149 and yXL150 were grown in SC-Trp (**Figure 2D**). In yXL149, the expression of the *TRP* genes within the synthetic module and their neighbouring genes at “*DAL* locus 1” were similar to the control, with no mRNAs changing levels by more than 2-fold (**Figure 2D** and **2F**). However, under the same selective conditions the second *DAL*-integrated strain, yXL150, displayed substantial upregulation of *TRP1* and *TRP2*, normal levels of *TRP3* and *TRP4,* and significant reduction in *TRP5,* which is the closest *TRP* gene to the main *DAL* cluster silencing region (**Figure 2D**). The flanking genes of the *TRP* module at this “*DAL* locus 2”, namely *DAL4*, *DCG1* and *DAL7*, all exhibited significant increases in transcription, as shown in **Figure 2G**.

Alongside these observations in selective media, we also used qPCR to investigate module transcription in rich media (**Figure 2E**). In yXL149, *TRP3*, situated in the centre of the *TRP* module at “*DAL* locus 1”, exhibited a 2-fold upregulation but all other genes exhibited less change (**Figure 2E**). However, in yXL150, all five *TRP* genes in the module at “*DAL* locus 2” displayed significant upregulation, going up to a 5.5-fold increase (**Figure 2E**). The upregulation of *TRP* genes at this locus in rich media was inconsistent with the observation of reduced tryptophan production in saturated cultures (**Figure S2C**), and the molecular mechanisms for this coordinated increase in *TRP* gene mRNA levels remain unclear to us. It is possible that the insertion of a cluster of actively transcribed genes between *DAL2* and *DAL4* disrupts the continuity of the *DAL* cluster, changing the local distribution of Htz1-activated domains (HZADs)^17,34^. This perturbation may disrupt the original repressive chromatin state at *DAL* locus to trigger dysregulation of *TRP* genes and alter *DAL* gene expression.

### Engineering synthetic epigenetic control of the synthetic modules

To enable external control of the functions of synthetic genome modules, we next sought to develop a synthetic system that can give ‘master switch’ regulation over all genes in a module, without needing to alter the DNA sequence of the genes and their promoters. For this we turned to the long-range regulation afforded by epigenetic control and attempted two main strategies: (1) the directed recruitment of chromatin regulators to the synthetic module DNA to initiate and maintain stable chromatin states^27^; and, (2) the strategic tethering of module DNA into sub-nuclear regions associated with heterochromatin, such as the nuclear periphery, in order to leverage spatial gene regulation^35–37^.

### Chromatin-based co-regulation of genes within synthetic modules

To achieve chromatin-based regulation at synthetic modules, we focused on manipulating the chromatin states of the clustered genes via inducible synthetic chromatin regulators (CRs). Usually in yeast, targeted regulation of genes is achieved by either manipulating the DNA sequences of promoters^38–40^ or recruiting transcriptional repressors to the core promoter region using a specific DNA binding domain (DBD)^27,41–44^. However, the unique design of our synthetic modules enabled us to instead make use of the intergenic linkers between genes that was used for their construction. Within these genes we selected the 34 bp loxPsym site as a potential target site for CRs to bind. Our synthetic design universally places this site between all module genes making it a great target for a single regulator to simultaneously bind to multiple positions in a gene cluster. Initially, we considered an approach where the CRISPR-based regulator dCas9 was used to bind loxPsym sites via a loxPsym-targeting guide RNA. However, the site sequence lacks a suitable Protospacer Adjacent Motif (PAM) that can be recognised efficiently by dCas9 protein. We thus developed an alternative strategy, where we would develop a catalytically dead mutant of Cre recombinase to act as a DNA binding domain (DBD) instead of using dCas9. To enable this to be inducible we fused this to the estrogen binding domain (EBD), so that the protein is excluded from the yeast nucleus unless β-estradiol is given in the growth media. We named this recombination inactive Cre-EBD fusion as dCre (“dead Cre”). Ideally, dCre should be able to specifically bind to each loxPsym site in a module upon the induction with β-estradiol, without generating any cleavage or recombination.

To obtain a suitable dCre for our needs, we first validated the cleavage activity of previously reported Cre recombinase mutants using a GFP reporter system (**Figure S3A**). This system involves two successive steps of recombination, inversion and then excision between the two pairs of orthogonal loxP variant sites, as shown in **Figure S3B**. The occurrence of both recombination events generates the irreversible flipping of the *mGFPmut2* gene, resulting in stable GFP expression, thus serving as an indicator of the cleavage activity. Using this reporter, we then validated a group of Cre mutants that had been previously reported to be catalytically inactive for cleavage or recombination^45^. Unlike with the wild type (WT) and K86A mutant versions of Cre, no GFP fluorescence was detected when the mutants K201A, R173K and Y324F were used, confirming their inability to cleave and recombine DNA.

Among these 3 candidate mutants, we selected Cre K201A as the best DNA binding domain, as it was reported to have higher synapsis activity compared to Y324F and R173K^45^. Synapsis is a process of bringing together the Cre-bound loxP sites to form the tetrameric protein complex^45^. This process of dCre tetramer assembly could facilitate the looping and compaction of the 3D DNA structure at loxPsym sites with the structural change likely helping suppress gene expression of genes flanked by loxPsym sites.

We next constructed a set of modular synthetic loxPsym-binding CRs that we called ‘dCre-CRs’ with these composed of an N-terminal dCre-EBD fusion as the DBD, then a (GS5)6 linker polypeptide and then a CR domain (**Figure S4A**). These were expressed from cassettes with constitutive promoters that were integrated at the *LEU2* locus, to ensure single gene copy per cell and stable and consistent expression during cell propagation. Upon induction with β-estradiol, the expressed dCre-CR fusion proteins are translocated to the nucleus and specifically bind to the loxPsym sites embedded within the linkers flanking each gene (**Figure 3A**).

**Figure 3.**
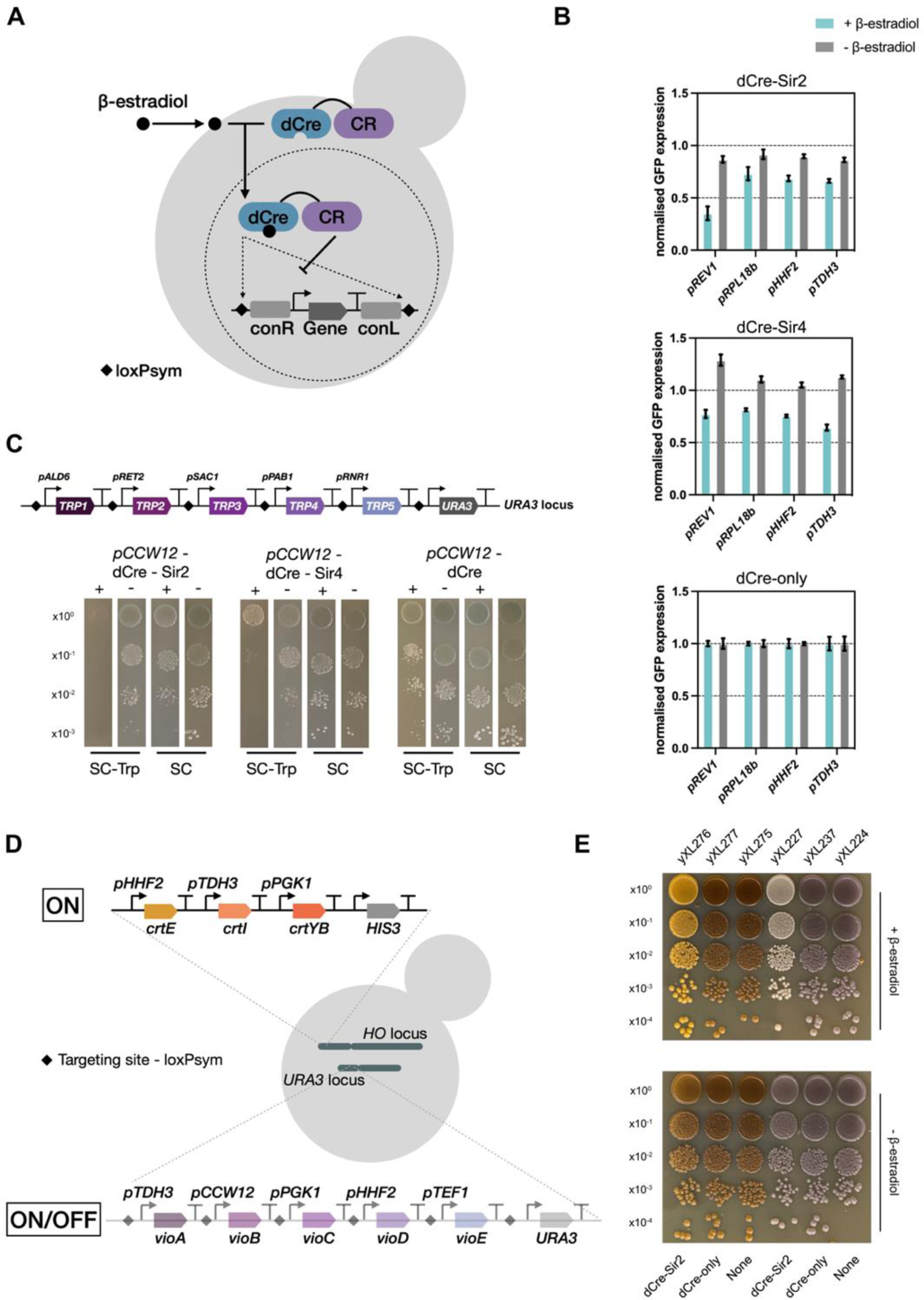
Targeted silencing of synthetic modules by dCre-CR silencing systems. (A) Schematic of targeted transcriptional regulation of genes flanking by linkers embedded with loxPsym sites. Each CR was fused to dCre and was individually recruited to loxPsym sites upon β-estradiol induction. (B) Characterisation of Sir protein silencing on the different synthetic constitutive promoters using a transcriptional reporter. (C) Spot assays showing the silencing effects on a synthetic *TRP* module when inducing the dCreSIR system under SC-Trp and SC conditions. Each *TRP* gene is regulated by a constitutive promoter selected from the yeast MoClo toolkit^29^. Images cropped to show comparisons were taken from the same plate incubated at 30 °C for 2 days. Full images were shown in Figure S6. (D) Schematic of strain with violacein biosynthesis module integrated at *URA3* locus, while simultaneously co-expressing a β-carotene synthesis pathway that is integrated at *HO* locus. (E) Spot assays showing targeted downregulation of the violacein synthesis in YPD medium when inducing the dCre-Sir4 silencing system. Plates were incubated at 30°C for 3 days.

Using this design, we investigated the silencing effects of 7 selected CRs (Sir2, Sir3, Sir4, Tup1, Mig1, Rph1, and Mxi1) using a synthetic transcriptional reporter (**Figure S4A**). The first three CRs, Sir2, Sir3, and Sir4 are essential components of the SIR protein complex, which is involved in silencing genes at mating type loci and telomeres^26^. Mig1, Tup1, and Mxi1 are well-characterised repressors frequently used for CRISPRi that can enhance transcriptional repression by influencing local histone modifications^41,43,46^. Lastly, Rph1 has been previously fused to a synthetic zinc finger (ZF) protein to generate a ZF-Rph1 fusion regulator previously shown to give targeted long-range repression activity using a triple gene fluorescent reporter^27^.

For the reporter, we designed a genome-integrated cassette where the expression of *sfGFP* is controlled by a synthetic constitutive promoter (∼700 bp) and a terminator (∼220 bp) selected from the yeast MoClo toolkit^29^, with these flanked by module linkers each with a loxPsym site in the centre. We generated a set of strains with this reporter, where each dCre-CR candidate is expressed by one of three constitutive promoters classified as strong (*pCCW12*), medium (*pALD6*) and weak (*pPSP2*).

We then used flow cytometry to measure the green fluorescence per cell of these strains with and without β-estradiol inducer in exponential and in stationary growth phases. Repression efficiency of the dCre-CRs was calculated by normalising the single cell GFP measurement to that of a control strain expressing dCre without any fusions (’dCre-only’).

In our first set of strains, the sfGFP reporter was expressed from the *HHF2* promoter. With this we found that the dCre-CR fusions downregulated GFP output with varying efficiency (**Figure S4B**). dCre-CRs expressed from medium or weak promoters showed negligible repression and only after overexpression of dCre-Sir2, dCre-Sir3, dCre-Sir4, and dCre-Tup1 from the *CCW12* promoter was notable GFP repression seen (around 30% decrease). Minimal repression was seen with dCre-Mig1,-Mxi1, and -Rph1 constructs (**Figure S4B**), which contrasts with many reported successes in fusing Mxi1 and Mig1 to dCas9, dCas12a, and ZF proteins and using them to repress transcription by targeting their binding to just upstream of a minimal promoter or at transcription start sites (TSSs)^27,41,43,44,46^. Interestingly, overexpression of dCre-Sir2, dCre-Sir3, dCre-Sir4, and dCre-Mig1 all reduced GFP expression by up to 56% when the assay was done in stationary phase (**Figure S4C**). It is important to note that as sfGFP has a long half-life in yeast, even perfect repression of its promoter would not lead to 100% reduction of GFP levels in cells in these assays, as once the promoter is fully repressed it still takes hours and many rounds of cell division to lose or degrade all sfGFP protein that was present in the cell before repression was started.

The repressive efficacy of specific CRs will likely differ depending on the sequence context of the local promoters in the region targeted for repression. To investigate if there are promoter-specific effects, we replaced the promoter for *sfGFP* in the reporter cassette with a set of synthetic constitutive promoters, each characterised by distinct strengths (*pREV1* < *pRPL18b* < *pTDH3,* see **Figure 3B** and **S4A**). Compared with dCre-only, overexpression of dCre-Sir2 fusions resulted in a consistent level of repression of the *sfGFP* gene with all three promoters, *pRPL18b*, *pTDH3* and *pHHF2*. Notably, dCre-Sir2 overexpression led to a 65% decrease in GFP expression from *pREV1,* indicating a potential improved silencing effect in the context of a gene target with weak promoter. Similarly, dCre-Sir4 overexpression resulted in consistent repression on all tested promoters but unlike dCre-Sir2, it showed no variation in repressive efficiency related to target promoter strength.

To verify that the changes in reporter expression were not due to non-specific effects of dCre-Sir protein overexpression, we rebuilt the reporter construct but now with linkers lacking loxPsym sites and we re-examined GFP expression following a 6-hour induction of β-estradiol. Overexpression of dCre-Sir2 exhibited a slight repression, whereas the overexpression of dCre-Sir4 showed negligible repression (**Figure S4D**), suggesting minimal off-target repression.

We next examined the silencing effects on larger multigene constructs, using biosynthetic clusters as an example. We introduced our dCre-CR silencing system into yeast strains with synthetic modules integrated into their genomes that express the β-carotene biosynthesis pathway (*Crt* cluster) or the violacein biosynthesis pathway (*Vio* cluster). Consistent with the findings from the GFP reporter assays, overexpression of dCre-Sir2, dCre-Sir4 and dCre-Tup1 in these yeast strains resulted in a noticeable reduction in visible amounts of β-carotene and violacein biosynthesis, compared to non-induced strains and in the controls with only dCre expression (**Figure S5A**). Growth assays during these experiments suggested that the overexpression of dCre-Sir2 and dCre-Sir4 fusions in these strains did not cause any notable growth defects (**Figure S5B**). However, cells overexpressing dCre-Tup1 displayed flocculation in liquid cultures (**Figure S5B**), indicating fitness defects possibly arising from the perturbation of the global transcriptional regulation mediated by the Tup1-Cyc8 complex^47^.

We next applied the dCre-CR silencing system to downregulate an engineered version of our *TRP* module that now has each *TRP* gene expressed from a weak constitutive promoter (**Figure 3C**). We examined the efficacy of silencing this synthetic *TRP* module through spot assays to test the growth of colonies of cells on selective and non-selective media. Following 48 hours of β-estradiol induction, high level expression of dCre-Sir2 and dCre-Sir4 in this strain showed the strongest growth inhibition on SC-Trp selective media, suggesting targeted repression of *TRP* gene expression (**Figure 3C** and **S6**). A slight recovery of growth after an extra day of incubation was seen and could be due to decreased inducer (e.g. due to degradation) weakening the silencing effect over time or might be explained by cells adapting over time in order to survive in a tryptophan-deprived environment.

We found that overexpression of dCre-Tup1 and dCre-Mig1 in this strain also resulted in growth inhibition upon induction (**Figure S6**), but these 2 regulators were not used in subsequent experiments due to Tup1-induced flocculation and the inconsistency of downregulation by dCre-Mig1. Overexpressing dCre-Rph1 and dCre-Mxi1 significantly impaired cell growth, but the molecular mechanism for this is unclear to us. Notably, overexpressing and recruiting dCre without CR fusion also retarded growth when tryptophan was absent in the media. This could be due to the tetramerisation of dCre, which can theoretically loop synthetic module DNA encoded and presumably inhibit *TRP* gene transcription.

From above results, we conclude that the dCre-Sir2 and dCre-Sir4 silencing systems can inducibly and efficiently repress both genomically integrated heterologous pathways and an endogenous tryptophan biosynthesis pathway when these are formatted as synthetic modules with intergenic loxPsym sites. As dCre-Sir2 and dCre-Sir4 displayed the highest efficiency on gene repression among the tested CRs, we then decided to use these in the further experiments described below. As a group, we call these general repressors as **dCreSIR** proteins.

### Using dCreSIR silencing for selective pathway control in dual pigment biosynthesis

Yeast cells offer an adaptable platform for metabolic engineering and could potentially be built to host many diverse heterologous pathways as distinct functional modules within a single strain. Such a strain design would ideally contain a genetic control system that allows targeted switching on or off of selected pathways in order direct metabolic fluxes while minimising metabolic burden and resource use. Using the dCreSIR silencing system developed above, we set out to demonstrate the feasibility of modular pathway control in a yeast strain where both violacein and carotene biosynthesis pathways are genomically integrated (see schematic in **Figure 3D**). We constructed a dual pigment-producing yeast strain by first integrating the violacein biosynthesis pathway (*Vio* cluster) at the *URA3* locus of the base strain yXL224. We used synthetic genome module design, so that all 5 *vio* genes are flanked by intergenic linker sequences with loxPsym sites.

We next generated strain yXL237 by inserting a dCre-Sir4 expression cassette into the *LEU2* locus and a control strain yXL245 with a dCre cassette integrated at the same locus. We then integrated a synthetic carotenoid biosynthesis cluster (*Crt* cluster) into the *HO* locus of the three above-described strains to generate three further strains (yXL275, yXL276, and yXL277 respectively) that each now contain two biosynthetic clusters producing pigments (violet and orange). Importantly, the *Crt* cluster was built to lack loxPsym sites and so should not be a target for dCre and dCreSIR proteins.

Induction with β-estradiol led to targeted repression by dCreSIR of the violacein biosynthesis pathway in strain yXL276 as we had anticipated (**Figure 3E**). This was confirmed by the change in colony pigmentation from brown (violet + orange) to just orange in the colonies that grew. This colour change indicates reduced violacein biosynthesis in these cells while β-carotene synthesis is not visibly affected. Importantly, the control strains yXL275 and yXL277 exhibited no notable differences in colony pigmentation during the same tests, validating the specificity of the pathway repression mediated by dCre-Sir4. Further analysis of mRNA levels in these strains by qPCR confirmed an approximate 2-fold decrease in transcript abundance for the 5 *vio* genes upon induction of dCre-Sir4 silencing, while the expression of the neighbouring genes (*URA3* and *GEA1*) flanking the *Vio* cluster and 3 untargeted *crt* genes was not affected (**Figure S7**). However, we note that there was also an approximate 2-fold decrease in transcription of one gene (*vioC*) when overexpressing dCre with no Sir fusion. The central location of this gene within the *Crt* cluster again suggests that dCre tetramerisation, which theoretically causes DNA looping could be behind the inhibition of transcription of this gene.

### Improving silencing by swapping DNA binding domains and redesigning binding sites

To improve the silencing efficiency, we engineered a fusion of Sir2 and Sir4 together with dCre, aiming to increase recruitment of CRs in a single event. This design was inspired by the native Sir2/Sir4 complex assembly, which forms a 1:1 ratio and is known to mediate silencing at mating-type loci and telomeric regions^25,26^. However, we did not observe significant improvement in gene repression by this Sir4-Sir2 fusion and overexpression of dCre-Sir4-Sir2 fusion resulted in a significant growth defect (**Figure S8**). Instead, to further improve targeting and CR recruitment, we changed the DNA binding domain (DBD) to that of the classic transcriptional repressor TetR, and we increased the number of binding sites in the intergenic regions by adding seven TetO repeats into each of our module synthetic linker sequences. We first quantified the silencing effects of the TetR-CR fusions using a fluorescent reporter (**Figure S9A**) where the reporter gene *sfGFP* is expressed by the *pHHF2* promoter, as in the design of the reporter for characterising dCre-CR silencing. TetR-CR fusions were expressed from the *LEU2* locus, either from the strong promoter *pCCW12* or from the medium strength promoter *pALD6* (**Figure S9B**). As shown in **Figure S9C**, the presence of anhydrotetracycline (aTc) prevents TetR-CR fusion binding to TetO repeats, allowing high GFP production, whereas aTc absence leads to GFP repression by enabling the recruitment of TetR-CR fusions to the locus. By comparing the single-cell green fluorescence intensities between samples induced with aTc for 6 hours and uninduced controls, we confirmed that TetR-CR fusions enhanced the repression efficiency across all tested CRs. Notably, expressing TetR-Sir4 expressed from *pALD6* and TetR-Sir2 expressed from *pCCW12* resulted in the most significant repression, with fold change expression reductions of 12.04 and 5.77, respectively (**Figure S9D**).

We next examined the silencing effects of the TetR-Sir2 and TetR-Sir4 fusions on a re-engineered β-carotene biosynthesis pathway. This *Crt* cluster was modified to include 7xTetO sequences in the intergenic linkers flanking each gene cassette (**Figure 4A**). The TetR-Sir protein expression cassettes were integrated into this strain at the *LEU2* locus (**Figure 4B**). This inducible TetR-Sir silencing system consistently repressed the β-carotene synthesis without aTc, and reversibly activated the synthesis when TetR-Sir fusions are released upon aTc induction (**Figure 4C**). We observed TetR-Sir2 fusion successfully maintaining the β-carotene synthesis in the OFF-state in the absence of inducer. After 48 hours of induction, the β-carotene synthesis restored to levels comparable to control samples without TetR-CR integration (**Figure 4D** and **4E**). In this experiment, TetR-Sir4 recruitment did not achieve the same level of repression as TetR-Sir2. However, it still exhibited an 8.2-fold decrease in β-carotene levels compared with induced samples. It was also noted that the expression of just the TetR DBD without any CR fusions, did not influence β-carotene production.

**Figure 4.**
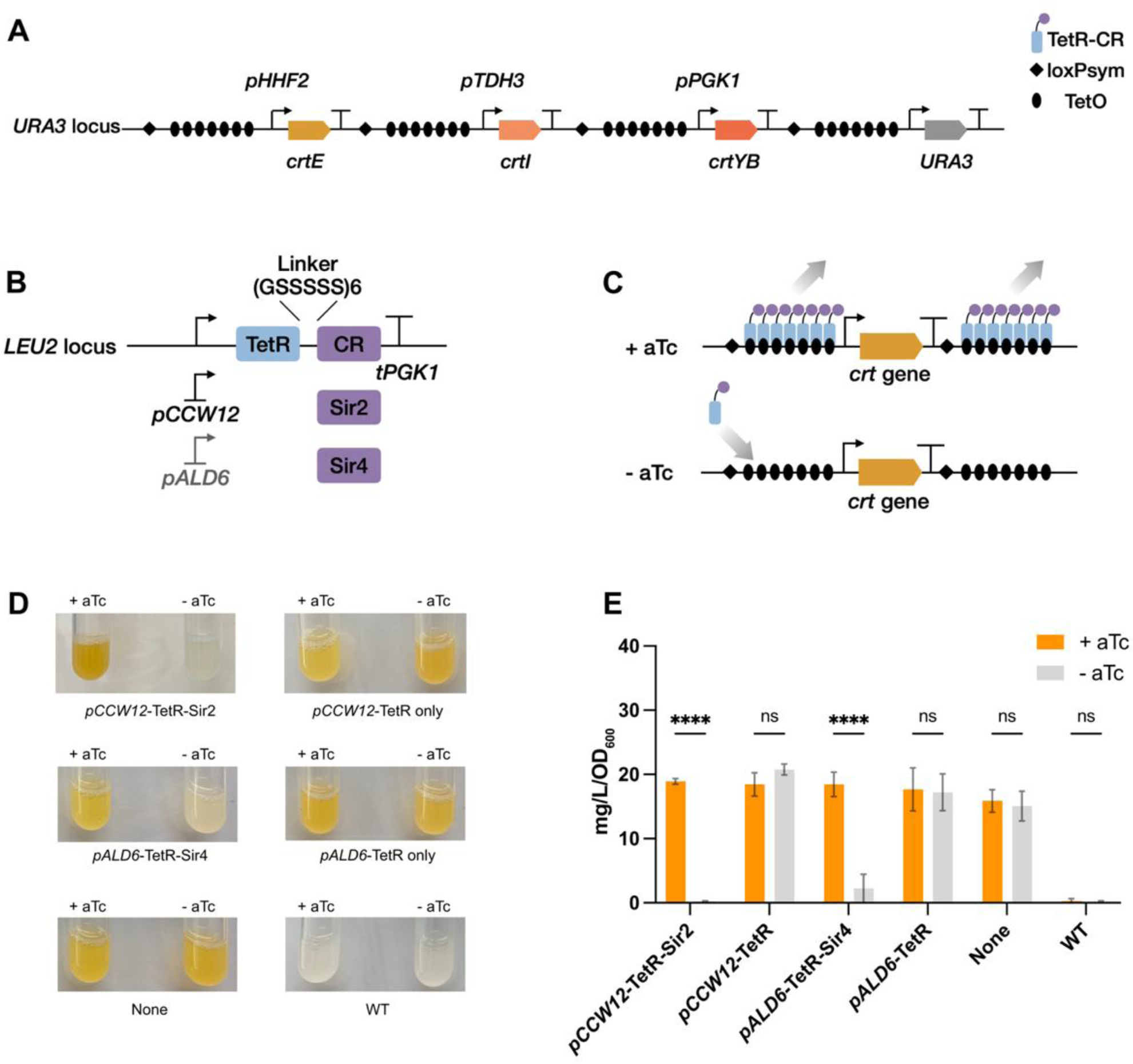
Improving targeted silencing by increasing local recruitment of SIR proteins. (A) Schematic of reformatting the synthetic *Crt* cluster by inserting seven TetO sites into each linker. (B) Schematic of the *LEU2* integrated TetR-Sir2 and TerR-Sir4 fusion protein expression cassettes. (C) Diagram showing the synthetic *tetO*-TetR-Sir protein regulation on individual *crt* gene in the presence and absence of 1 µM aTc. (D) β-carotene production in the strains overexpressing TetR-Sir2 and TetR-Sir4 in SC-Ura-Leu under induced and uninduced conditions for 2 days. (E) Quantification of β-carotene extracted from the 2 mL culture in SC-Ura-Leu after 48 hours in the presence and absence of 1 µM aTc. Experimental measurements are β-carotene concentrations determined by spectrophotometer at the absorbance of 453 nm in acetone, shown as the mean ± SD, n=2.

### Developing spatial control of synthetic modules by tethering to the nuclear periphery

An alternative strategy for synthetically modulating regulatory states of the synthetic modules is to physically tether them to native silencing foci. Programmable regulation by spatial redesign of the genome has been demonstrated in yeast cells and mammalian cells using CRISPR-Cas technologies and other DNA-protein binding systems^35–37,48–51^. However, the functional roles of these tethering approaches have led to mixed results. Nonetheless, anchoring to nuclear periphery is common in silencing of yeast telomeres and the mating type locus, with heterochromatin establishing proteins increased in local concentration at the nuclear periphery^51^. Following this concept, we investigated the feasibility of inducibly tethering synthetic genome modules to the nuclear periphery using dCre-anchor fusions.

We engineered synthetic tethers by fusing dCre to a set of periphery-associated proteins (Yif1, Nur1, Heh1, Mps3, Esc1, Yip1, Yku80), connecting them by a (GS_5_)_6_ linker, as illustrated in **Figure S10A**. We then assessed the repression effect of these tethers by visualising the production of pigment colour from a β-carotene biosynthesis pathway (*Crt* module) with intergenic loxPsym sites, that was integrated at the *URA3* locus. Our results revealed that only the dCre-Heh1 fusion gave visual repression of β-carotene synthesis (**Figure S10B**). However, the dCre-Heh1 tethering also resulted in an observable growth defect, potentially due to off-target effects or repression of essential genes flanking the *URA3* locus. In addition to repression seen with dCre-Heh1 overexpression, a small possible repression of β-carotene biosynthesis was also observed with dCre-Yif1 overexpression, when compared to control groups.

To further confirm the targeted silencing by dCre-Yif1 and dCre-Heh1 tethers, we overexpressed these two fusion proteins to assess their effects on repression of a synthetic *TRP* module with its genes expressed by weak constitutive promoters. After 22 hours of β-estradiol induction, strains expressing both dCre-Heh1 and dCre-Yif1 fusions exhibited approximately 3-fold reduction in OD_600_ compared to their uninduced counterparts in SC-Trp (**Figure S10C**). Notably, overexpression of dCre-Yif1 caused significant fitness defects as indicated by reduction in growth in the absence of β-estradiol induction (**Figure S10C**).

Based on the results above, we conclude that the dCre-Heh1 mediated synthetic tethering is a viable strategy for regulating synthetic modules. However, it’s worth noting that the effectiveness of inducible repression attained by dCre-Heh1 does not match that achieved by the dCreSIR or TetRSIR silencing systems. Moreover, the cellular fitness defects associated with this synthetic tethering method may limit its applications.

### Silencing an entire yeast chromosome with the dCreSIR system

We next asked whether our dCreSIR system could go beyond silencing of synthetic genome modules and extend to silencing of entire synthetic chromosomes. The most prominent design feature of the Sc2.0 genome is the placement of hundreds of loxPsym sites throughout each synthetic chromosome^8^, giving us the ideal opportunity to test whole chromosome silencing in yeast.

We introduced the dCre-Sir2 and dCre-Sir4 silencing systems into the *synXI* yeast strain generated by our group^52^. This strain contains a synthetic version of the 660 kb chromosome XI (called ‘*synXI’*) with loxPsym sites inserted into the 3’UTRs of almost all non-essential genes, and at other sites with key design modifications. In total there are 199 loxPsym sites in this chromosome with them distributed on average approximately every 3 kb^52^. In theory, in the presence of 1 µM β-estradiol, our dCreSIR regulators should bind to the loxPsym sites embedded in *synXI* and spread heterochromatic silencing across the entire chromosome, as illustrated in **Figure 6A**.

To study of the targeted gene silencing effects on *synXI* genes while preserving cell viability after silencing *synXI*, we introduced the dCreSIR constructs (and the dCre-Heh1 tethering construct too) into our haploid *synXI-*containing strain and then also into a heterozygous diploid strain made for this study. This diploid strain possesses two sets of chromosomes: one set from the *synXI* strain and the other from BY4742, and so contains 31 chromosomes including *chrXI* that are wildtype sequence and lack any loxPsym sites, and 1 chromosome (*synXI*) that is synthetic sequence and dense with loxPsym sites. Another group of control strains was also generated by integrating the selected dCre fusions into BY4742 yeast which contains no loxPsym sites.

We first did spot assays to characterise the silencing effects of our proteins on *synXI* upon the induction of 1 µM β-estradiol (**Figure 5B** and **S11**). The *synXI* haploid strain demonstrated complete growth inhibition when dCre-Sir2 was overexpressed, presumably because the only copy of the 200+ genes from this chromosome were being repressed. Overexpression of dCre-Sir4 and just dCre alone also significantly impacted growth when compared to the controls, while dCre-Heh1 overexpression also caused a noticeable, although less severe, growth inhibition (**Figure S11**). In the heterozygous diploid strain, where the cell has a non-synthetic version of the chromosome to fall back on, induction of dCre-Sir2 did not cause any fitness defects and only mild effects were seen when dCre-Sir4 and dCre-Heh1 were induced, perhaps due to non-specific binding of these two fusions to the wild type chromosomes.

**Figure 5.**
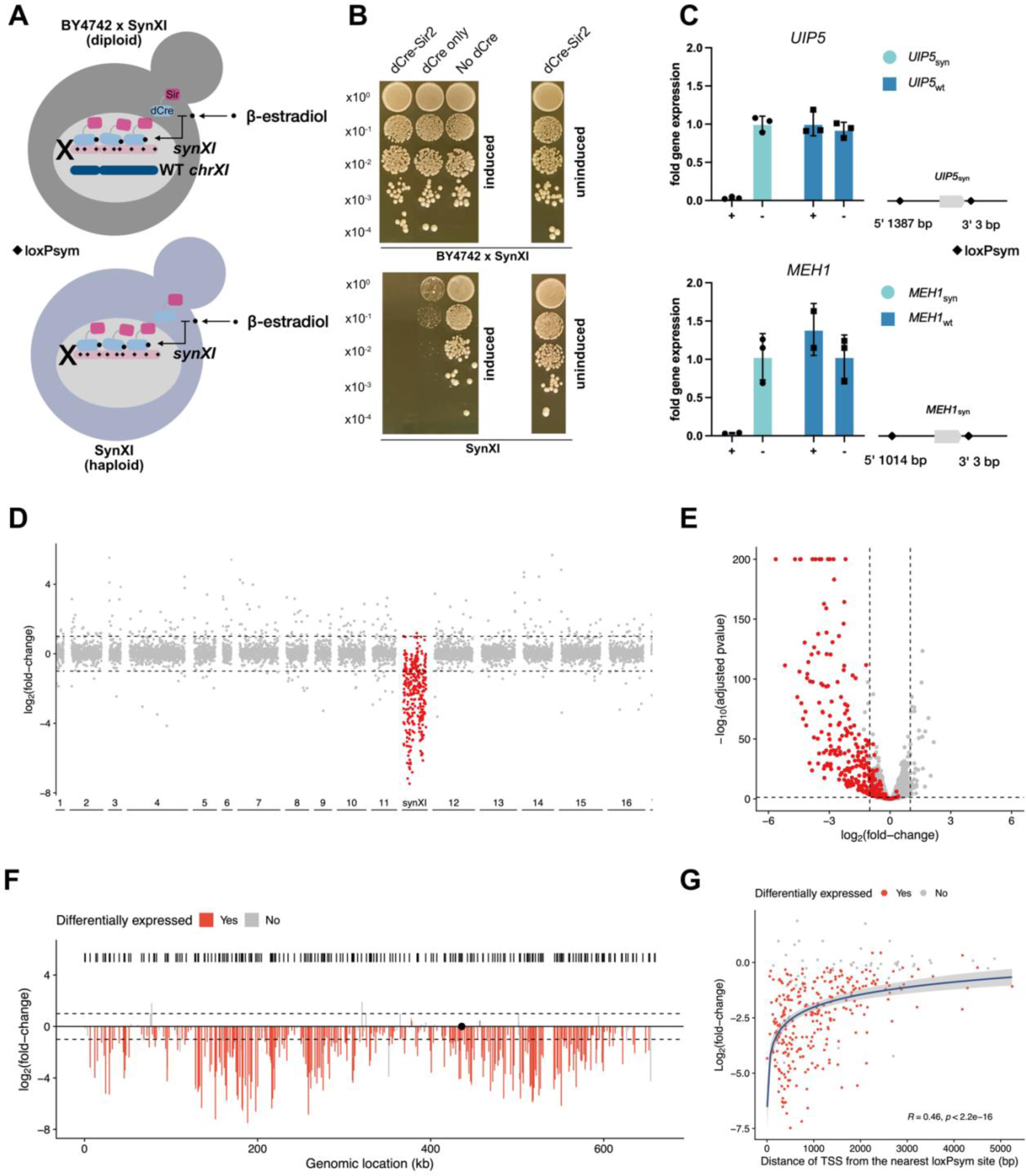
dCre-Sir2 mediated silencing of synthetic chromosome XI (*synXI*). (A) Schematic of the *synXI* silencing mediated by dCre-Sir2 system in a haploid strain and a heterologous diploid strain. In the presence of 1 µM β-estradiol, overexpressed dCre-Sir2 fusion binds to loxPsym sites across *synXI*, spreading heterochromatic silencing over the chromosome. (B) Spot assays showing cell viability of haploid strain *synXI* and heterologous diploid strain BY4742 x *synXI* overexpressing dCre-Sir2, respectively, upon induction with 1 µM β-estradiol. Cultures normalised to OD_600_ = 1.0 were serially diluted and spotted from top to bottom. Cells were incubated on YPD agar at 30°C for 3 days with and without 1 µM β-estradiol. Images were taken from the same plate, cropped, and reorganised to show comparisons. Full images, including all control groups, are shown in Figure S11. (C) Left: Transcripts of the synthetic and WT copies of two non-essential genes (*UIP5* and *MEH1*) on chromosome XI quantified by qPCR, using *ACT1* as reference. Experiments were performed in biological triplicate under induced and uninduced conditions. Individual data points of genes on *synXI* are plotted as round dots, individual data points of genes on WT *chrXI* are square dots, mean averages are denoted by bar height and error bars represent standard deviation. Right: schematic showing loxPsym insertion sites at each genomic loci of tested genes. Genomic loci of the genes, strategic qPCR primer design and transcripts of other tested genes are shown in Figure S12. (D) Manhattan plot of differential gene expression on *synXI* and WT chromosomes in heterologous diploid BY4742 x *synXI* overexpressing dCre-Sir2 fusion, as determined by RNA-seq. Comparisons were conducted between dCre-Sir2 induced vs uninduced. Adjusted p-value cutoff was set at 0.05. Dashed line represents fold change threshold of 2 (log_2_ fold change = 1 or −1). Genes on *synXI* are shown as dots in red. Genes on WT chromosomes are shown as dots in grey with the numbers indicates each WT chromosome. (E) Volcano plot showing differential gene expression on *synXI* in heterologous diploid BY4742 x *synXI*, as determined by RNA-seq. Comparisons were conducted between samples dCre-Sir2 induced vs uninduced. X-axis represents log_2_ fold change in gene expression between groups. Y-axis shows log_10_ of the p-value from the statistical test, with threshold of 0.05. Dashed line represents fold change threshold of 2 (log_2_ fold change = 1 or −1) and the p-value threshold of 0.05. Genes on *synXI* are shown as dots in red. Genes on WT chromosomes are shown as dots in grey. (F) Bar plot showing differential gene expression across *synXI*. X-axis represents genomic location of genes. Y-axis represents log_2_ fold change. Locations of loxPsym sites are shown as the black barcode. Comparisons were conducted between samples dCre-Sir2 induced vs uninduced. Genes differentially expressed are marked in red. Genes not differentially expressed are grey. Dashed line represents fold change threshold of 2 (log_2_ fold change = 1 or −1). Centromere is labelled as a black dot. (G) Correlation of fold change expression of down-regulated genes on *synXI* with distance of the nearest loxPsym sites to the gene start codon. X-axis represents distance of nearest loxPsym site to the gene start codon. Y-axis represents the log_2_ fold change in gene expression. Comparisons were conducted between samples dCre-Sir2 induced vs uninduced. Genes differentially expressed are marked in red. Genes not differentially expressed are grey.

To determine whether observed growth defects upon dCre-Sir2 silencing was a consequence of *synXI* repression as we had intended, we performed qPCR to quantify the mRNA levels of 6 selected genes distributed across both *synXI* and its wildtype equivalent chromosome XI (*chrXI*) (**Figure S12A**). We utilized the unique PCRtags sequences in the Sc2.0 chromosome protein coding sequences for discerning the transcripts from *synXI* from those from *chrXI* (**Figure S12B**). We found that the induction of the dCre-Sir2 silencing system led to varied levels of transcription changes in the selected six *synXI* genes when compared to uninduced samples and to *chrXI* gene expression changes (**Figure 5C** and **S12C**). Four of the *synXI* genes assayed showed insignificant downregulation relative to uninduced samples and their *chrXI* counterparts (**Figure S12C**). However, we observed almost complete transcriptional shutdown for two of the representative genes; *UIP5*_syn_ and *MEH1*_syn_, with approximately 20-fold and 27-fold decreases in their transcript levels, respectively, when compared with uninduced samples (**Figure 5C**). Expression of their *chrXI* copies was not affected either (**Figure 5C**). *UIP5*_syn_ and *MEH1*_syn_ are both non-essential genes so their nearest loxPsym sites are at their 3’UTR (**Figure 5C**). Coincidently, both these genes also have an additional loxPsym site designed for their neighbouring genes closely located upstream (**Figure 5C**). These qPCR results indicated that the efficiency of gene repression may be modulated by the proximity of adjacent loxPsym sites.

To assess the impact of dCreSIR mediated silencing on the transcriptome, we performed RNA-seq analysis on the heterozygous diploid strain expressing dCre-Sir2 (with and without 1 µM β-estradiol induction). As a comparative control, we also performed RNA-seq on the same strain expressing dCre-only with 1 µM β-estradiol induction. We analysed the differential expression levels of the synthetic genes on *synXI* and their native copies on *chrXI* by doing pairwise comparisons among the 3 groups of tested samples (dCre-Sir2 induced, dCre-Sir2 uninduced and dCre-only induced). Induction of the dCre-Sir2 silencing system resulted in a significant downregulation of 221 genes on *synXI* (log2 fold change > 1 and p-value < 0.05), with minimal expression changes observed from *chrXI* or other native chromosomes (**Figure 5D** and **5E**). A similar gene repression profile was detected when comparing dCre-Sir2 induction to dCre-only induction, with 190 genes on *synXI* significantly downregulated in the dCre-Sir2 overexpressing samples (**Figure S13A** and **S13C**). Interestingly, expression of just dCre alone resulted in a significant downregulation of 67 genes on *synXI* upon induction (**Figure S13B** and **S13D**). These findings indicate major and specific transcriptional repression of *synXI* genes by the dCre-Sir2 silencing system. However, the dCre-only fusion protein also repressed a subset of *synXI* genes too.

To investigate the pattern of repression by dCre-Sir2 across *synXI*, we mapped log_2_ fold change gene expression along the chromosome sequence to compare differential gene expression (**Figure 5F**, **S14A**). Notably, we found that chromosomal segments where the loxPsym sites are more densely distributed exhibited strongest repression. A *synXI* left arm region immediately upstream of the centromere showed minimal gene repression and had the lowest density of loxPsym sites. In contrast, regions on the right arm and in the centre of the left arm show very strong repression and had the highest loxPsym density (**Figure 5F**). Interestingly, regions proximal to the chromosome ends showed less repression (**Figure 5F**), possibly due to fewer loxPsym sites and pre-existing silent heterochromatin at yeast telomeres that may make them less accessible for further repression. Some chromosome regions also showed consistent repression when just dCre alone was expressed (**Figure S14B**), including a region in the centre of the *synXI* left arm. However, in general the magnitude of this repression was much less.

Further analysis assessed the impact of loxPsym site proximity to the genes on silencing efficacy. For dCre-Sir2, a moderate and significant positive correlation (*R* = 0.46) was identified between the degree of gene repression and the distance between the gene’s ATG start codon and the nearest loxPsym site (**Figure 5G**). A similar correlation (*R* = 0.41) was observed when comparing dCre-Sir2 induced samples with dCre-only induced samples (**Figure S15A**). However, the correlation was notably weaker (*R* = 0.15) when comparing for repression by dCre alone (**Figure S15B**). Non-essential genes in *synXI*, especially those with loxPsym sites inserted into their 3’ UTRs, exhibited significantly more repression compared to that of essential genes when dCre-Sir2 silencing was induced (**Figure S16**). These results indicate that the proximity of loxPsym sites to the gene start codon indeed affects the efficiency of dCreSIR silencing.

## Discussion

Our findings suggest that the yeast genome may be able to tolerate genomic “defragmentation”, as shown by the successful reorganisation of tryptophan biosynthesis genes into a synthetic module. Furthermore, the dCreSIR system we developed offers a new master regulation switch for all loxPsym-containing modules and is efficient and specific and can scale to repress entire synthetic yeast chromosomes.

The standardised structure of synthetic modules coupled with advances in multiplex CRISPR/Cas editing enables module construction into various genome loci to be relatively straightforward. We observed that *TRP* gene relocation did not substantially affect function, although slightly slower cell growth was observed. This validates that genes that encode a conditional essential pathway - amino acid biosynthesis pathway - are feasible for defragmentation and can function even when inserted in non-ideal genomic locations like sub-telomere regions. This confirms the robustness of the amino acid biosynthesis pathway and of yeast itself, in terms of being able to tolerate gene reorganisation. Alongside the work of others in clustering glycolysis pathway genes^13,14^ and building a neochromosome for tRNAs genes^12^, there is now plenty of evidence that genome defragmentation is possible and should be considered for future synthetic yeast genome projects.

Placement of the *TRP* module in the “*DAL* locus 2” site inhibited tryptophan biosynthesis, possibly by the module acquiring the local silencing-chromatin state^17,34^. Insertion into the “*DAL* locus 2” site (between *DAL2* and *DAL4* genes) triggered dysregulation of local silenced genes, and perturbed the transcription of the *TRP* genes too, when the cells were grown in selective media. We speculate that this effect could be due to the upregulation of *TRP* genes mediated by GCN4 transcription factor binding to its promoters in response to tryptophan starvation^53–55^. GCN4 binding could disrupt the silenced chromatin state of *DAL* region, leading to the *DAL* genes changing their expression. In future synthetic module designs, it will be important to carefully select the genome integration sites for different synthetic modules and consider the potential local interaction of both the re-located genes and their neighbouring genes. To aid this selection, we recommend using extensive datasets from genome-wide functional assays and expression experiments^33^ to understand and predict the interactions between a chosen genome location, and local gene expression and regulation.

As well as assessing native silencing by relocating modules to heterochromatic loci, we also developed the inducible dCreSIR silencing system to repress loxPsym-containing synthetic modules. We showed the targeted silencing of a synthetic *TRP* module with its genes regulated by weak constitutive promoters and then also demonstrated the selectivity of the repression by using dCreSIR to inhibit a carotene synthesis pathway (*Crt* module) while not affecting a violacein pathway co-expressed in the same strain. In our experiments, the mRNA level of each targeted gene decreased by approximately 2-fold, as determined by reporter gene and qPCR assays. This repression is not as stringent as what is seen in previous studies that target regulators to the core promoter regions^27,43,44^. One possible explanation for reduced repression is due to the weaker binding affinity of dCre to loxPsym sites compared to transcription factors and dCas9 binding to their ideal target sequences^45,56–60^. Another explanation is that in past studies there are usually multiple repressor binding sites per gene, and these are placed close to the transcriptional start site of the promoter^61^. Both these reasons were addressed with the alternative TetR-SIR system we tested. TetR binds tightly to its target site and by incorporating 7x tetO sequences in tandem into the intergenic linkers we achieved near-complete repression of module gene expression, even for genes expressing from strong constitutive promoters. While TetR-SIR is more efficient, it is a less elegant approach to silencing gene modules as the 7x tetO array needs to be inserted into every intergenic linker in the module, and this large amount of repetitive DNA risks the genetic stability of the synthetic modules^62^. Having just a single universal site, the loxPsym, acting as the target for our system is more straightforward. It may be possible to improve the repressive potential of this site by optimising the flanking sequence of the intergenic linkers it is part of. Likewise, dCre could be improved, e.g. by directed evolution, to become a stronger loxPsym binder.

As well as recruiting CRs to a single locus, we also explored the targeted tethering of synthetic modules to the nuclear periphery using synthetic anchors including dCre-Heh1. However, we found this method to be less efficient than dCre-SIR and also caused fitness defects. A recent study employed a similar strategy but, in this case, attempted to tether the entire synthetic yeast chromosome *synIV* to the inner nuclear membrane (INM) using a custom ZF-Heh1-Heh2 fusion protein designed to target the loxPsym sequences^37^. Their chromosomal tethering approach showed significant growth defects upon induction, possibly due to ∼2-fold downregulation of chromosomal genes, particularly essential genes^37^. Consistent with our findings, a modest growth defect was observed when expressing either ZF-Heh1 or ZF-Heh2 alone, indicating potential off-target effects^37^. These off-targeting effects could be attributed to the non-specific binding of DBD or the correlation of Heh1/Heh2 with other essential cellular functions.

The hundreds of loxPsym insertions in Sc2.0 chromosomes enabled us to validate that dCre-SIR can specifically repress both at module scale (10 to 20 kb) and at whole chromosomal scale (∼0.7 Mb). More than half of genes on *synXI* were significantly downregulated upon induction of dCre-Sir2 silencing system in our RNA-seq data, and we found that targeting *synXI* by just dCre alone could also downregulate a subset of genes on *synXI,* possibly due to the synapsis activity of dCre, which loops DNA and thus facilitates inhibition of gene transcription^63^. The concept of DNA looping or bending to facilitate gene repression is well studied in bacteria^64^. Eukaryotes, including mammalian cells, exhibit a more complex DNA looping system, in which chromatin folds into a solenoid structure^65^. This chromatin compaction into a solenoid structure is known to facilitate transcriptional repression^66^. Given that Cre is known to self-assemble into tetramers^67^ this suggests a potential for dCre to compact DNA upon its binding, possibly changing the 3D structure of chromosomal DNA in a manner analogous to the solenoid structure. Thus, chromosome conformation changes and histone modifications related to epigenetic silencing are avenues worth investigating further in the future. We believe that the dCreSIR silencing system will be a versatile tool for a wide range of future applications, from extending our knowledge of genome organisation and evolution to practical uses such as for metabolic engineering and gene therapies. For future modular genomes it would be desirable to have multiple different versions of dCreSIR, induced by different small molecules and targeted different intergenic sequences, not just loxPsym.

With the Sc2.0 synthetic genome project now nearing completion^9^, understanding how to design and build new genome modules and defragmented chromosomes, and how to switch them on and off, are key step towards future minimal, modular synthetic genomes designed for specific tasks.

## Acknowledgements

We would like to thank Fankang Meng for his help in providing the (GS_5_)_6_ linker part plasmid pMFK179 and the TetR part plasmid pMFK488, Czarina Choi Lo for providing her strain yCL002 as the base strain for testing our dCreSIR silencing system, Jack Ho for providing the *vio* genes part plasmid and Cre recombinase part plasmid, Klaudia Ciurkot, Benjamin Blount, Patrick Yizhi Cai for their support and advice, and Sud Pinglay for telling us that we’d already come up with the perfect name for our dCreSIR system before we’d even realised.

## Methods

### Strains

All yeast strains generated in this study are derived from BY4741 yeast (*<u>MATa</u> his3Δ1 leu2Δ0 met15Δ0 ura3Δ0*)^68^, including the Sc2.0 project strain containing the now-published *SynXI* chromosome^52^. See **Table S1** for a full list of derived yeast strains. NEB Turbo competent *Escherichia coli* (*E. coli*) from New England Biolabs (NEB was used for all DNA cloning and plasmid propagation work.

### Growth media and conditions

<u>Y</u>east extract <u>P</u>eptone Dextrose (YPD) media (10 g L^−1^ yeast extract (VWR), 20 g L^−1^ peptone (VWR), 20 g L^−1^ glucose (VWR)) was used for general culturing of yeast cells. <u>S</u>ynthetic <u>C</u>omplete media (SC; 6.7 g L^−1^ Yeast Nitrogen Base without amino acids, 1.4 g L^−1^ Yeast Synthetic Drop-out Medium Supplements without L-uracil, L-tryptophan, L-histidine, L-leucine, 20 g L^−1^ glucose) was used for auxotrophic selection experiments, or was used with all amino acids supplemented as a defined complete medium. Amino acids such as 20 mg L^−1^ L-tryptophan, 20 mg L^−1^ L-histidine, 20 mg L^−1^ uracil and 120 mg L^−1^ L-leucine were supplemented into SC media depending on the required auxotrophic selection. For growth on plates, media were supplemented with 20 g L^−1^ bacto-agar (VWR). All other components used in the media were supplied by Sigma Aldrich.

Luria-Bertani (LB) medium was used for culturing *E. coli*. LB agar was prepared by dissolving 37 g L^−1^ LB agar powder (VWR) into required amount of distilled water. Antibiotics such as ampicillin (100 μg mL^−1^), chloramphenicol (25 μg mL^−1^), kanamycin (50 μg mL^−1^) and spectinomycin (100 μg mL^−1^) were supplemented when necessary.

### Plasmids

gRNA plasmids were generated by T4 PNK phosphorylating (NEB) and annealing two oligos, followed by a BsmBI Golden Gate assembly to insert the small fragment into the SpCas9 sgRNA Dropout vector (Ellis lab plasmid pWS2069). Oligos for gRNAs were designed by adding the sequence 5’-AGAT-3’ at the 5’ end of the forward primer and 5’-AAAC-3’ at the 5’ end of the reverse primer. For a full list of their sequence, see in **Table S5**.

Donor DNA plasmids used for CRISPR/Cas9 genome editing were generated by cloning the gap repair donors into the pYTK001 entry vector by BsaI Golden Gate assembly. Donor DNA sequences consist of the 500 bp homology regions amplified from the BY4741 genomic DNA, flanking a 23 bp landing pad sequence serving as the CRISPR/Cas9 target^28^. The donor DNAs were then amplified from these pre-assembled plasmids for yeast transformation alongside the CRISPR DNA to facilitate homologous-directed-repair (HDR) facilitated targeted yeast genome editing. For a full list of plasmids with donor DNAs, see in **Table S2**.

The gene fragments for the assembly of the defragmented synthetic modules were firstly constructed into pYTK001 entry vector using a homemade enhanced Gibson Assembly master mix, adapted from a recipe in a preprint by Rabe *et al*.^69^ For a full list of plasmids with gene fragments, see in **Table S2**.

The defined linkers used to link the gene fragments were generated by inserting the loxPsym sequence into the centre of the connectors by amplifying the whole plasmid sequence of the entire connector plasmids using forward and reverse primers each containing half of the loxPsym sequence. The PCR amplicon was ligated as a plasmid through T4 PNK phosphorylation and ligation. For a full list of part and pre-assembled vector plasmids with synthetic linkers, see in **Table S3** and **Table S7**, respectively.

Cassette and multi-cassette plasmids containing the transcription units of the genes in tryptophan (refactored), violacein, and carotene biosynthesis pathways and for targeted genomic integration of the dCre-CR silencing system were assembled using the MoClo Yeast Toolkit (YTK) system^29^ into the pre-assembled Dropout vectors containing the defined linkers with loxPsym sites. For a full list of part plasmids, cassette-level plasmids and multi-cassette level plasmids, see in **Table S6**, **Table S8** and **Table S9**, respectively.

DNA sequences being assembled through Golden Gate Assembly in this study are free of recognition sites for the type IIs restriction enzymes used by the Golden Gate reaction, i.e. BsmBI, BbsI, or BsaI. For the assembly reactions, plasmids or DNA fragments used in the Golden Gate assembly were normalised to the concentration of 50 nM. Golden Gate assembly reactions were typically prepared by mixing the components as follows: 1 μL of each plasmid or DNA fragment, 1 μL of entry vector (backbone), 0.5 μL of type IIs restriction enzyme (BsaI, BsmBI or BbsI-HF) (NEB), 0.5 μL of T4 DNA Ligase (NEB), 1 μL of T4 DNA ligase buffer (NEB), made up to 10 μL with nuclease-free water. The mixtures were then transferred to a thermocycler using the Golden Gate assembly program: 25 cycles of (2 min at 42°C or 37°C, 5 min at 16°C), followed by 10 min at 60°C for a final digestion, and 10 min at 80°C for heat inactivation. The entire reaction was then transformed into *E. coli*.

### CRISPR/Cas9 genome engineering

For multiplex gene deletion, 50 ng of the CRISPR/Cas plasmid (Ellis lab plasmid pWS2081 – *URA3*), 600 ng of each sgRNA plasmid were mixed together with 0.5 μL BpiI (ThermoFisher), 1 μL of 10X Buffer G (ThermoFisher), and nuclease-free water to make up to 10 μL. This mixture was then incubated at 37°C for 8 hours followed by 80°C heat inactivation for 10 min. 5 μg of each donor DNA was added to this mixture to a total volume of 64 μL to be used for the yeast transformation. Donor DNA was generated by PCR amplification from the pre-assembled donor DNA plasmids and purified by DNA Clean & Concentrator Kit (Zymo Research).

For genome integration and other genome editing experiments, 250 ng of the CRISPR/Cas plasmid and 500 ng of each DNA fragment was combined with 10 μL boiled salmon sperm DNA, made up to 64 μL with nuclease-free water to be used for the yeast transformation. DNA fragment was generated by PCR amplification and purified by DNA Clean & Concentrator Kit (Zymo Research). gRNA plasmids were constructed according to the ‘gRNA-tRNA Array Assembly’ methods described in the Multiplex MoClo Toolkit^29^. Individual gRNA-tRNA fragment was amplified by setting up a 50 μL of Q5 PCR reaction according to the manufactures’ instructions, using 1 μL of diluted pWS3178 plasmid (∼2 ng μL^−1^) as the template and 1 μL of each primer (100 μM, ordered from IDT). After gel electrophoresis, gel bands were cut and DNA was isolated using Zymoclean Gel DNA Recovery kit (Zymo Research), following the manufacturer’s instruction. The purified DNA fragments were then assembled into pWS3932 by a BsaI Golden Gate assembly.

For a full list of gRNAs used for CRISPR/Cas9 genome engineering, see **Table S5**.

### DNA extraction

Yeast genomic DNA for PCR verifications was isolated following a LiOAc/SDS isolation protocol^70^. Colonies were inoculated into appropriate media and grown overnight to saturation (30°C, 250 rpm). 200 μL of overnight culture was pelleted by centrifugation at 8000 rpm for 1 min. The pellet was then resuspended in 200 μL of LiOAc/SDS solution (200 mM LiOAc, 1% (w/v) SDS) and incubated at 70°C for 5 min, followed by adding 300 μL of 100% ethanol. The suspension was then vortexed and centrifuged at 15,000 g for 3 minutes. After removing supernatant, the pellet was resuspended in 100 μL nuclease-free water, following a 20 s final spin at 15,000 g. 80 μL of the supernatant was then transferred to a new tube and can be directly used as the template for Q5 PCR.

Plasmids were propagated by transformation in Turbo Competent *E. coli* cells. Plasmid DNA was isolated according to the Qiagen Miniprep protocol following the manufacturer’s instructions, using the homemade buffers (https://openwetware.org/wiki/Qiagen_Buffers). Alternative purification columns (NBS Biologicals) were also used in some cases but following the same protocol. DNA quantification was analysed using the Nanodrop 1000 spectrophotometer (ThermoFisher).

### *E. coli* transformation

Chemically competent cells were prepared following the TSS protocol for KCM transformations^71^. Turbo Competent *E. coli* cells were firstly streaked onto LB agar plates to obtain single colonies. Next day, a colony was picked out from the plate and grown to saturation overnight (37°C, 250 rpm) in 10 mL LB. 5 mL of this culture was inoculated into a 2 L baffled flask with 500 mL LB, and then grown to OD_600_ ∼ 1.0 (37°C, 250 rpm). Cells in this flask were chilled on ice to stop growth, then separated into 50 mL conical tubes, and harvested by centrifuging at 4°C (4000 rpm, 10 minutes). Cell pellets were resuspended in the ice-cold TSS buffer (85 mL LB, 10 g PEG-3350, 5 mL DMSO, and 2 mL 1 M MgCl_2_) and then aliquoted into PCR tubes. Each tube containing ∼200 μL of the cell suspension was rapidly frozen on dry ice and stocked in the −80°C freezer.

For DNA transformation, 50 μL of 5 x KCM (250 mM MgCl_2_, 500 mM KCl, 150 mM CaCl_2_) was added into the thawed Turbo Competent *E. coli* cells. 50-80 μL of this mixture was then added to the DNA (1-10 μL) and transferred to a thermocycler following the heat-shock program: 4°C for 10 min, 42°C for 1 min, 4°C for 1 min and 37°C for 40-60 min. This mixture was then plated onto the LB agar supplemented with specific antibiotics for selection.

### Yeast transformation

A colony was picked out from the plate and grown to saturation in 2 mL appropriate media overnight (30°C, 250 rpm). The next day, cell culture was diluted to OD_600_ ∼0.2 in a 50 mL Falcon centrifuge tube with 10 mL fresh media and grown for ∼6 h to OD_600_ = 0.8-1.0. Cells were pelleted by centrifuging at room temperature (2000 rcf, 10 minutes), then washed once with 10 mL 100 mM lithium acetate (LiOAc, Sigma Aldrich). Centrifugation was repeated and cell pellet was resuspended in ∼600 μL 100 mM LiOAc. 100 μL of this mixture was added to 64 μL DNA cocktail containing 10 μL of boiled salmon sperm DNA (ThermoFisher) per transformation, and then gently mixed with 296 μL PEG-3350/LiOAc mixture (260 μL 50% (w/v) PEG-3350 and 36 μL 1M LiOAc). This mixture was then placed in a heat block at 42°C for 40 min and cells were pelleted by centrifugation at 8000 rpm for 1 min. Pellets were resuspended in 100-200 μL 5 mM CaCl_2_ and plated onto the appropriate agar media for selection.

### Plasmid curing

For curing *URA3* containing plasmids, 5-FOA (5-fluoroorotic acid) counterselection was used^72^. Colonies were inoculated into 2 mL YPD media and grown overnight (30°C, 250 rpm). This culture was streaked using a 10 μL loop onto the agar plate supplemented with 5-FOA (Formedium). Colonies showing growth after incubation for 3 days at 30°C suggested successful plasmid curing.

For an auxotrophic marker that is not *URA3*, strains were firstly streaked onto the YPD agar plate. After 3 days of incubation at 30°C, colonies were picked and inoculated into 2 mL of fresh YPD media and grown overnight (30°C, 250 rpm). This culture was then streaked using a 10 μL loop onto the YPD agar plate and incubated at 30°C for 3 days. Colonies were then streaked onto both the agar media plates with selection and also agar media plates lacking selection. Strains showing growth on the media lacking selection but not the media with selection suggested successful plasmid curing.

### Yeast mating

A heterozygous diploid strain was generated by streaking two haploid strains onto YPD agar, incubating at 30°C for 2 days and then mixing patches of colonies from cells of the opposite mating type together on a fresh YPD agar plate. Cells were incubated for 4 h at 30°C before re-streaking onto a fresh media plate with appropriate selection for further growth.

### Standard PCR

Standard Q5 high-fidelity DNA polymerase (NEB) PCR conditions and protocols were used for the DNA amplification. The amount of template used in a 50 μL Q5 PCR reaction was 1 μg for genomic DNA or 10 ng for plasmid DNA.

GoTaq Green (Promega) PCR was used for direct PCR from yeast and bacterial colonies when doing colony PCR for screening. A colony was picked into 50 μL nuclease-free water and 1 μL of this mixture was used as the PCR template. 5 μL GoTaq Green Master Mix, 2 μL of each 10 μM primer and 1 μL of the template, were mixed, bring to a total volume of 10 μL. DNA purification after the PCR and gel electrophoresis was performed using the DNA Clean & Concentrator Kit (Zymo Research) and Zymoclean Gel DNA Recovery kit (Zymo Research), respectively, according to the instructions from the manufacturers.

### Plate reader assay

Overnight cultures were harvested, washed and used to inoculate 100 μL cultures in a 96-well plate with a starting OD_600_ normalised to 0.02. Plates were incubated and measured in a Synergy HT Microplate Reader (Biotek) shaking at 30°C. Mean absorbance values of equivalent blank media wells were subtracted from data points. Mean fluorescence values of equivalent blank media wells were subtracted from data points.

### Spot assay

Saturated overnight yeast cultures were used to inoculate 5 mL of appropriate media. Cultures were grown to mid-exponential phase for 4-6 hours, normalised to OD_600_ = 1, pelleted by centrifugation, washed in water, pelleted again and resuspended in water. Washed normalised cells were serially diluted in water in one-in-ten steps. Diluted cells were plated in 8 μL spots onto appropriate media plates and incubated at 30°C for the assay.

### Standard microscopy

A single yeast colony was used to inoculate 2 mL of appropriate media. Cultures were grown overnight (30°C, 250 rpm) and visualised on a Nikon Eclipse Ti inverted microscope at 20x magnification and optical conf. Bright field (BF) images were captured using the Nikon NIS-Elements Microscope Imaging Software. Fiji^73^ was used to process the images and add the scale bars.

### Flow cytometry

The fluorescence of cells was measured by an Attune NxT Flow Cytometer (ThermoFisher). The following settings were used for measuring the size of the cell, complexity of the cell and fluorescence of the cell: FSC 100 V, SSC 355 V, BL1 450 V. 10,000 events of yeast population gated by forward and side scatter were collected for each experiment and analysed by FlowJo. Geometric means of the fluorescence distributions were calculated by FlowJo. The autofluorescence value of the non-fluorescent cells that have no silencing cassette integrations was subtracted from these values. ‘Normalised GFP expression’ values were calculated as the ratio of fluorescence values from dCre-CR integrated to those with dCre-only integrated. All values obtained were the means of three biological repeats. Unless otherwise stated, data visualisation was performed using GraphPad Prism.

### Carotenoid extraction and quantification

Colonies were inoculated into appropriate selective media and grown overnight to saturation. Saturated cultures were washed twice in water and resuspended in 1 mL PBS. Culture was diluted to an OD_600_ of 0.1 in 2 mL fresh media and grown for 2 days. Cells were pelleted by centrifugation at 4000 rpm for 2 min. Supernatants were discarded and the pellets were then resuspended in 1 mL acetone (VWR), followed by adding 200 μL glass beads (Sigma). Cells were lysed using a homogeniser (VWR) by beads beating 8 times at 8000 rpm, with 1 min ON and 15 s OFF pulses. The supernatant was filtered by a 0.22 μm filter (VWR). 500 μL of the supernatant was taken for absorbance measurement on a spectrophotometer at 453 nm. The measured absorbance was then converted into concentrations using standard curves of β-carotene ranging from 0 to 5 mg L^−1^, 1 to 10 mg L^−1^, 5 to 100 mg L^−1^.

### Whole genome sequencing and data analysis

Whole genome sequencing of yeast strains was performed by SeqCenter, Pittsburg PA, USA using an Illumina NextSeq 500 system with pair-end protocol. All raw reads following sequencing were processed to remove the adaptor sequence using the Trimmomatic tool, and then mapped to the reference genome sequences using BWA-MEM and Samtools on the Galaxy platform^74^. The genomic coverage for each locus was analysed and plotted by tinycov (https://github.com/cmdoret/tinycov.git) from BAM files generated by the Galaxy platform.

### RNA isolation

Cells were grown in YPD media at 30°C until mid-exponential growth phase (OD_600_ ∼2). Cell culture corresponding to ∼3 × 10^8^ cells was harvested by centrifugation, washed in 0.8% physiological salt solution and resuspended in 500 mL solution of 1M sorbitol and 100 mM ethylenediaminetetraacetic acid (EDTA). Spheroplasts were generated by digesting cells with 50 U Zymolyase (Zymo Research) at 30°C for 30 min. Spheroplasts were collected by centrifugation and RNA was isolated using the NucleoSpin RNA Plus kit (Macherey-Nagel). RNA quality and integrity was determined by Qubit fluorometry using an RNA BR Assay Kit (ThermoFisher), spectrophotometry with a NanoDrop (ThermoFisher) and on a 2100 BioAnalyzer using an RNA 6000 Nano Kit (Agilent).

### RT-qPCR

2 mg of the isolated RNA was digested with DNAse I (Roche) and cDNA was synthesized using the GoScript reverse transcription kit A5001 (Promega), according to the manufactures’ instructions. PCR assays were performed using the Luna Universal qPCR Master Mix (NEB) in a MasterCycler RealPlex 4 (Eppendorf). Each 20 μL qPCR reaction contained 90 ng of cDNA. The fold change of gene expression was calculated using the DDCt method^75^, using *ACT1* as the reference gene. Two technical repeats were performed for each of three biological replicates. Primers used for qPCR are listed in **Table S4**.

### RNA-Seq and data analysis

RNA sequencing was performed by Azenta Life Sciences. Briefly, mRNA was purified from total RNA with poly-T oligo-attached magnetic beads prior to cDNA synthesis, adaptor ligation and sequencing on an Illumina platform.

RNA-Seq data was analysed using a custom pipeline developed by Anima Sutradhar and Dr Giovanni Stracquadanio from the University of Edinburgh. Briefly, the Illumina unstranded paired-end reads were pre-processed by trimming adapters and removing low quality bases using fastp^76^. Then, a reference genome of the heterozygous diploid strain BY4742 x *synXI* was created with annotations. An annotated reference transcriptome was created by considering only protein-coding genes. Reads were aligned to the references using STAR^77^. Transcripts abundance of genes was quantified using featureCounts^78^ and then differential expression analysis carried out using DESeq2^79^ with default settings, and using independent filtering to optimise the number of adjusted p-value, obtained using Benjamini-Hochberg procedure, at the significance threshold ɑ= 0.05. Fold change estimates for volcano plots were obtained using the fold change shrinkage function in DESeq2^79^.

## Supplementary Figures

**Figure S1.**
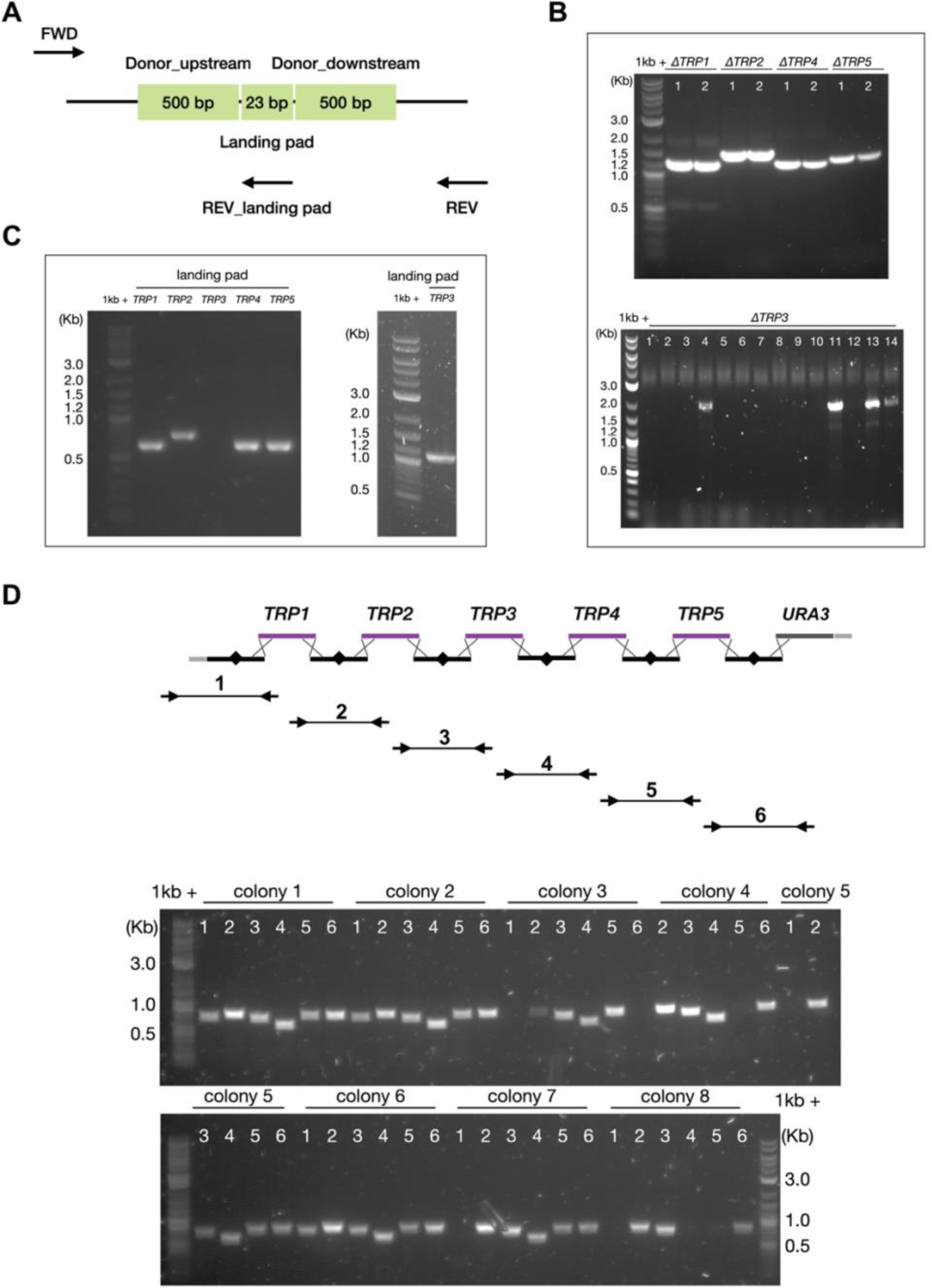
Deletion of *TRP* genes using markerless CRISPR editing and colony PCR verifications. (A) Schematic of primer design to identify *TRP* gene deletion and landing pad insertion. (B) Colony PCR of transformants to identify *TRP* gene deletion. (C) Colony PCR of transformants to identify landing pad insertion. Numbers indicate the various tested colonies. (D) Schematic of primer design and colony PCR of transformants to identify synthetic *TRP* module integration. Arrows in black represent the primers targeting at the junctions for PCR. Lines connected the arrows indicate the various tested junctions. Numbers indicate the various tested junctions.

**Figure S2.**
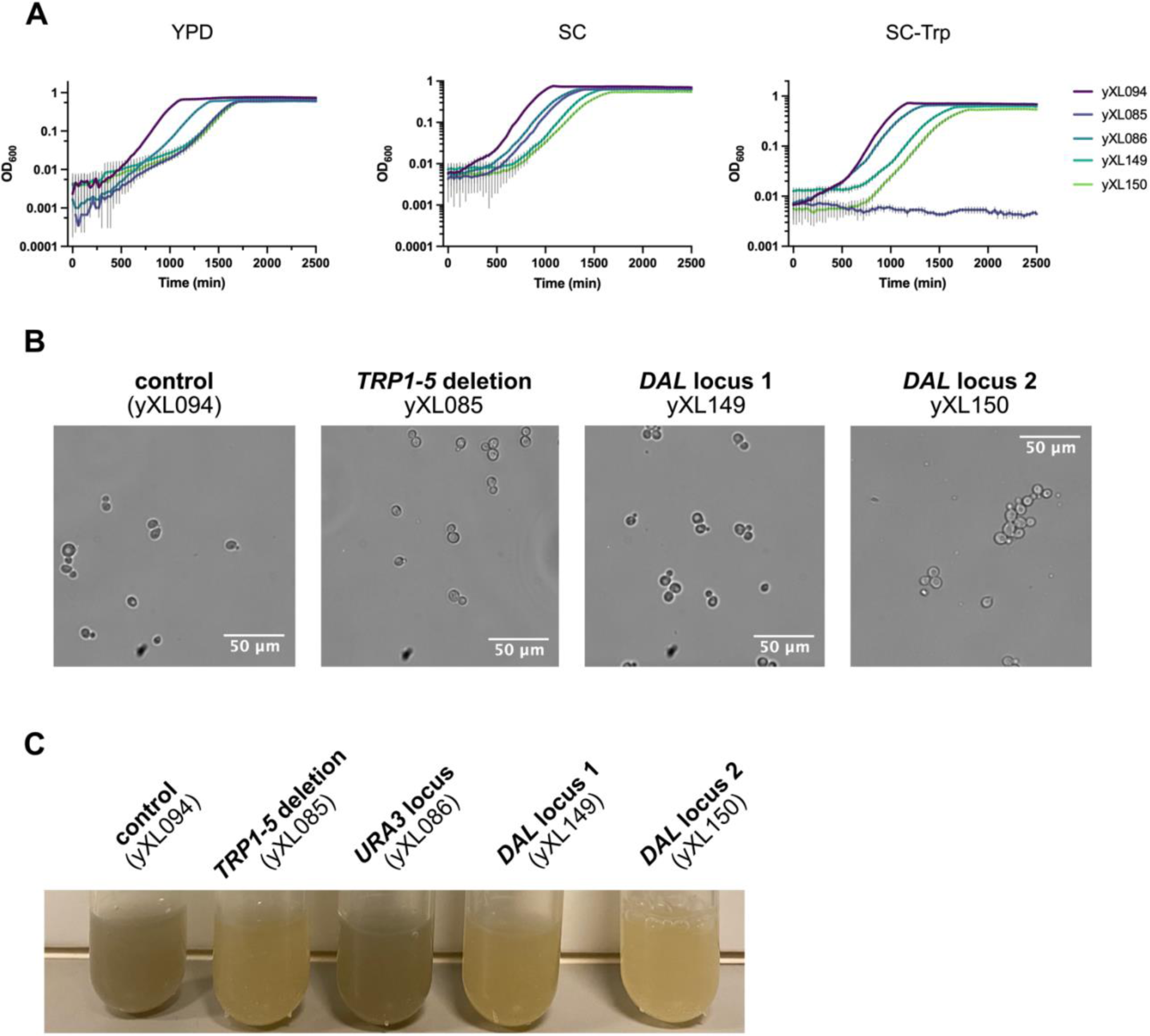
Growth and morphology assay of the strains with the defragmented *TRP* module at various genomic positions. (A) Growth curves of the strains yXL094, yXL085, yXL086, yXL149 and yXL150 in YPD, SC and SC-Trp, n=3. (B) Microscopy images of the strain yXL094, yXL085, yXL149 and yXL150. (C) Cultures of yXL094, yXL085, yXL086, yXL149 and yXL150 after growing for 48 hours in YPD.

**Figure S3.**
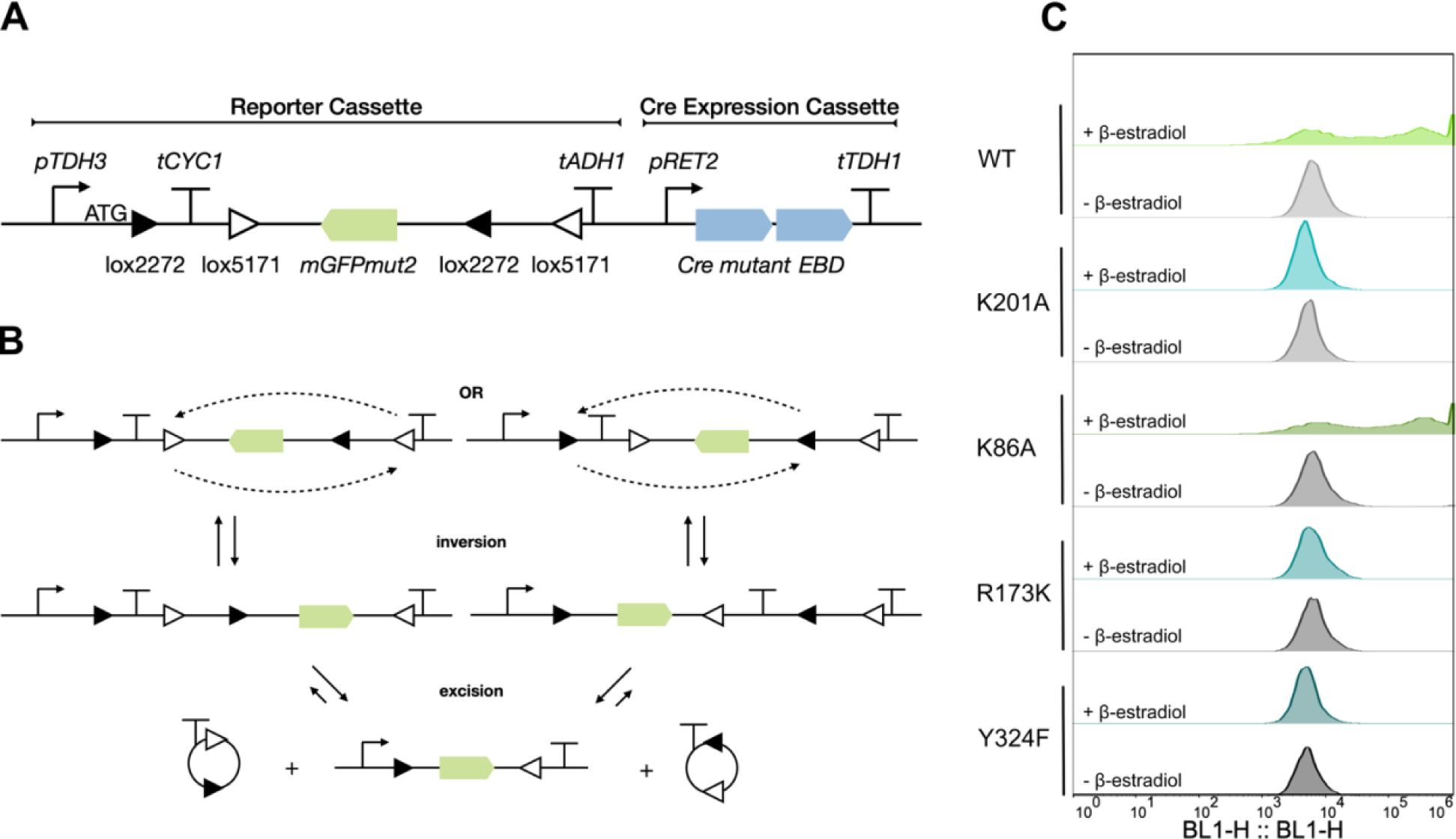
Validation of the Cre recombinase mutant that binds loxPsym but does no cleavage. (A) Schematic of the design of the validation system adapted from FLEX Cre-Switch system to examine the excision inactivity of Cre recombinase mutants using a GFP reporter. (B) Schematic of the process to validate the Cre excision activity by generating the flip of the *mGFPmut2* gene. The validation system consists of an antisense-orientated GFP gene flanking by two pairs of orthogonal loxP variant sites, namely loxP2272 and loxP5171. In the presence of Cre, two steps of recombination – inversion and excision occur at each pair of loxP variant sites respectively, resulting in the stable inversion of the GFP gene and deletion of one from each pair of recombination sites. (C) Population of GFP fluorescence, determined by flow cytometry in the presence and absence of 1 µM β-estradiol, of strains expressing various Cre mutants.

**Figure S4.**
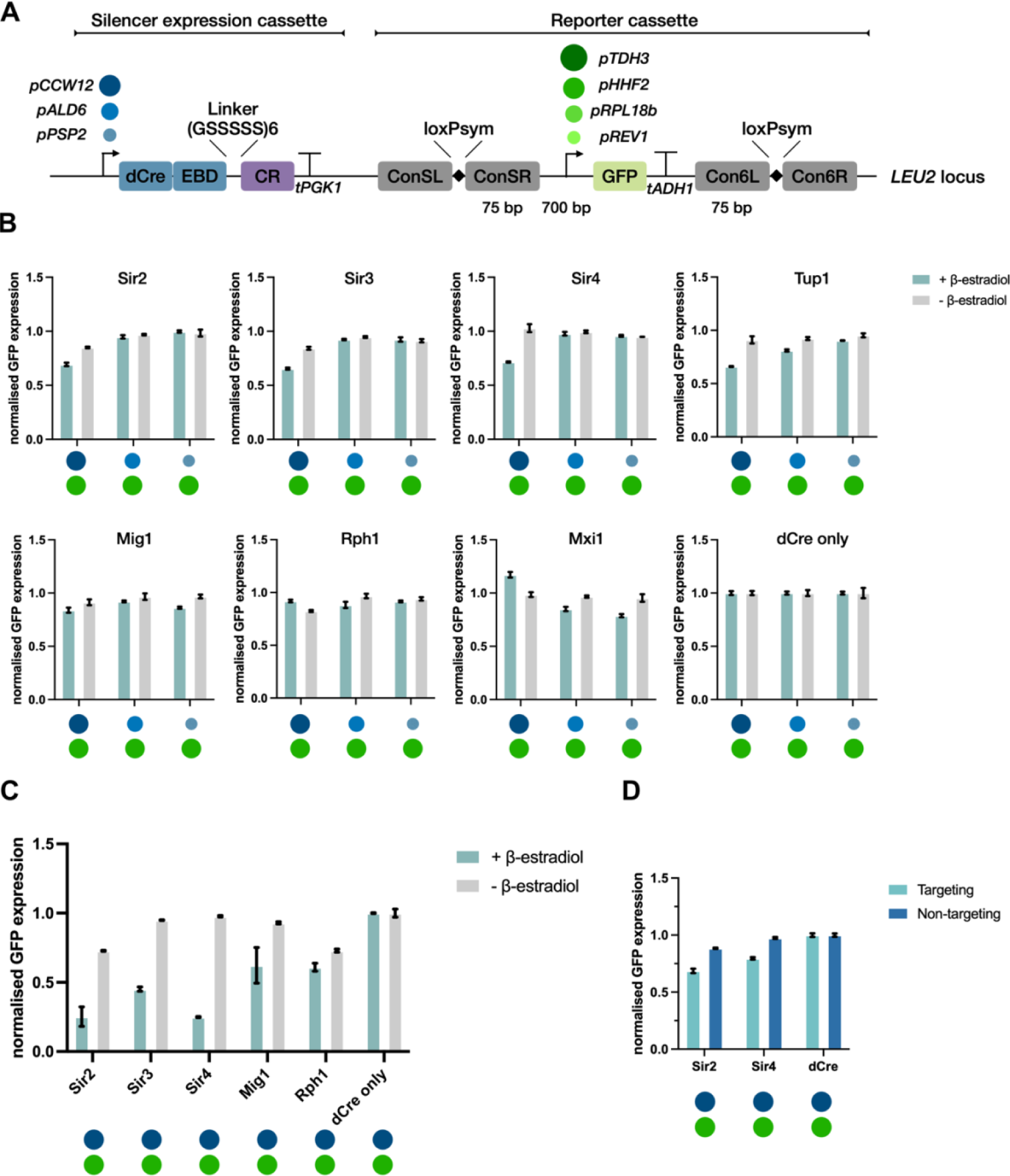
Characterising the transcriptional regulation of the dCre-chromatin regulators (CRs) using a fluorescent reporter, related to Figure 3. (A) Schematic of the cassette design of the transcriptional reporter system to analyse the silencing effects of different chromatin regulators. The promoters regulating the CRs and sfGFP expression are represented in circles with different colours. (B and C) Fluorescence measurements of sfGFP expression per cell determined by flow cytometry in the presence and absence of 1 µM β-estradiol across the various dCre-CR fusions after (B) 6 hours and (C) 29 hours. Relative sfGFP expression is normalised to a dCre-only control, shown as the mean ± SD, n=3. (D) Single cell sfGFP expression level normalised to a dCre-only control showing the targeting and non-targeting effects of dCre-Sir2, dCre-Sir4 and dCre driven by a *CCW12* promoter, n=3.

**Figure S5.**
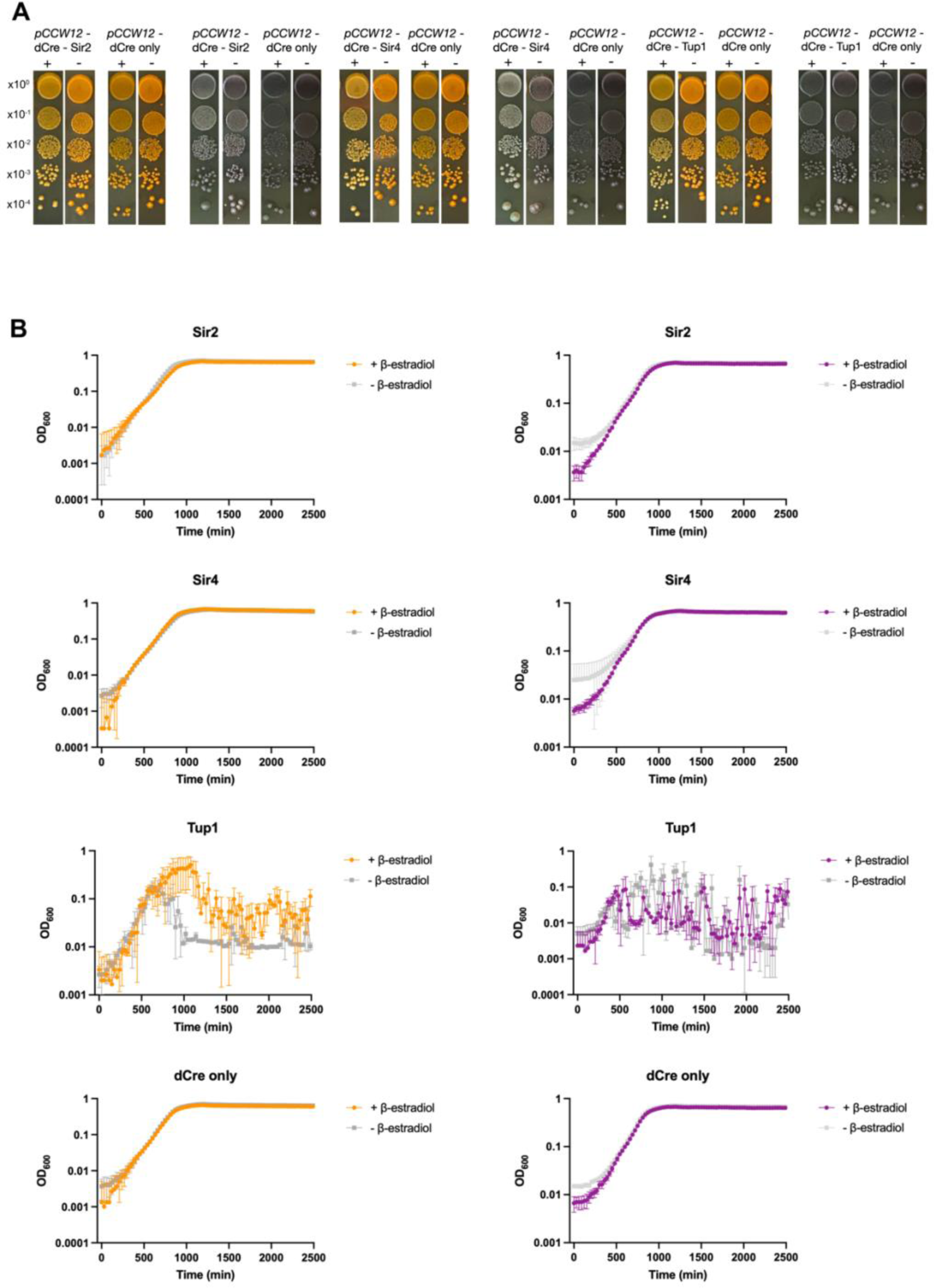
Characterising the silencing effects and the fitness cost of dCre-CR overexpression. (A) Spot assays showing the silencing effects on a carotene biosynthesis pathway (colonies producing orange pigments) and a violacein biosynthesis pathway (colonies producing dark green pigments) when inducing the dCre-CR system in YPD media. Cultures normalised to OD_600_ = 1.0 were serially diluted and spotted from top to bottom. Images were taken from the same plate, cropped, and reorganised to show comparisons. (B) Growth curves of the selected strains from panel (A) when culturing in SC-Leu under induced and uninduced conditions, n=3.

**Figure S6.**
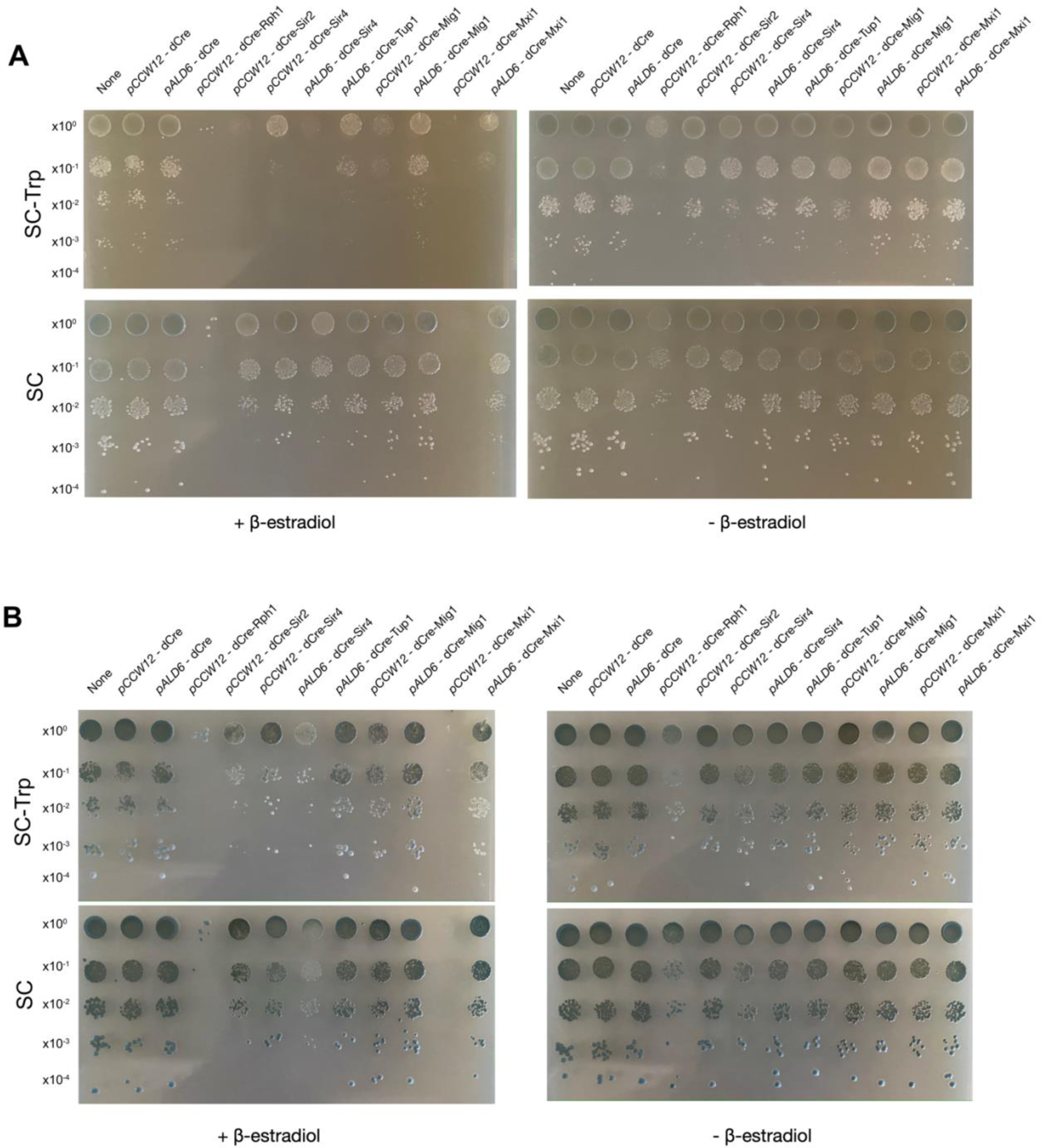
Characterising the silencing effects of dCre-CRs using a synthetic *TRP* module coupled with the violacein reporter in a yeast strain (yCL002) that has each *TRP* gene expressed from a weak constitutive promoter, related to Figure 3. Spot assays showing the silencing effects on the recoded *TRP* module when overexpressing various dCre-CR fusions. Strains were grown on SC and SC-Trp plates for 3 days with and without 1 µM β-estradiol. Cultures normalised to OD_600_ = 1.0 were serially diluted and spotted from top to bottom. Images were taken on the second day (as shown in panel A) and the third day (as shown in panel B) of the spot assay experiment.

**Figure S7.**
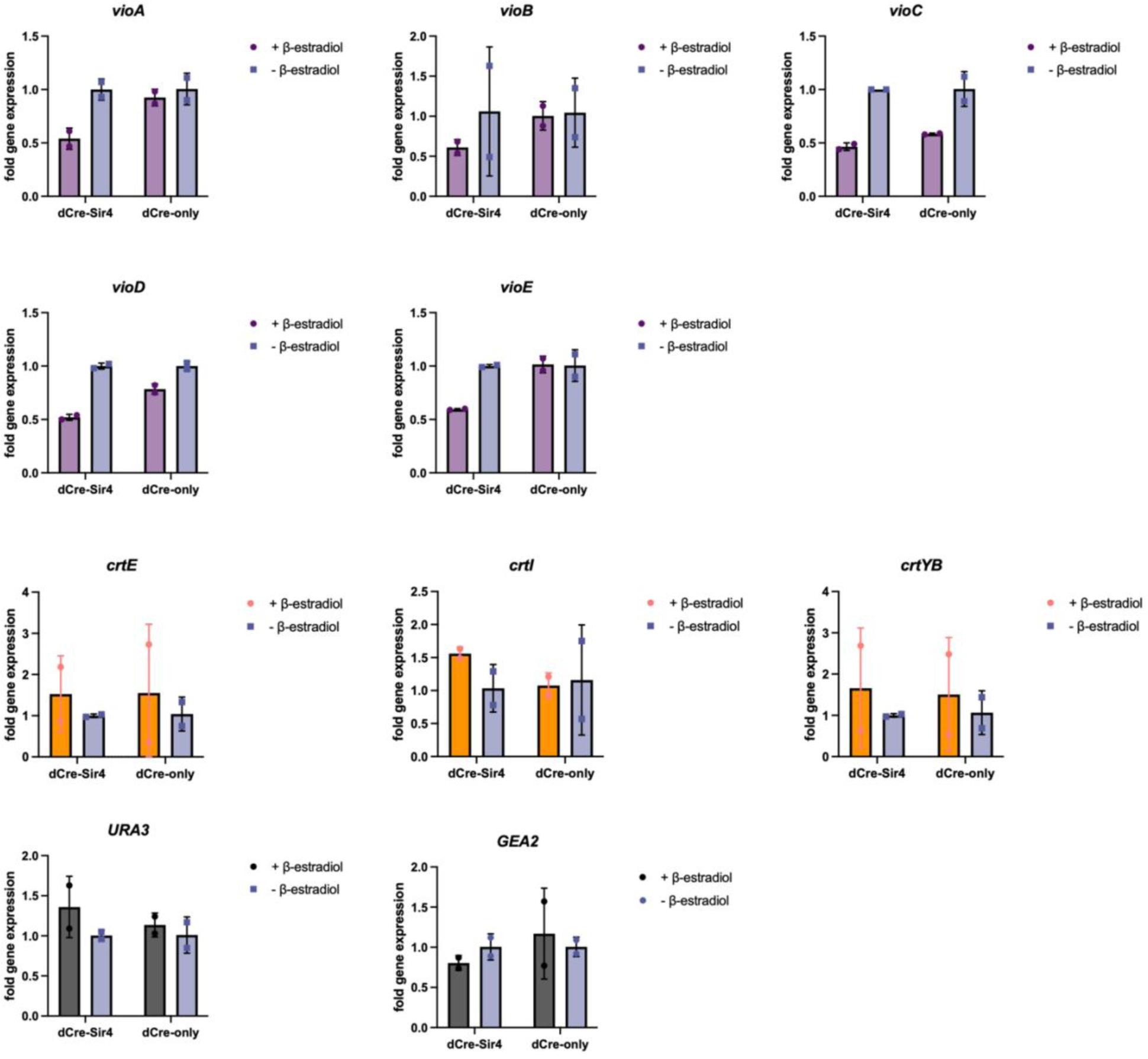
Quantification of transcript levels of genes that are synthetically targeted and untargeted silenced. Yeast strains (yXL276 and yXL277) simultaneously expressing a carotene biosynthesis pathway and a violacein biosynthesis pathway were grown for 6 hours in YPD with or without induction. Transcripts of the genes in the targeted cluster (*vioA*, *vioB*, *vioC*, *vioD* and *vioE*), genes in the untargeted cluster (*crtE*, *crtI*, *crtYB*), and neighbouring genes of the targeted region (*URA3* and *GEA2*), were quantified by qPCR using *ACT1* as a reference gene. qPCRs were performed in technical triplicates. Individual data points from samples induced with 1 µM β-estradiol are plotted as round dots, while the data points from the uninduced samples are plotted as squares. Mean averages of fold change in gene expression are shown as the mean ± SD, n=2.

**Figure S8.**
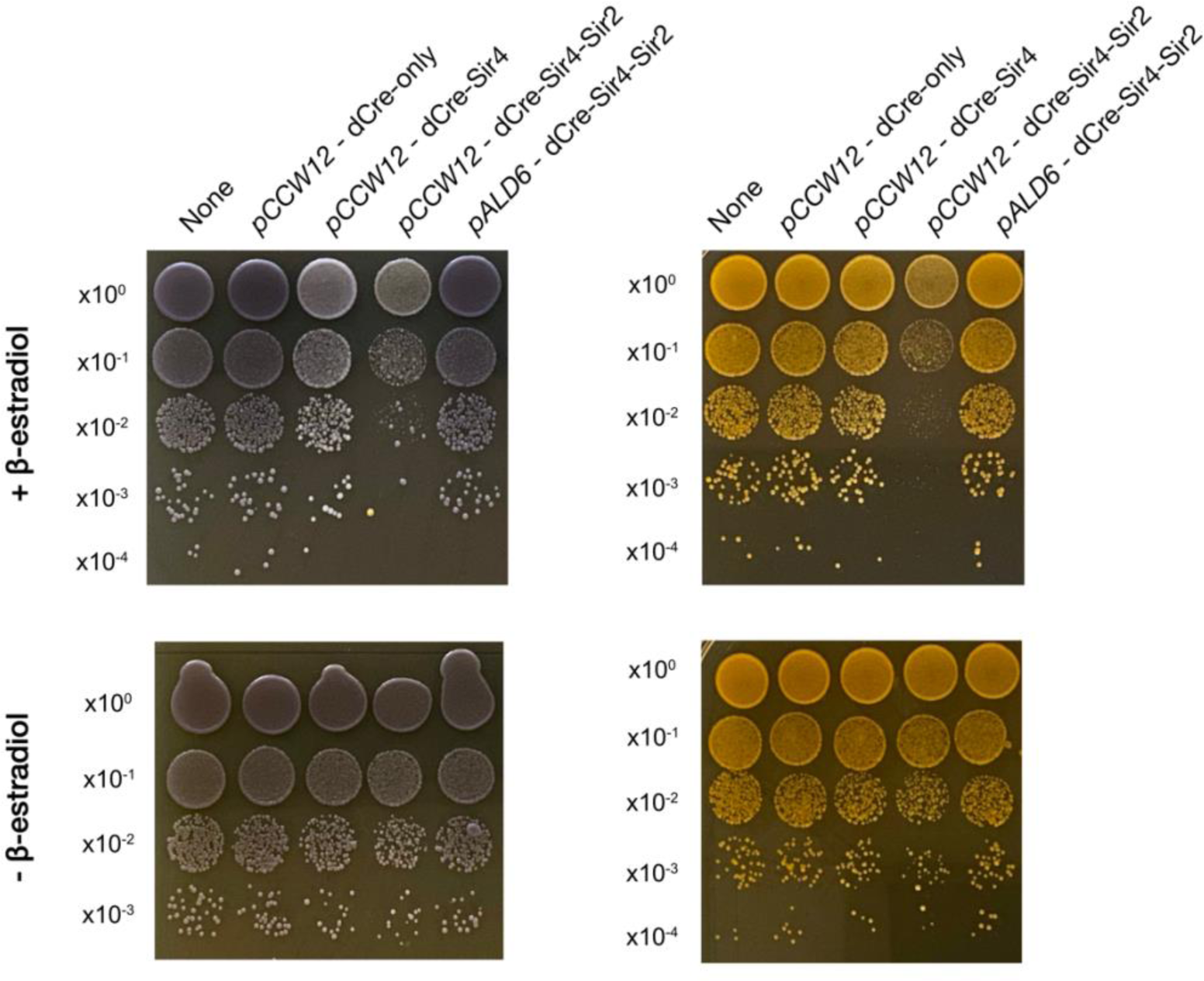
Characterising the silencing effects of dCre-Sir4-Sir2 fusion. Spot assays showing the silencing effects on the violacein and β-carotene biosynthesis pathways when overexpressing dCre-Sir4-Sir2 fusion driven by *pCCW12* and *pALD6* promoters, respectively. Cultures normalised to OD_600_ = 1.0 were serially diluted and spotted from top to bottom. Images were taken after 2 days of incubation on YPD plates with and without 1 µM β-estradiol.

**Figure S9.**
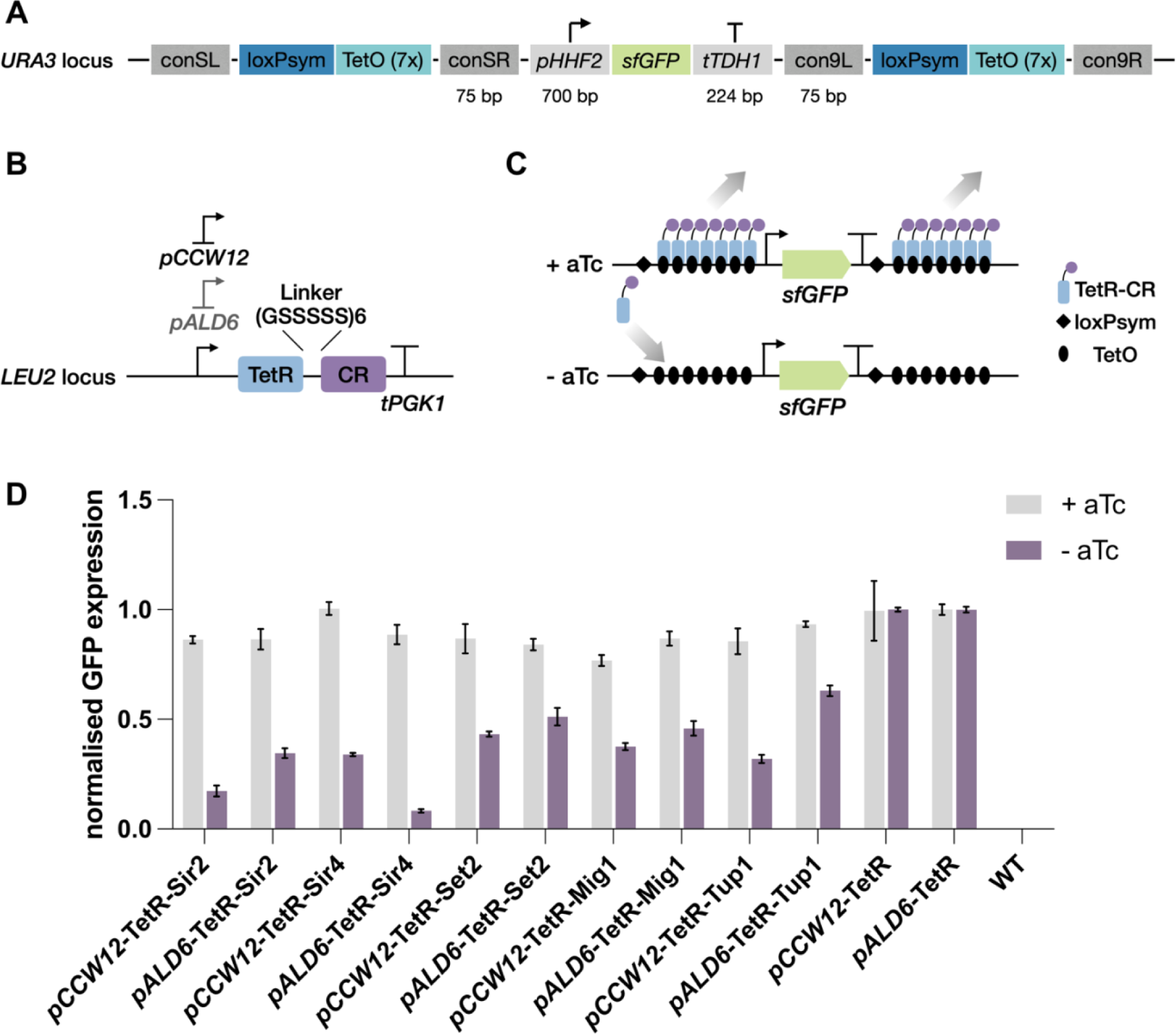
Characterising the transcriptional regulation of the TetR-CR fusions using a TetO sponge reporter. (A) Schematic of the cassette design of a *sfGFP* reporter to determine the silencing effects of various Tet-CR fusions. (B) Schematic of the TetR-CR fusion protein expression cassette for integration at the *LEU2* locus. (C) Diagram showing the sponged TetR-CR regulation on individual *crt* gene in the presence and absence of 1 µM aTc. (D) sfGFP fluorescence per cell determined by flow cytometry in the presence and absence of 1 µM aTc across the various TetR-CR fusions after growing for 6 hours. Relative sfGFP expression is normalised to a TetR-only control, shown as the mean ± SD, n=3.

**Figure S10.**
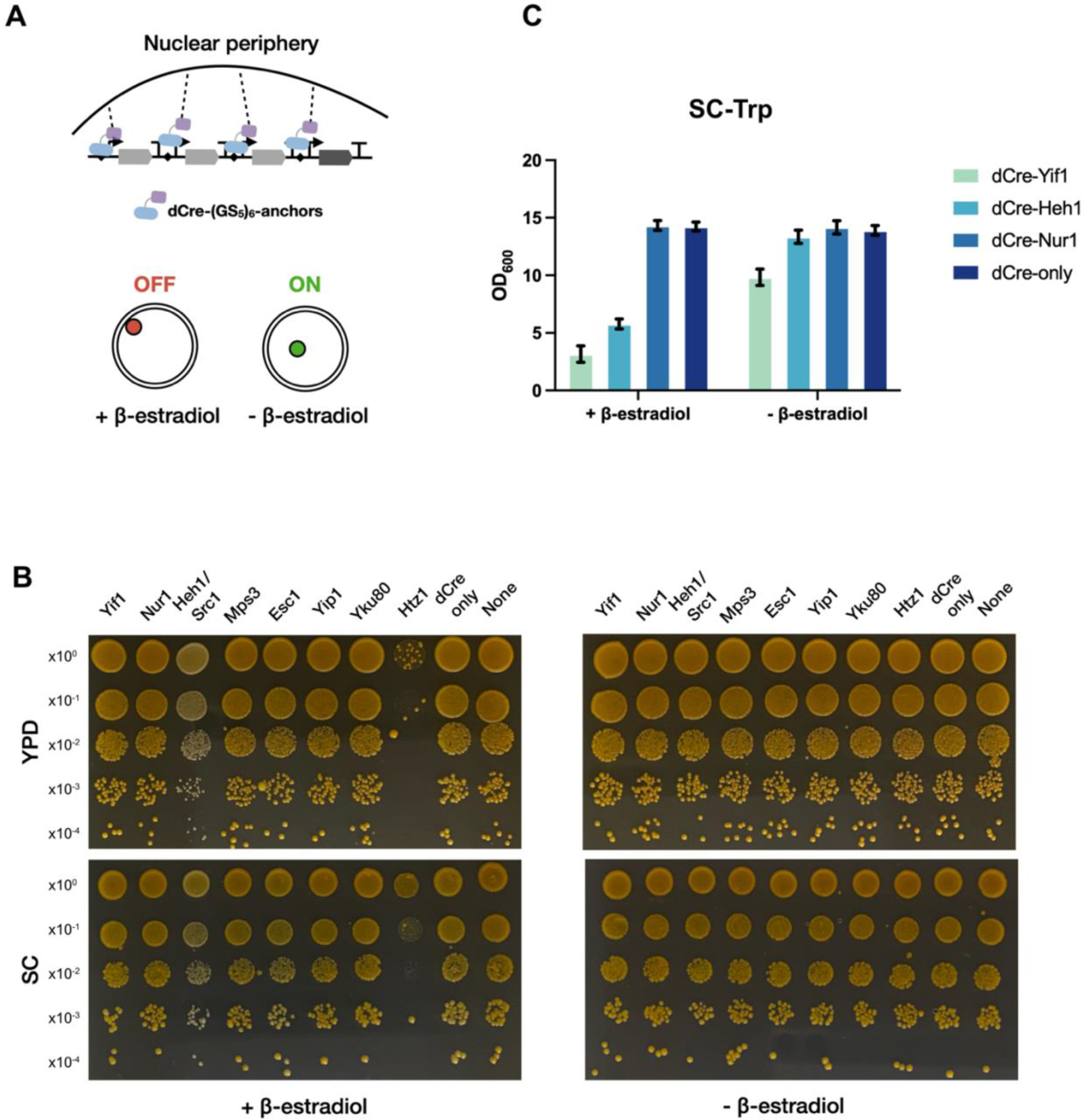
Multigene cluster silencing by synthetic tethering a genomic region to the nuclear periphery. (A) Schematic of relocating a synthetic cluster containing loxPsym sites to the nuclear periphery to simultaneously silence multiple genes at one single genomic locus utilising synthetic tethers, which are proteins fusions composed of a dCre, a (GS_5_)_6_ linker and a nuclear anchor. The genes that are targeted and relocated to the nuclear periphery under 1 µM β-estradiol induction are repressed in transcription, while under the uninduced conditions, those genes maintain their original transcription. (B) Spot assays to examine the silencing effects of the synthetic tethers on a constitutively expressed β-carotene producing pathway. (C) Growth assays to test the silencing effects on a constitutively expressed synthetic *TRP* cluster in SC-Trp. OD_600_ of the cultures are determined by spectrophotometer after 22 hours of growing the strains overexpressing synthetic anchors with 1 µM β-estradiol induction, shown as the mean ± SD, n=3.

**Figure S11.**
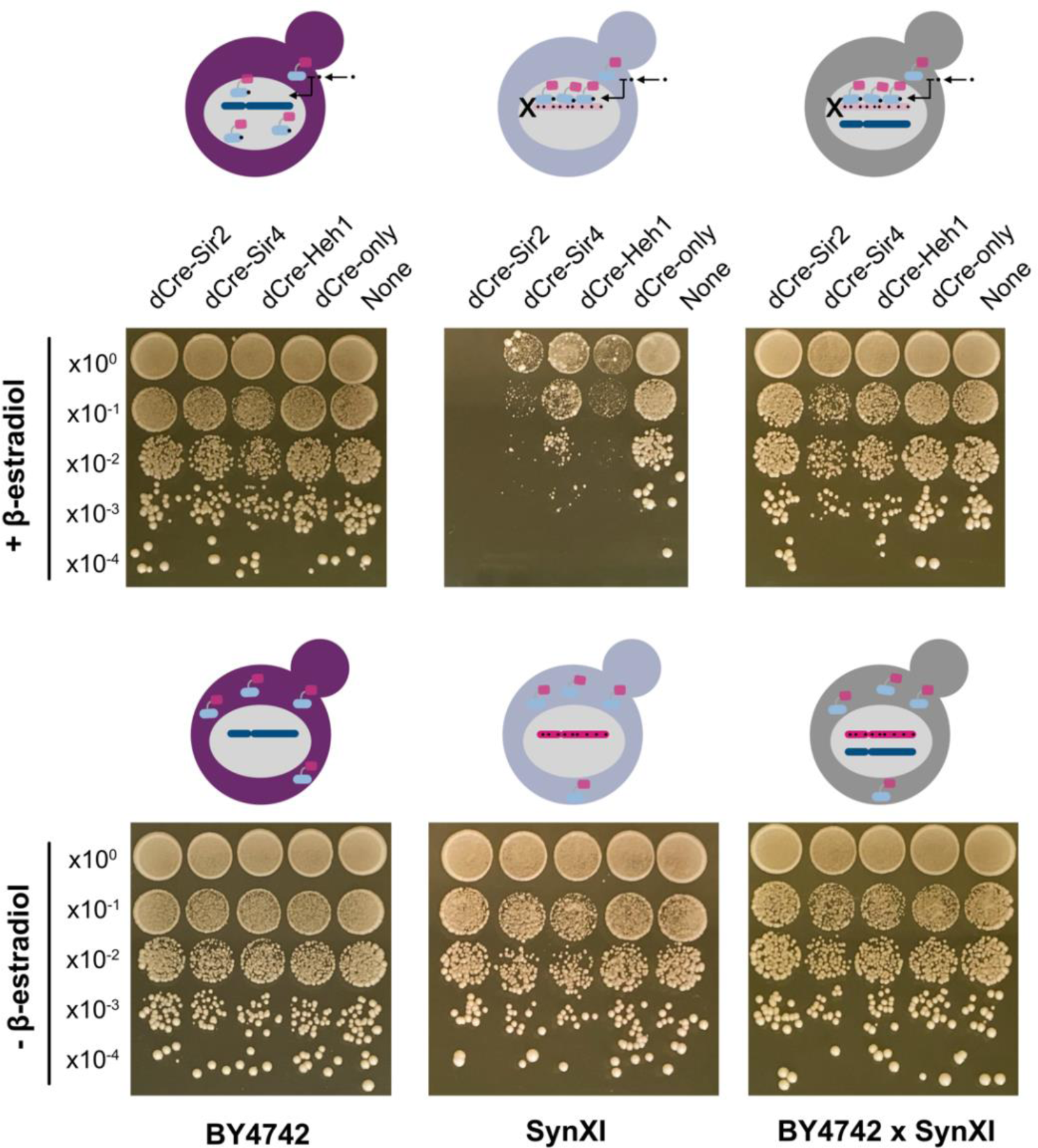
Spot assays to examine the silencing effects on *synXI*, related to Figure 5. Cultures of the haploid strain SynXI, the haploid control strain BY4742 and the heterologous diploid strain BY4742 x SynXI normalised to OD_600_ = 1.0 were serially diluted and spotted from top to bottom. Images were taken after 3 days of incubation on YPD plates with and without 1 µM β-estradiol.

**Figure S12.**
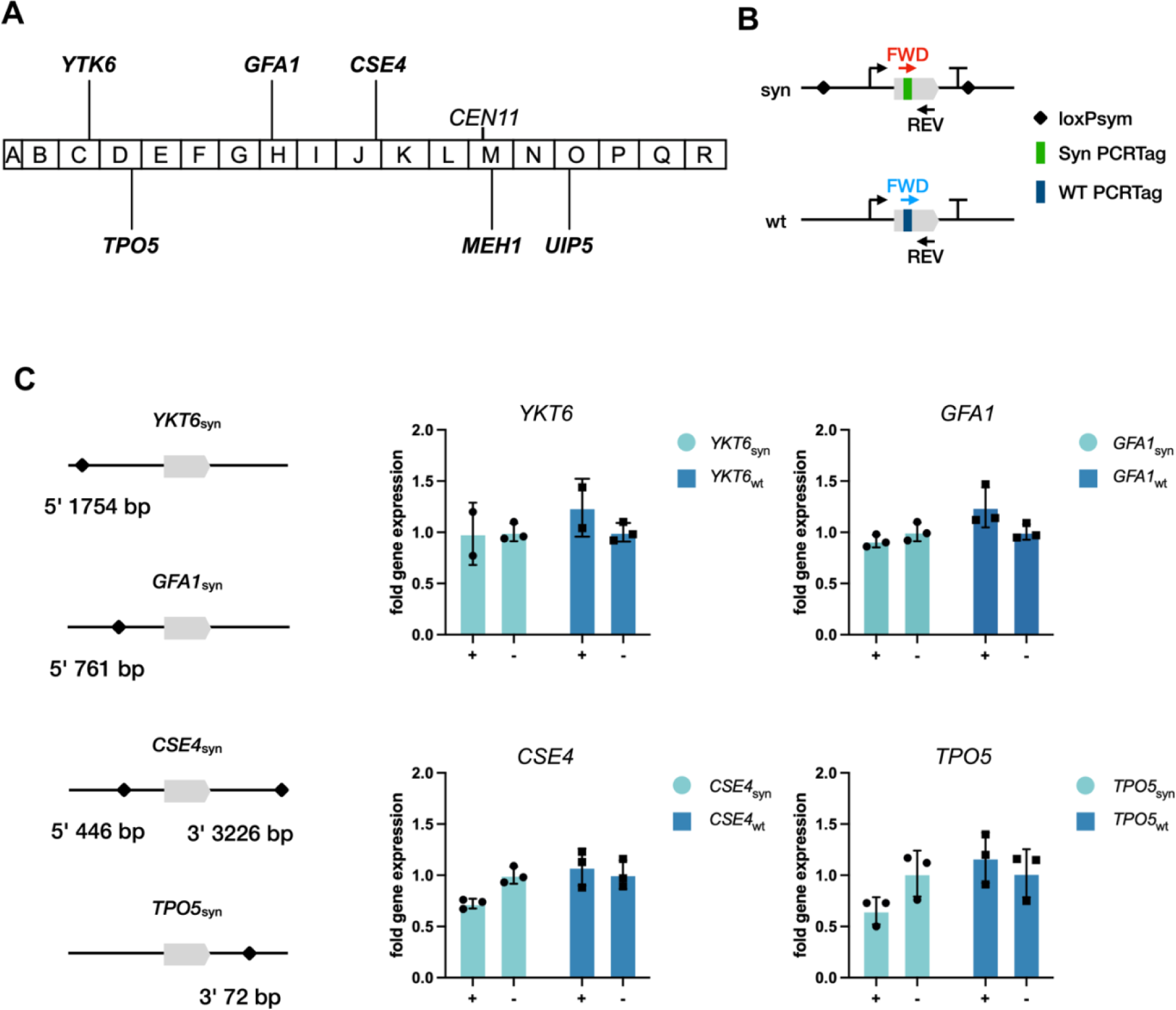
Quantification of the transcript levels of the synthetic and WT copies on chromosome XI by qPCR, related to Figure 5. (A) Schematic of the selected gens from various genomic positions on *synXI* for qPCR. Capitalised letters indicate the synthetised DNA chunks that are assembled into the complete *synXI*. (B) Schematic of the primer design for qPCR assay. The forward or reverse primer of qPCR primer pair is designed to target unique sequences within the synthetic PCRtags and the corresponding WT PCRtags. The other primer from the primer pair (corresponded reverse or forward primer) that targets both the synthetic and WT copies is same in sequence. (C) Left: Schematic showing the loxPsym insertion sites at each genomic locus of the tested genes. Right: Transcripts of the synthetic and WT copies of the tested genes under induced and uninduced conditions were quantified by qPCR using *ACT1* as a reference gene. Experiments were performed in biological triplicate. Individual data points of genes on *synXI* are plotted as round dots, individual data points of genes on WT *chrXI* are plotted as square dots, mean averages are denoted by bar height and error bars represent standard deviation.

**Figure S13.**
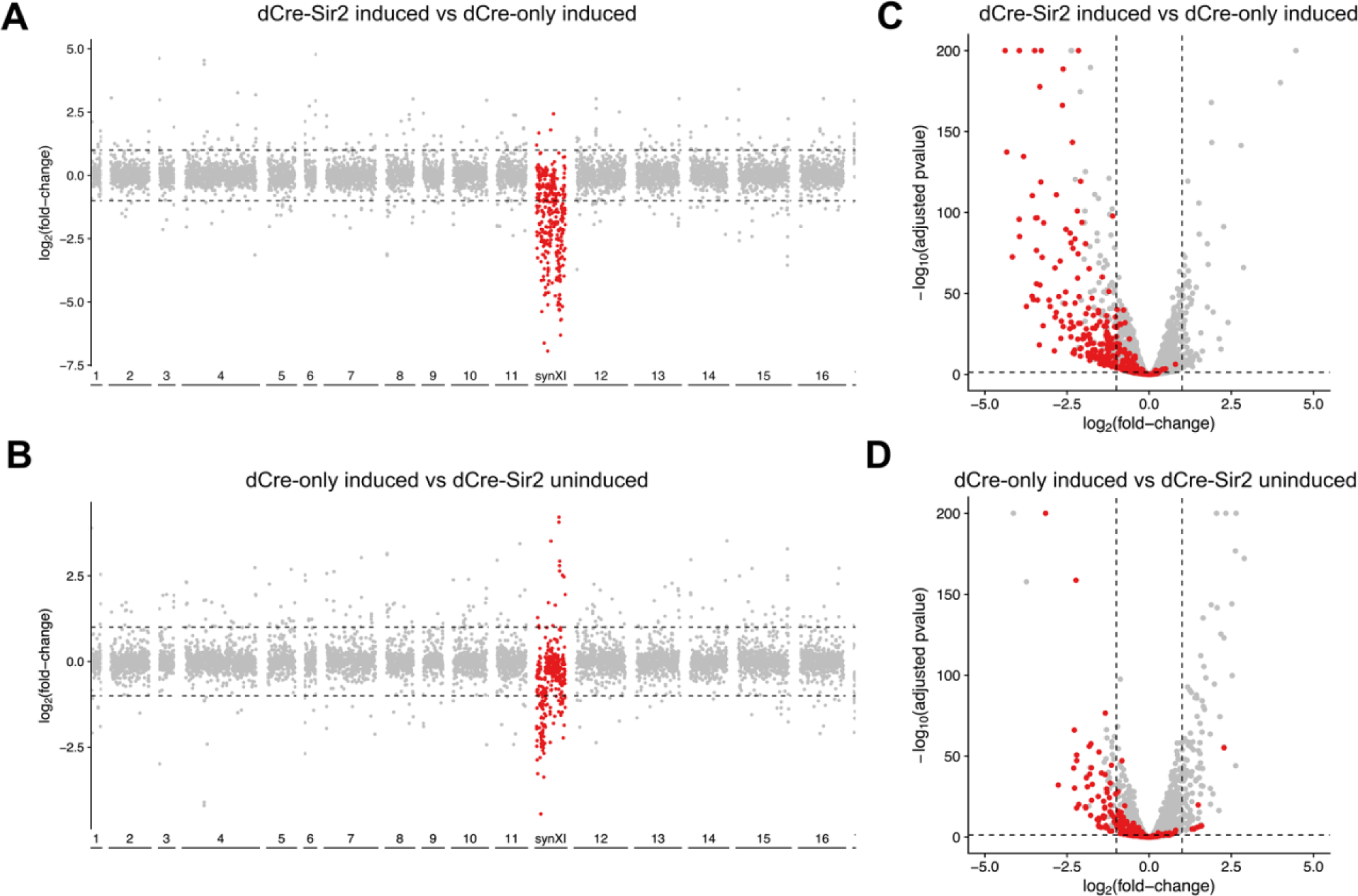
Differential gene expression on *synXI* and WT chromosomes in a heterologous diploid strain BY4742 x *synXI*, as determined by RNA-seq, related to Figure 5. (A and B) Manhattan plot of differential gene expression on *synXI* and WT chromosomes in heterologous diploid BY4742 x *synXI* overexpressing dCre-Sir2 fusion, as determined by RNA-seq. Comparisons were conducted between (A) dCre-Sir2 induced vs dCre-only induced, and (B) dCre-only induced vs dCre-Sir2 uninduced, respectively. Adjusted p-value cutoff was set at 0.05. Dashed line represents fold change threshold of 2 (log_2_ fold change = 1 or −1). Genes on *synXI* are shown as dots in red. Genes on WT chromosomes are shown as dots in grey with the numbers indicates each WT chromosome. (C and D) Volcano plot showing differential gene expression on *synXI* in heterologous diploid BY4742 x *synXI*, as determined by RNA-seq. Comparisons were conducted between (C) dCre-Sir2 induced vs dCre-only induced, and (D) dCre-only induced vs dCre-Sir2 uninduced, respectively. X-axis represents log_2_ fold change in gene expression between groups. Y-axis shows log_10_ of the p-value from the statistical test, with threshold of 0.05. Dashed line represents fold change threshold of 2 (log_2_ fold change = 1 or −1) and the p-value threshold of 0.05. Genes on *synXI* are shown as dots in red. Genes on WT chromosomes are shown as dots in grey.

**Figure S14.**
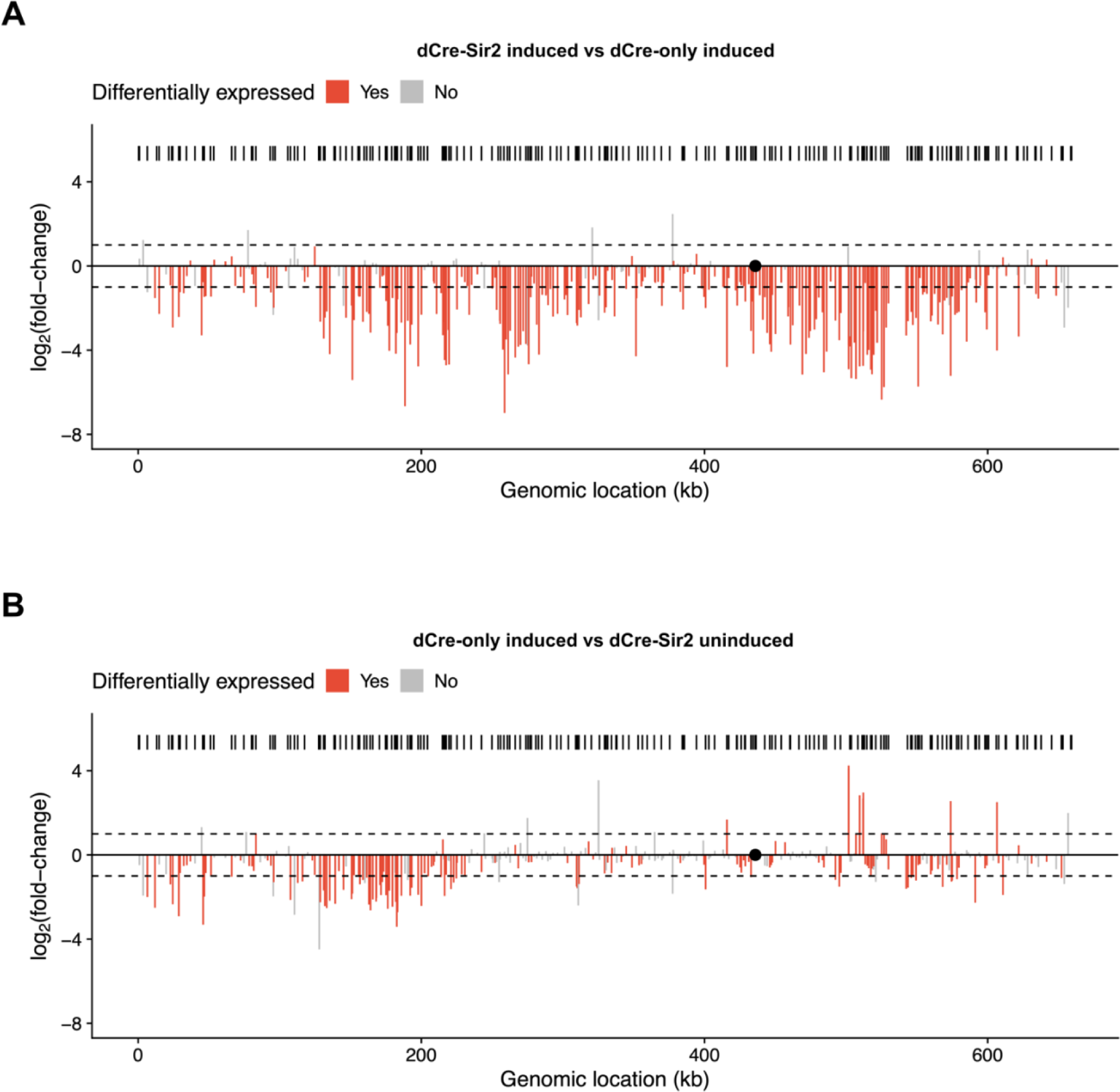
Bar plot showing differential gene expression across *synXI*, related to Figure 5. (A) Comparisons were conducted between samples dCre-Sir2 induced vs dCre-only induced. (B) Comparisons were conducted between samples dCre-only induced vs dCre-Sir2 uninduced. X-axis represents genomic location of genes. Y-axis represents log_2_ fold change. Locations of loxPsym sites are shown as the black barcode. Genes differentially expressed are marked in red. Genes not differentially expressed are grey. Dashed line represents fold change threshold of 2 (log_2_ fold change = 1 or −1). Centromere is labelled as a black dot.

**Figure S15.**
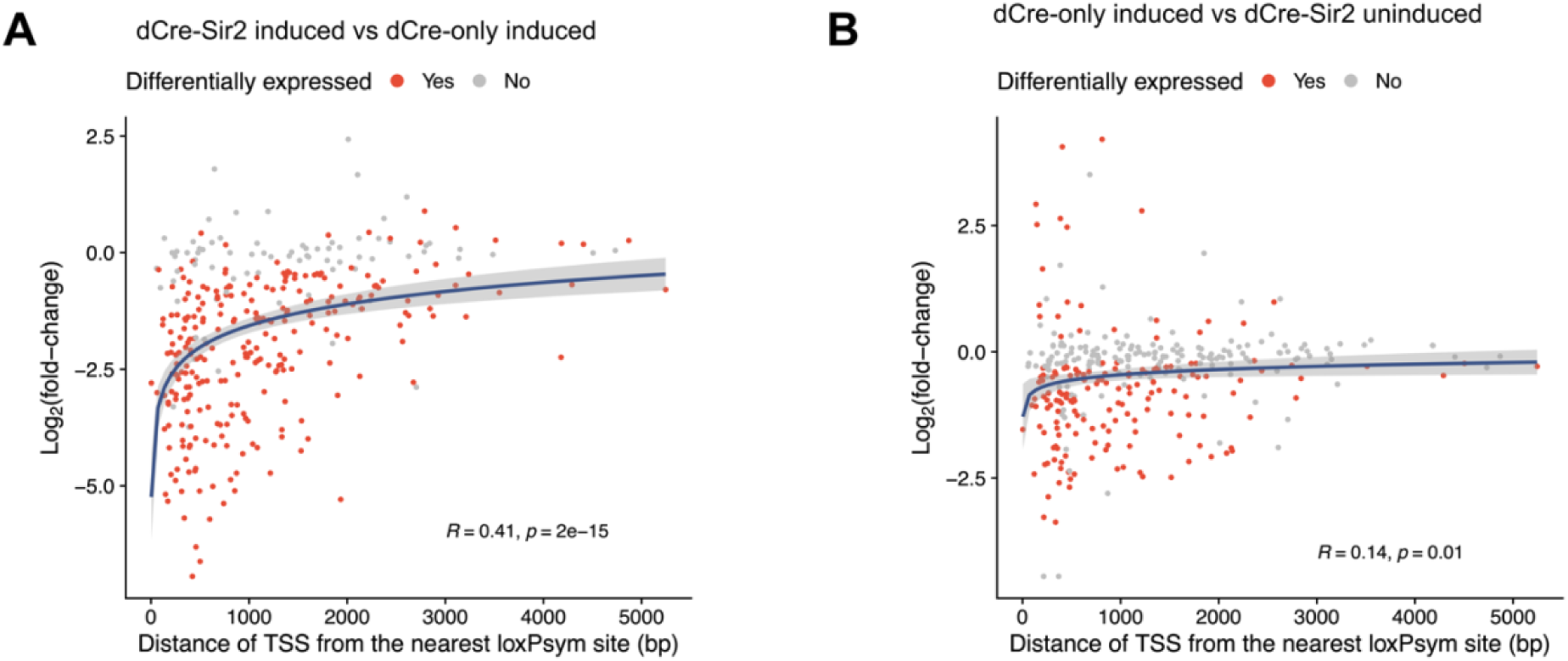
Correlation of the fold change expression of down-regulated genes on *synXI* with the distance of the nearest loxPsym sites to the gene start codon, related to Figure 5. Comparisons were conducted between (A) samples dCre-Sir2 induced vs dCre-only induced, and (B) dCre-only induced vs dCre-Sir2 uninduced, respectively. X-axis represents distance of nearest loxPsym site to the gene start codon. Y-axis represents the log_2_ fold change in gene expression. Genes differentially expressed are marked in red. Genes not differentially expressed are grey.

**Figure S16.**
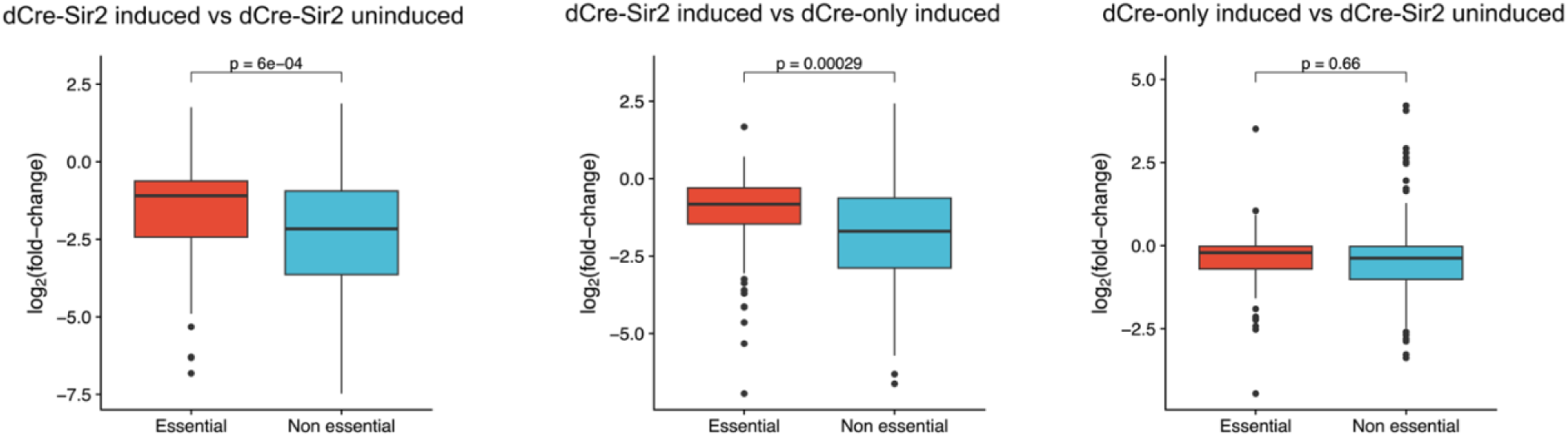
Correlation of the fold change expression of down-regulated genes on *synXI* with the essentiality of the genes. Left, comparison between samples dCre-Sir2 induced vs dCre-Sir2 uninduced. Middle, comparison between samples dCre-Sir2 induced vs dCre-only induced. Right, comparison between samples dCre-only induced vs dCre-Sir2 uninduced. X-axis represents the essential and non-essential genes. Y-axis represents the log_2_ fold change in gene expression of two groups of genes. Red represents essential genes and blue represents non-essential genes.

## Supplementary Tables

**Table S1.**
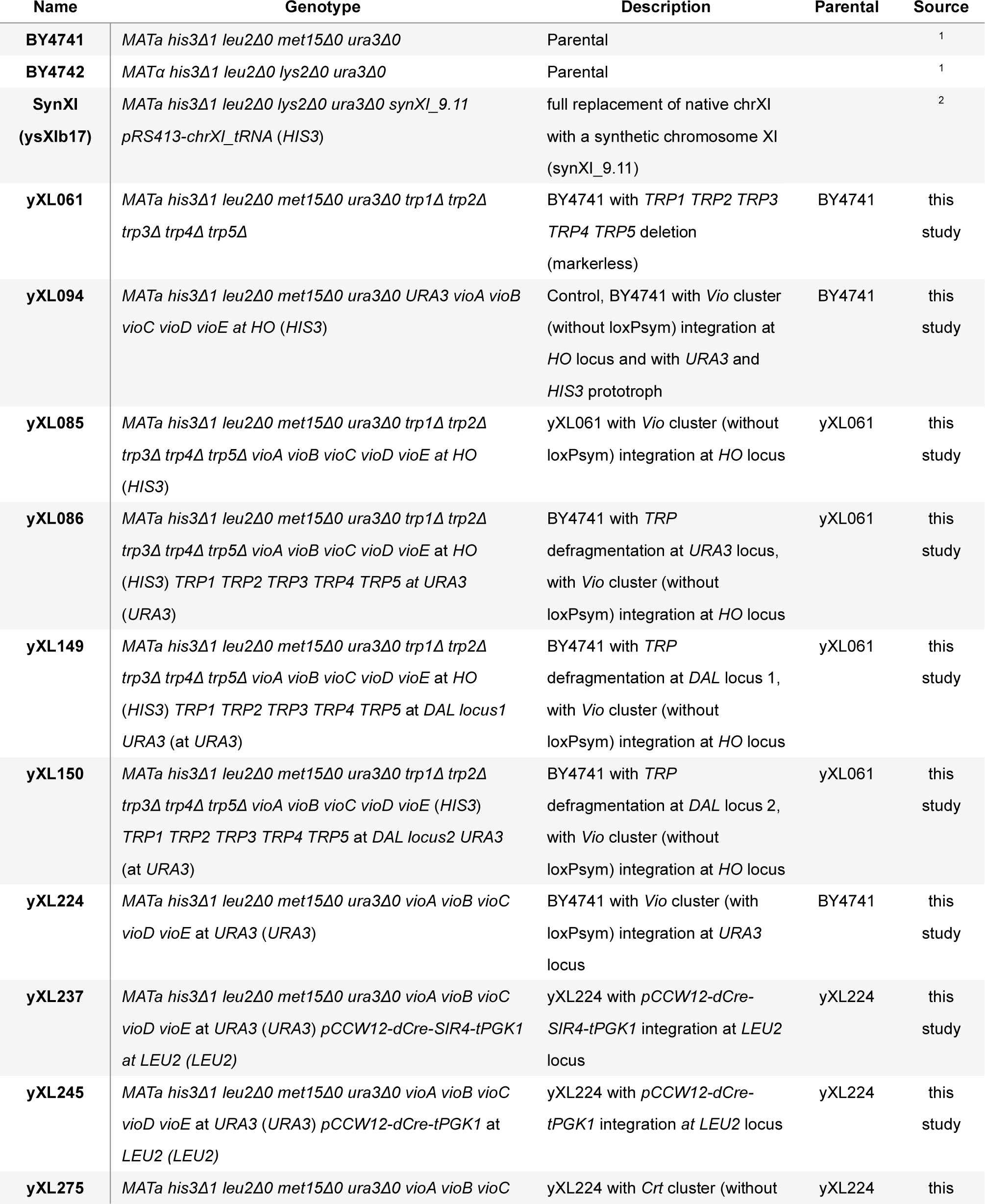
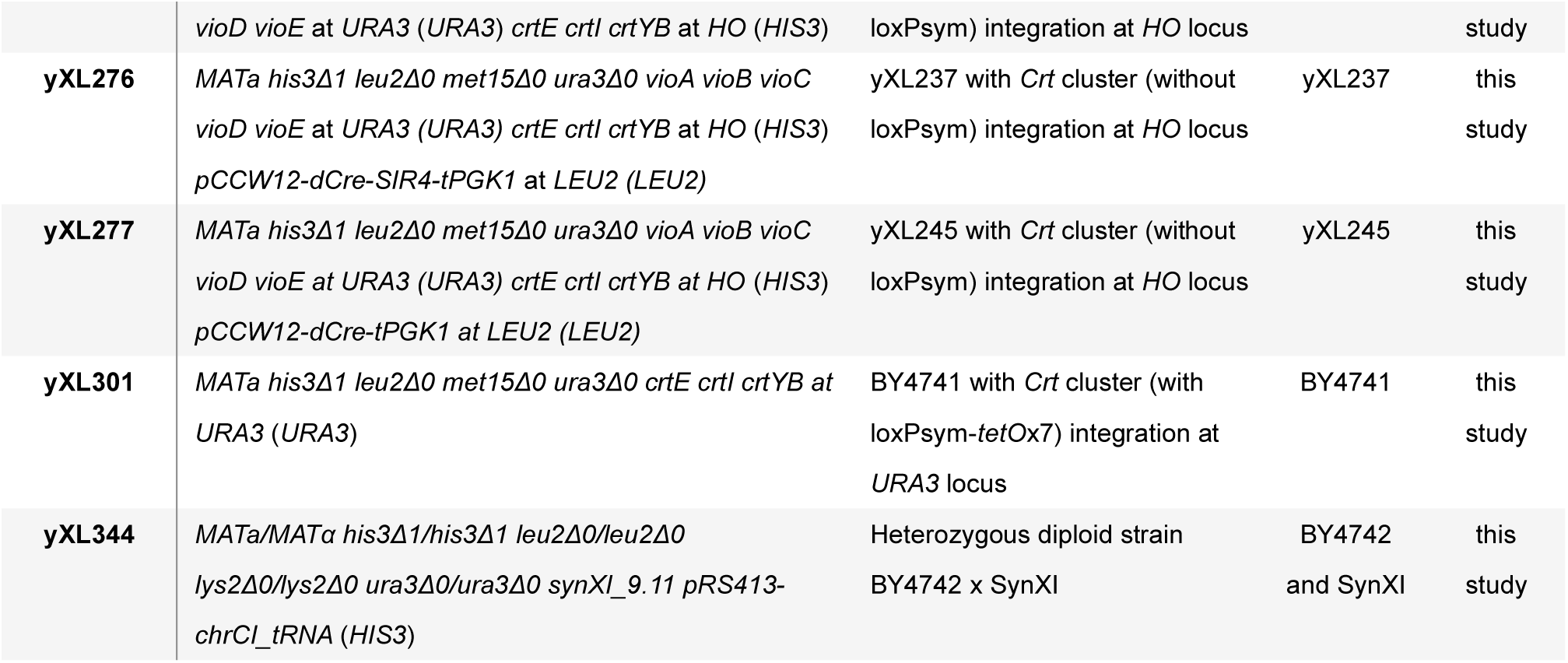
List of yeast strains used in this study.

**Table S2.**
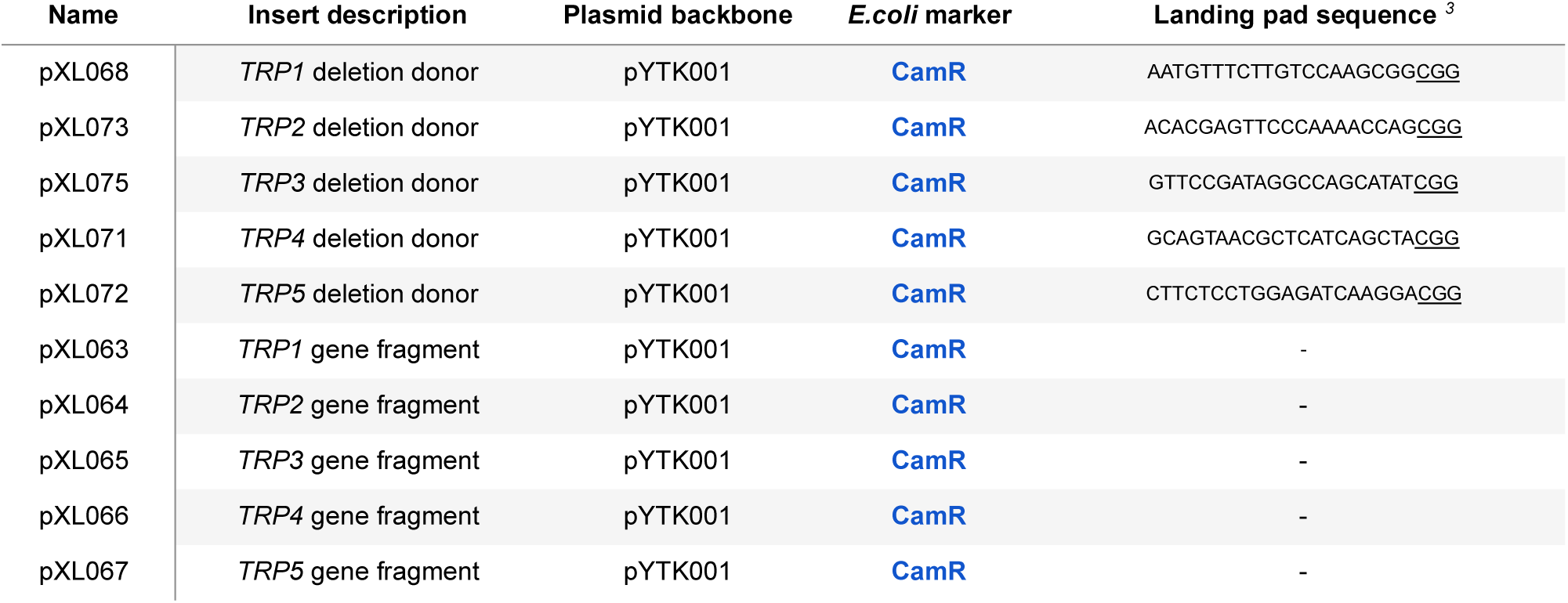
List of gap repair donor and gene fragment plasmids used for defragmentation of *TRP* biosynthesis. To ease future engineering of the sites left behind by gene deletion, we substituted each deleted sequences with an individual 23 bp ‘landing pad’^3^ that encodes a unique CRISPR/Cas9 target sequence. Landing pad sequences are shown in the right column with protospacer adjacent motif (PAM) sequence highlighted by underlining.

**Table S3.**
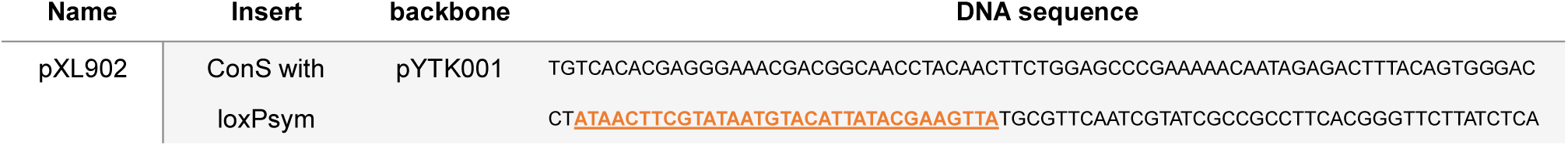
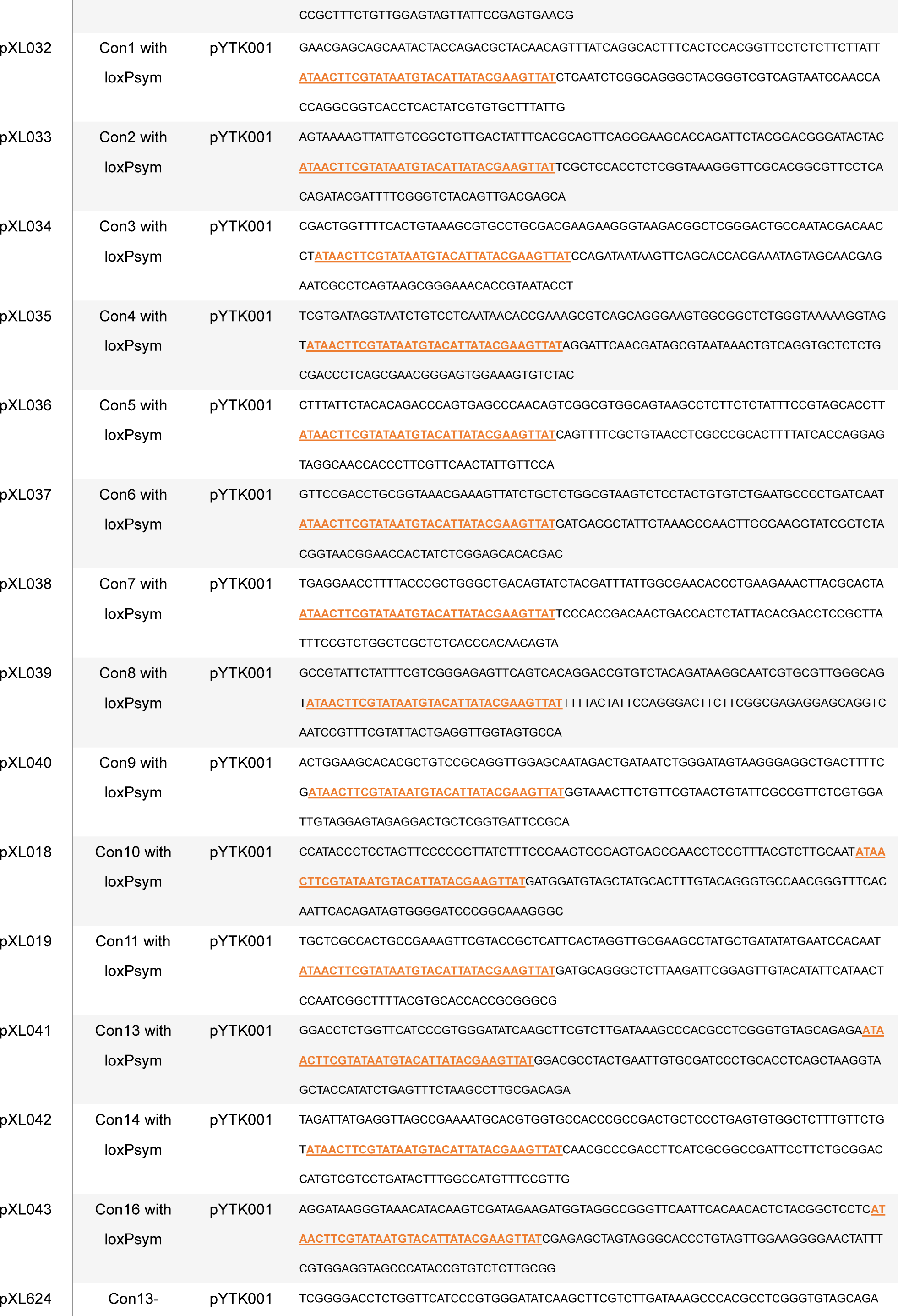
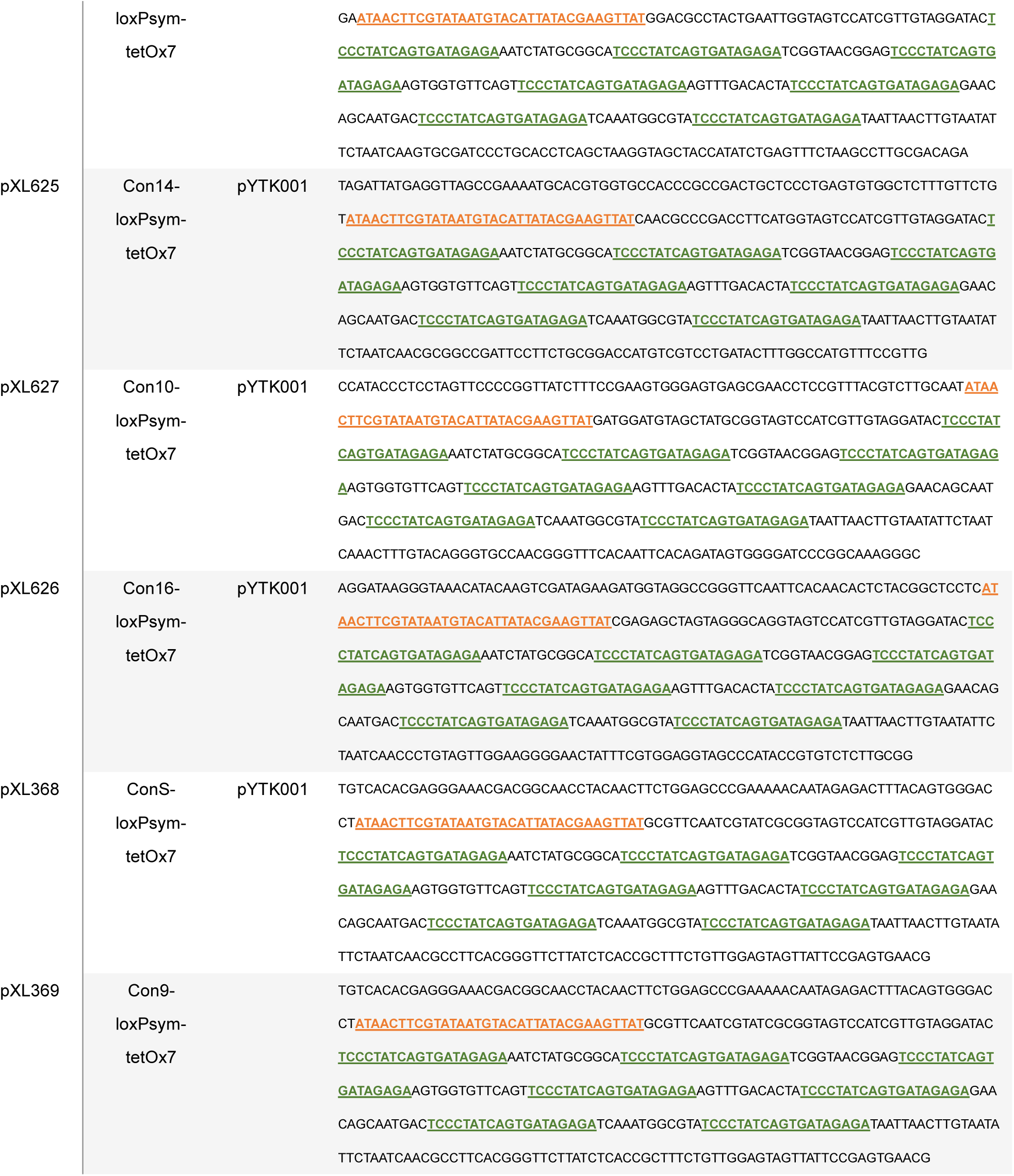
List of linker plasmids used in this study. Sequence of loxPsym is highlighted in **<u>orange</u>**. Sequence of tetO is highlighted in **<u>green</u>**.

**Table S4.**
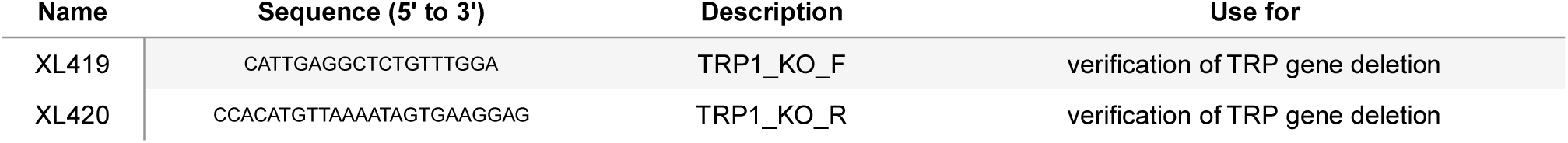
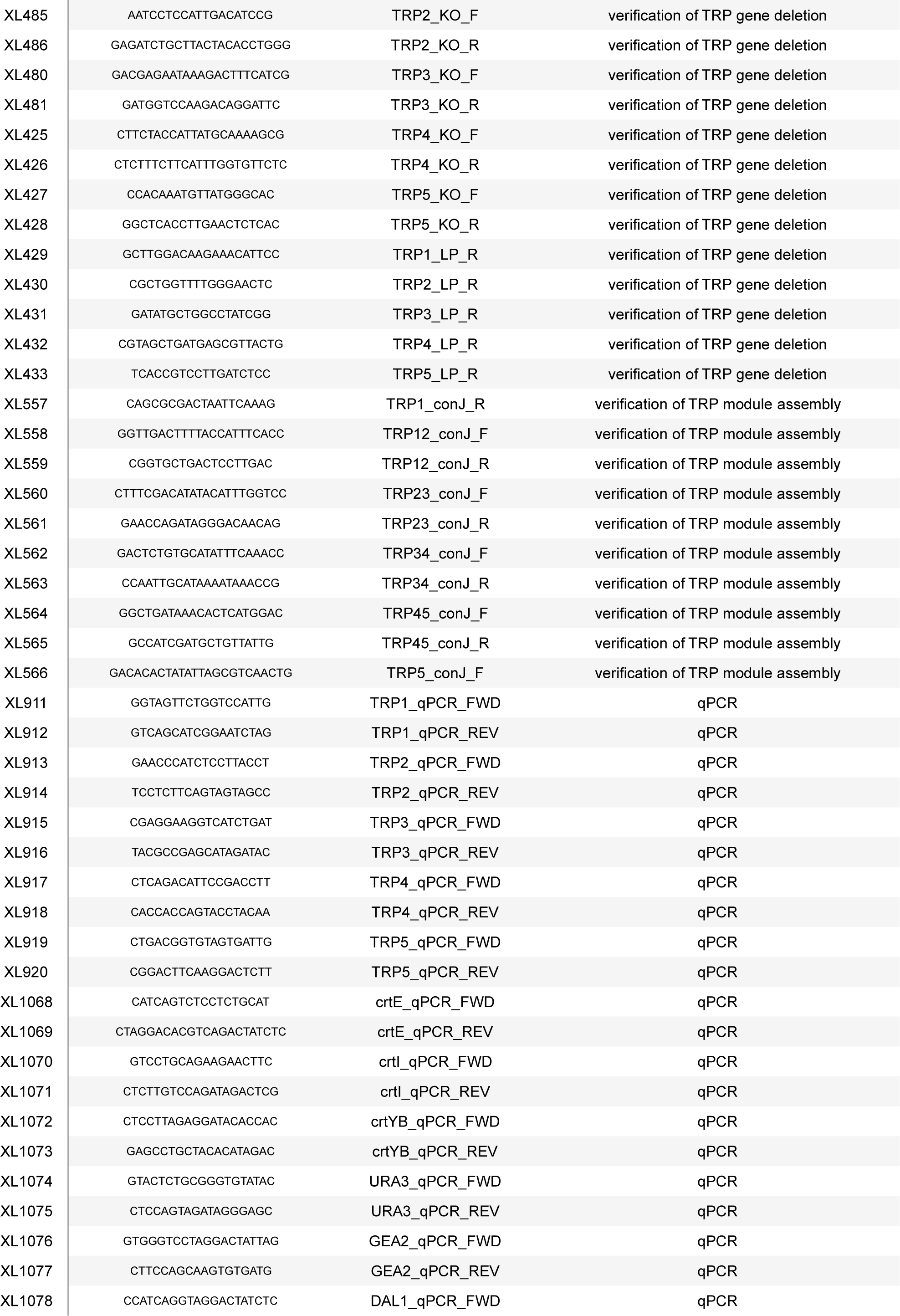
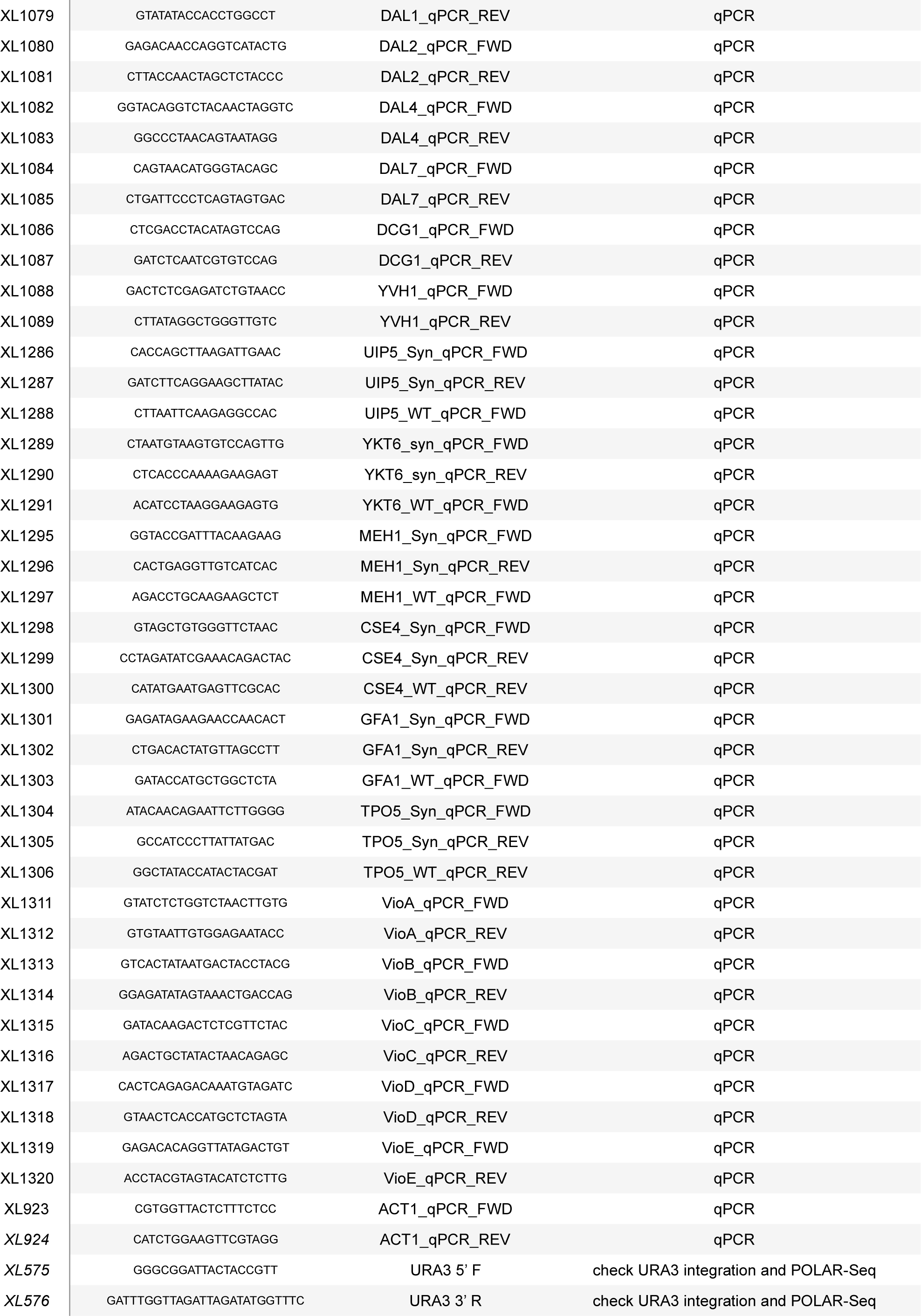
List of primers used in this study.

**Table S5.**
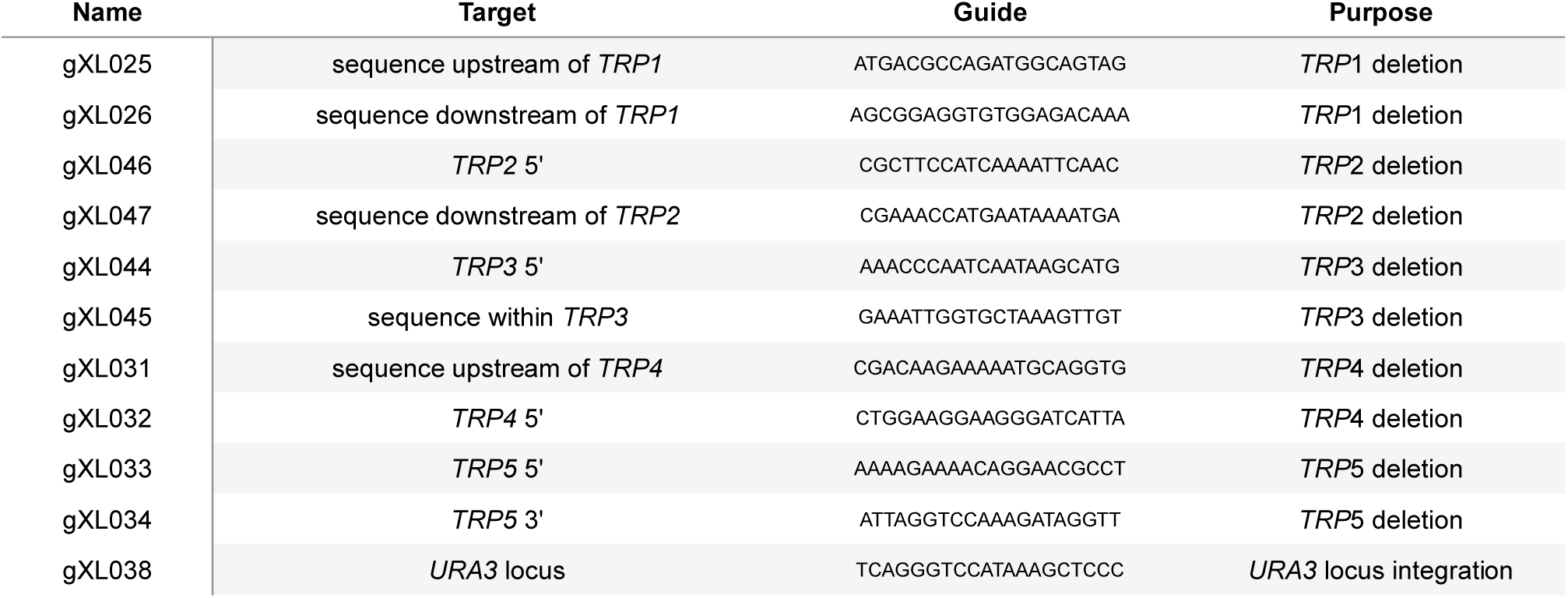
List of gRNAs used in this study.

**Table S6.**
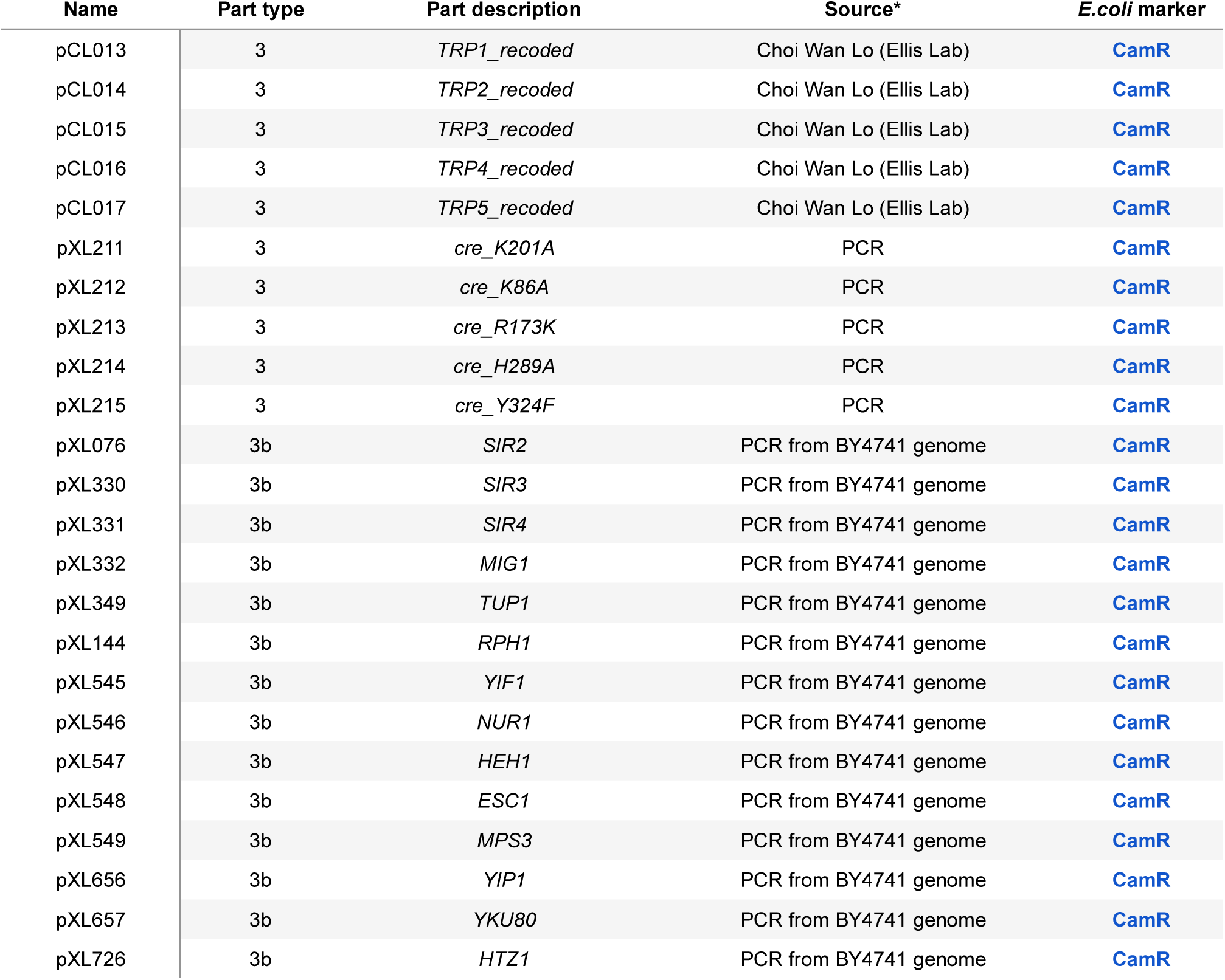
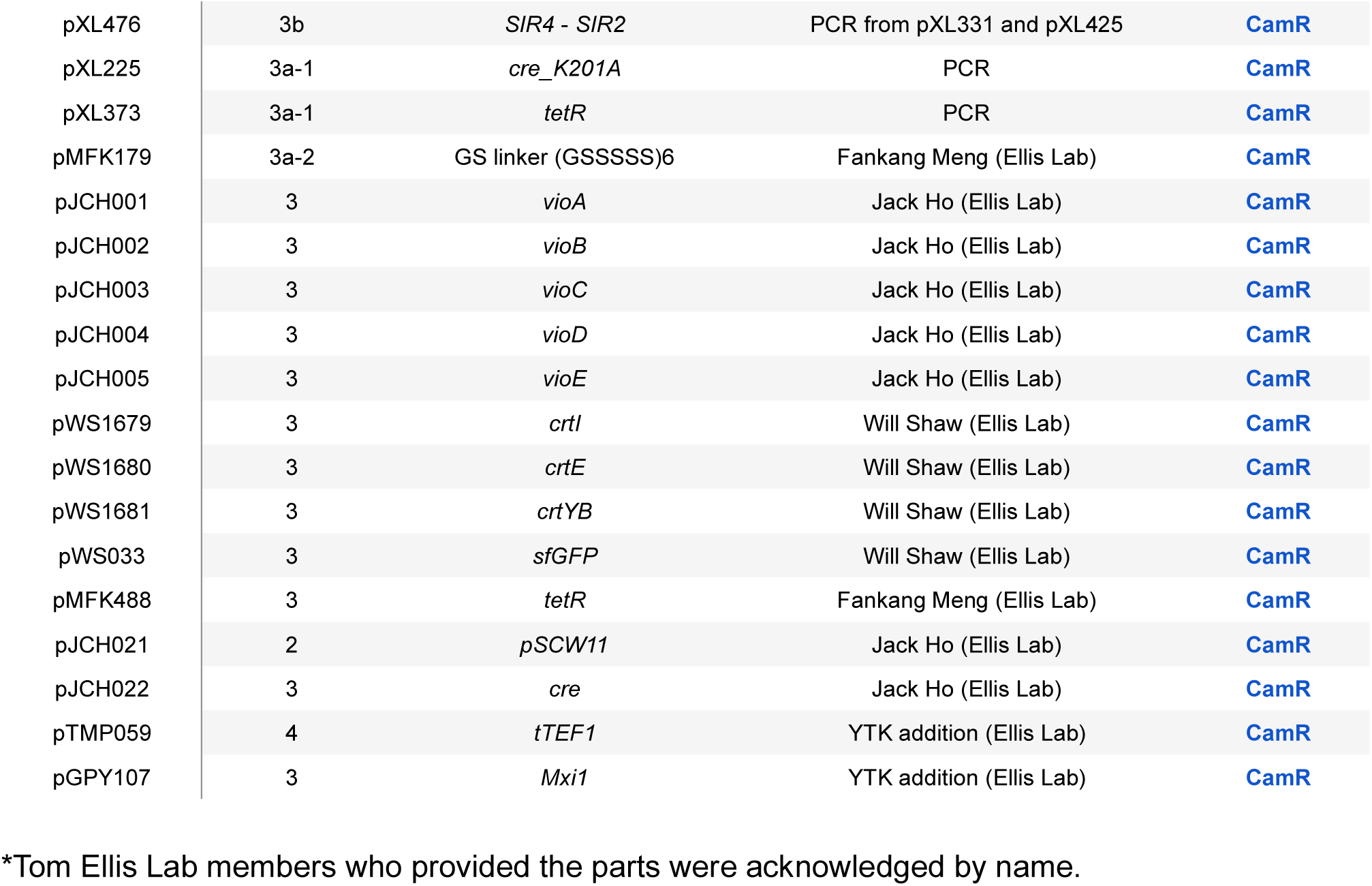
List of parts in YTK format used in this study.

**Table S7.**
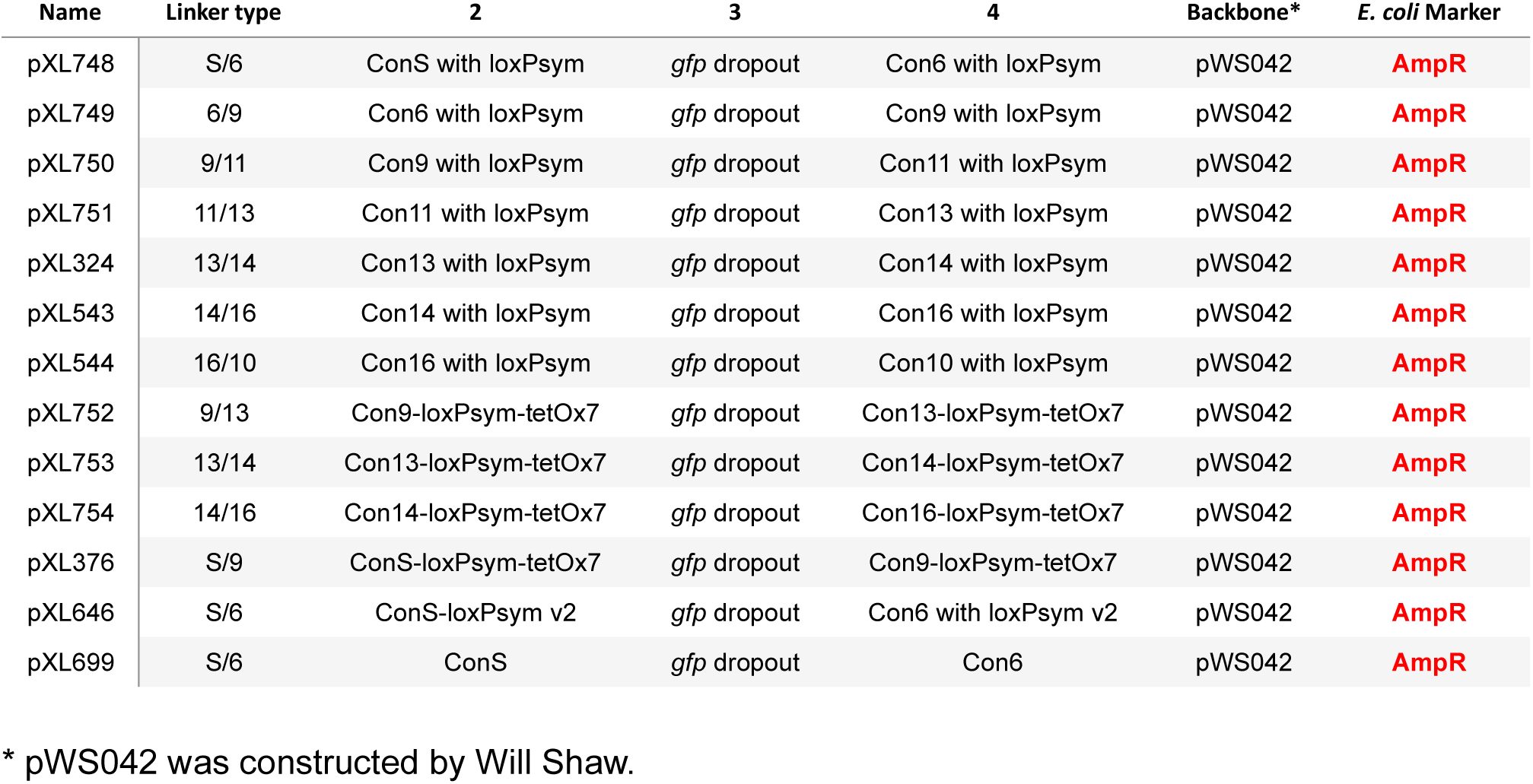
List of pre-assembled linker vectors used in this study.

**Table S8.**
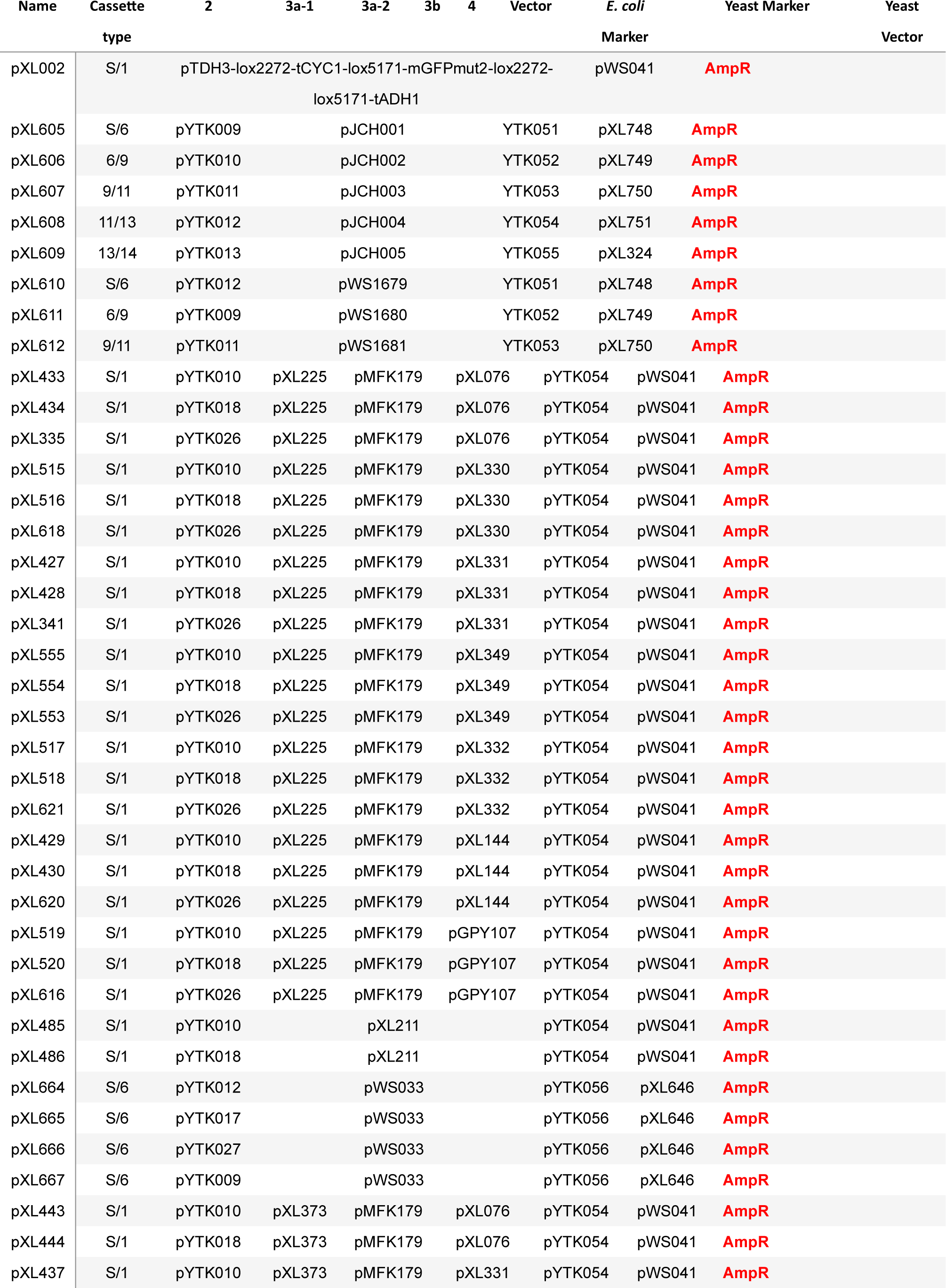
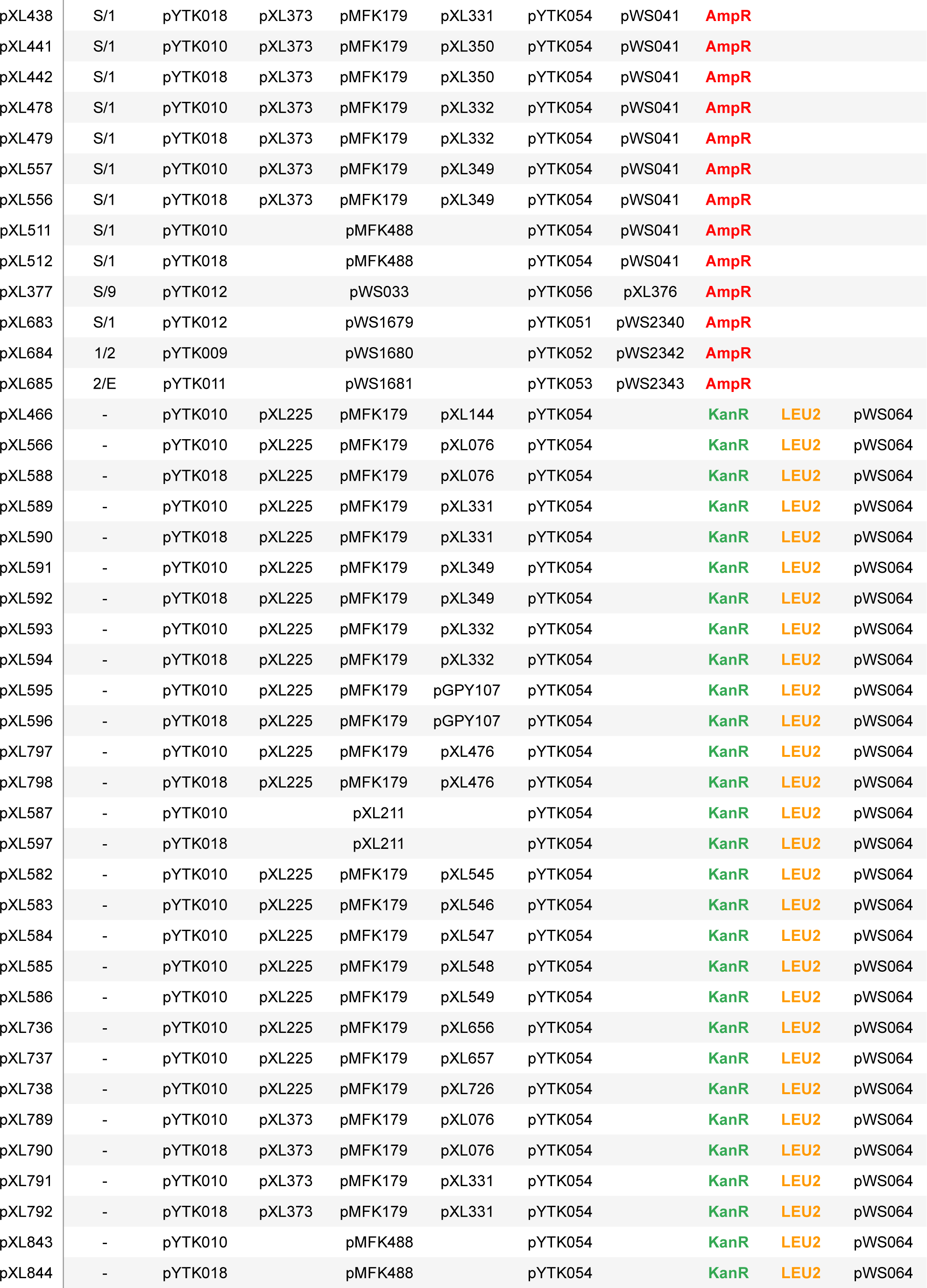
List of cassettes used in this study.

**Table S9.**
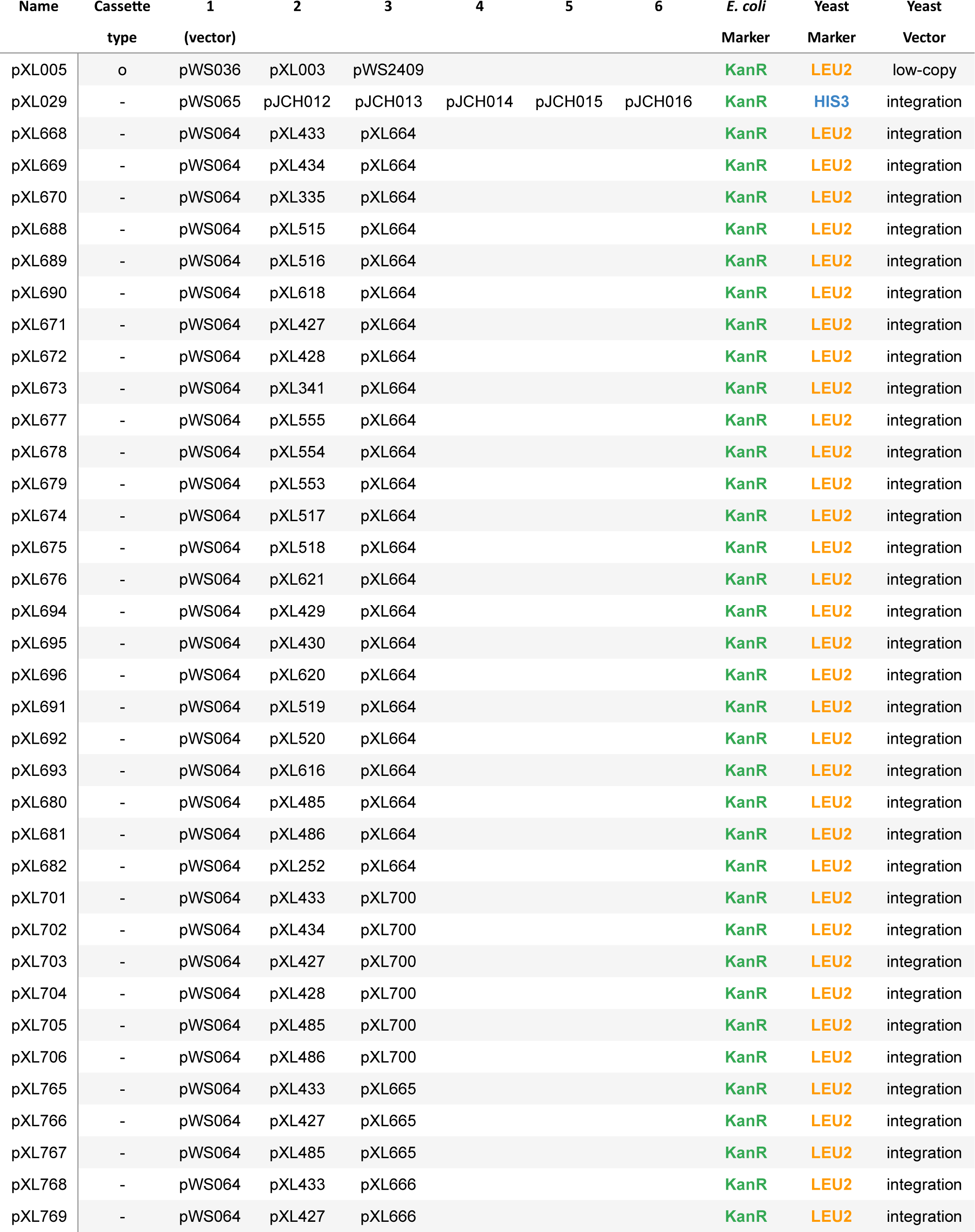
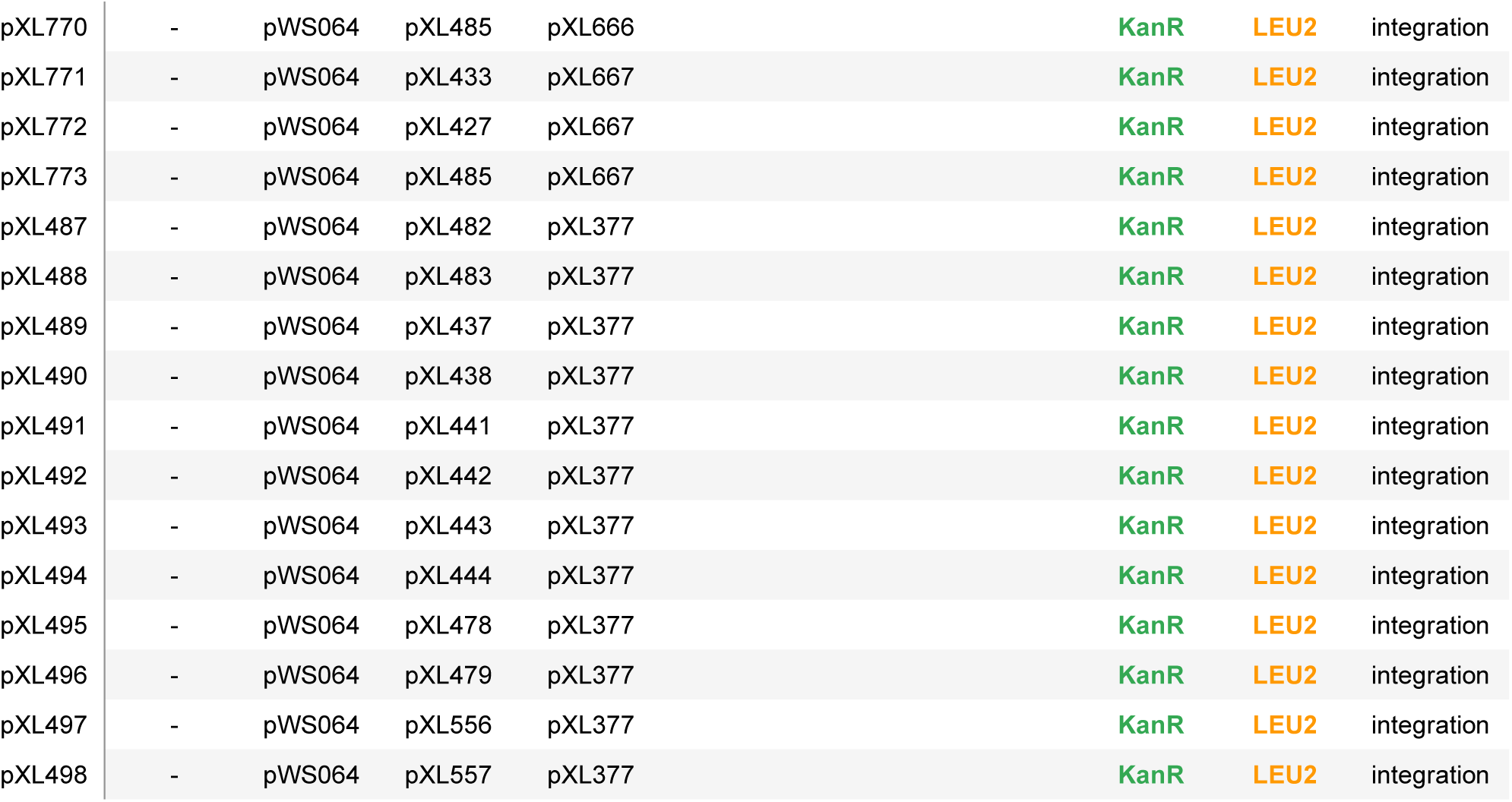
List of multigene cassettes used in this study. Plasmids named under ‘pWS’ were constructed by Will Shaw. Plasmids named under ‘pJCH’ were constructed by Jack Ho.

**Table S10. Differential gene expression analysis of the RNA-Seq data.**

Table can be found in “Table S10.xls”.

